# Identifying and targeting the Mg-Fe-PmrAB regulatory circuit reverses phosphate starvation-driven polymyxin tolerance in *Enterobacteriaceae*

**DOI:** 10.1101/2025.08.18.670855

**Authors:** Guangming Zhang, Ziqing Deng, Jiezhang Jiang, Minji Wang, Xiaoyuan Wang, Xiaoyun Liu, Aixin Yan

## Abstract

Phosphate (Pi) scarcity, a pervasive stressor prevalent in bacterial infections and dysbiotic host environments, potently alters antimicrobial susceptibility against cationic antibiotics like polymyxins across diverse bacterial taxa. While established Pi depletion-induced antimicrobial tolerance mechanisms often involve membrane phospholipid remodelling, *Enterobacteriaceae* largely lack these pathways, leaving the mechanism underlying their adaptation to Pi scarcity unresolved. Here, we reveal that Pi limitation robustly induces polymyxin tolerance in *Enterobacteriaceae* through 4-amino-4-deoxy-L-arabinose (L-Ara4N) modification of the Gram-negative outer membrane component lipid A, reflecting a distinct adaptive strategy. This modification is driven by strong *arn* operon induction, mediated by the PmrAB two-component system. We discover that Pi depletion triggers cellular Mg²⁺ release, prompting compensatory Fe³⁺ mobilization to the cell envelope that directly activates PmrAB. Crucially, this metal-dependent signaling axis offers a directly targetable mechanism for reversing polymyxin tolerance, a significant deviation from the less accessible, PhoBR-regulated phospholipid remodelling strategies in other bacteria. We demonstrate that Mg²⁺ supplementation or Fe³⁺ chelation effectively suppresses PmrAB activation and restores polymyxin susceptibility, thereby establishing a novel, metal-centric paradigm for understanding and pharmacological intervention in stress-induced antimicrobial tolerance.

## Introduction

Within the dynamic and often hostile environment of the host, nutrient scarcity presents a pervasive challenge for invading bacteria, promoting the evolution of survival strategies and shaping pathogen persistence.^1,2^ Among the macronutrients essential for bacteria, phosphorus (P) serves as a critical element for life. It is a fundamental component of key biomolecules such as nucleic acids, phospholipids, and ATP, and plays crucial roles in the essential processes of genetic inheritance, energy metabolism, membrane integrity, and signal transduction.^3,4^ The primary assimilable form of phosphorus for bacteria is the orthophosphate anion (PO_4_^3-^), commonly known as inorganic phosphate (Pi). However, Pi is typically found at very low concentrations (<4μΜ) in diverse environments, from oceans (0.1-3μΜ),^5,6^ and soil (0.0105 to 10.5μΜ, <1μΜ near plant roots),^7,8^ to critically host-associated niches including macrophages,^9^ Salmonella-containing vacuoles,^10^ and cystic fibrosis sputum.^11^ This localized Pi scarcity is a direct consequence of host nutritional immunity and intense microbial competition.^12,13^ Evidence for widespread bacterial adaptation to Pi limitation is abundant, with metagenomic and metatranscriptomic studies consistently revealing robust transcriptional upregulation of high-affinity phosphate acquisition (Pho) regulon genes across various chronic infections and dysbiosis conditions.^11,14,15^ Bacteria cope with these pervasive conditions via the conserved PhoBR two-component system, which activates genes for Pi acquisition, transport, and storage when extracellular Pi falls below 4 μM.^3,16,17^

Beyond direct Pi assimilation, Pi depletion triggers broader adaptive responses crucial for pathogen survival, including enhanced virulence and, critically, reduced antimicrobial susceptibility.^18–20^ For example, Pi limitation upregulates virulence factors in *Vibrio cholerae*, EHEC, and *Pseudomonas aeruginosa*, and conversely, Pi supplementation can reduce infection severity.^21–24^ More importantly, Pi depletion is increasingly recognized as a potent environmental cue altering antimicrobial susceptibility across diverse bacterial taxa, particularly against cationic antibiotics like polymyxins. This is evidenced by increased vancomycin resistance in *Streptomyces coelicolor*,^25^ and enhanced polymyxin B/E tolerance in *Pseudomonas fluorescens*, *P. aeruginosa*, and *Acinetobacter baumannii* under Pi-limited conditions.^11,26–28^ Given that Pi limitation is a prevalent and dynamic stressor in host niches, understanding its role in promoting antimicrobial tolerance is vital for developing effective counter-strategies.

Mechanistically, Pi depletion-induced antimicrobial tolerance is often linked to the remodelling of bacterial membranes, where phosphorus-containing phospholipids are substituted with diverse non-phosphorous lipids. This strategy spares intracellular phosphate and often modifies membrane charge, which is critical for tolerance to cationic antimicrobials. Examples include the accumulation of ornithine lipids (OLs) in *Vibrio cholerae,*^29^ diacylglyceryl-N,N,N-trimethylhomoserine (DGTS) in *Rhizobium meliloti*,^30^ glycolipids in *Agrobacterium tumefaciens*,^31^ and extensive lipidome remodelling in *Mycobacterium tuberculosis*.^32^ Specifically for polymyxin tolerance, *P. aeruginosa* synthesizes neutral monoglucosyl diacylglycerol (MGDG) and glucuronic acid diacylglycerol (GADG),^11^ while *A. baumannii* produces positively charged ornithine- and lysine-containing aminolipids (OL and LL) under Pi limitation.^27^ These diverse lipid modifications effectively reduce the net negative charge of the bacterial membrane, decreasing polymyxin binding and improving tolerance.

However, these established lipid-remodelling strategies are not universally conserved. Critically, genes for OL biosynthesis are absent in most *Enterobacteriaceae*, a family of paramount clinical importance, and MGDG/GADG synthesis is not a predominant mechanism for polymyxin tolerance in this group.^33–35^ Given the widespread prevalence of Pi limitation in host environments and the escalating threat of polymyxin tolerance in *Enterobacteriaceae*, a fundamental question with significant clinical implications arises: what alternative, yet unidentified, mechanisms underpin Pi depletion-induced polymyxin tolerance in this critical bacterial family, especially in the absence of the established OL and MGDG/GADG pathways?

In this study, we address this critical gap by reporting that Pi limitation also induces polymyxin tolerance in *Enterobacteriaceae*, but through a distinct mechanism. Using *E. coli* K-12 MG1655 as a model, we demonstrate that this process involves the upregulation of conserved Ugd and ArnBCADTEF proteins, which catalyse the addition of the positively charged 4-amino-4-deoxy-L-arabinose (L-Ara4N) to lipid A. This adaptive lipid A modification is driven by a novel Pi-Mg-Fe-PmrAB regulatory circuit: loss of membrane-stabilizing magnesium (Mg^2+^) and compensatory iron (Fe^3+^) mobilization to the cell membrane activate the PmrAB two-component system, leading to upregulation of the *ugdarn* operon. Crucially, interfering with this circuit, via Mg^2+^ supplementation or iron chelation, effectively reverses tolerance in representative *Enterobacteriaceae* species, but not in non-enteric species like *P. aeruginosa*. Phylogenetic analyses further reveal distinct clades for *E. coli* PmrB (EcoPmrB) and *P. aeruginosa* PmrB (PaePmrB), with PaePmrB lacking the conserved “ExxE” motif required for Fe^3+^ sensing. Our findings uncover a conserved, targetable stress-responsive circuit underlying Pi depletion-induced polymyxin tolerance within *Enterobacteriaceae*, offering new avenues for managing stress-induced antibiotic tolerance.

## Results

### Pi depletion induces polymyxin tolerance in *Enterobacteriaceae*

To investigate whether Pi limitation induces antibiotic tolerance in *Enterobacteriaceae*, we assessed the susceptibility of two representative species, *E. coli* MG1655 and *Salmonella enterica* serovar Typhimurium ATCC14028. Bacteria were initially cultured in standard MOPS synthetic minimal medium containing replete Pi (Pi+,1.32 mM) for 5 hours. Subsequently, cultures were transferred to acute Pi-depleted (Pi-, 0 mM) conditions for 3 hours, a protocol adapted from Groat et. al.^36^ Following this, bacterial survival was determined via 3-hour polymyxin B killing assays (Figure S1). Polymyxin B concentrations for these assays were established based on the minimum inhibitory concentrations (MICs) of strains in MOPS minimal medium (Table S1) and a preliminary dose-dependent killing assay (Figure S2). Results showed that Pi depletion enhanced the survival of both species against polymyxin B (Figure 1A). Similar Pi depletion-induced polymyxin tolerance was also observed in *E. coli* O157 and four uropathogenic *E. coli* strains isolated from urinary tract infection patients in the Queen Mary Hospital, Hong Kong (Figure 1B). Consistent with previous reports, *P. aeruginosa* PAO1 subjected to Pi depletion also display enhanced tolerance against polymyxin B (Figure 1B). These results suggest that Pi depletion-induced polymyxin tolerance is a widespread phenomenon among Gram-negative bacteria.

**Figure 1.**
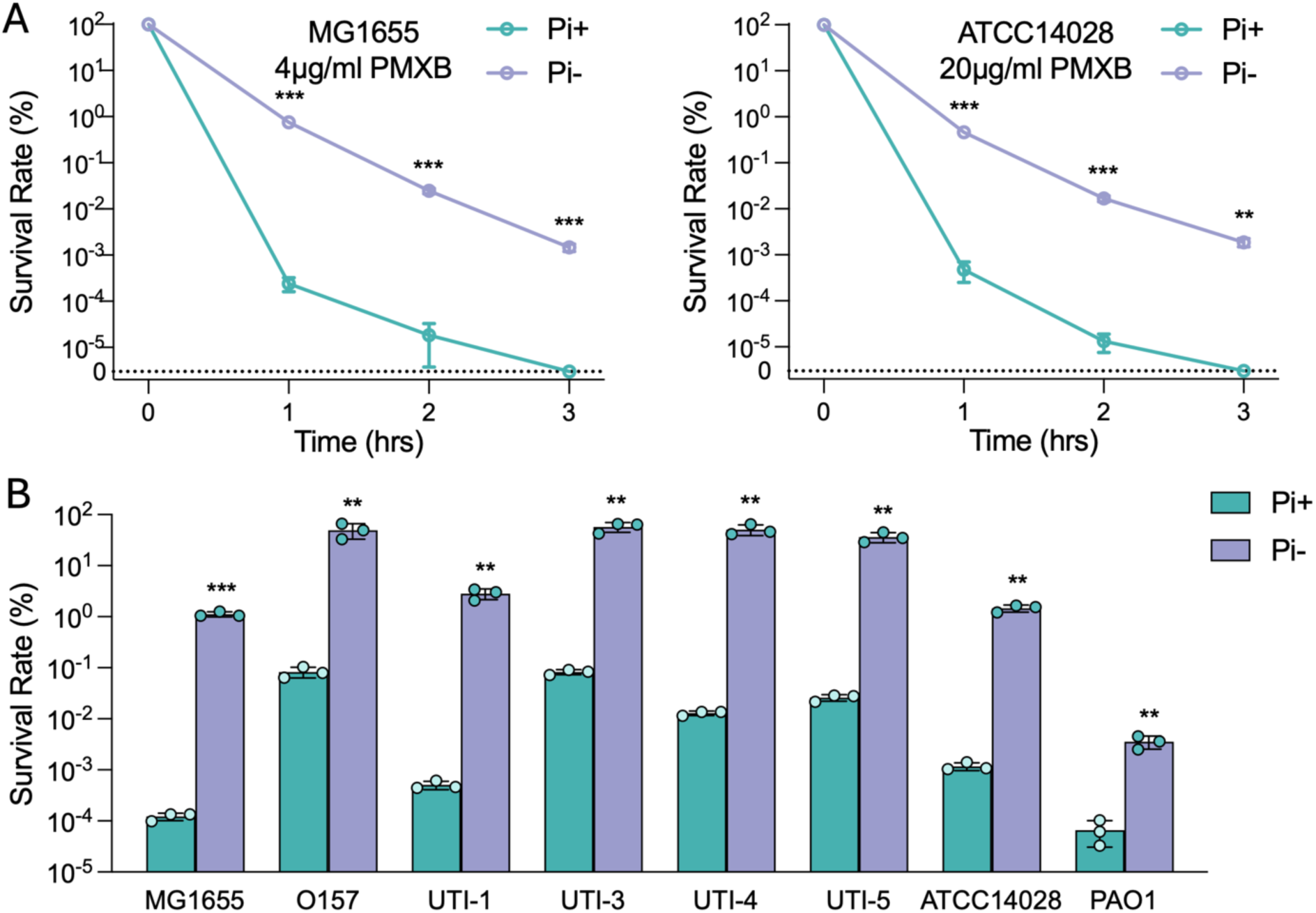
Pi depletion induces polymyxin tolerance in diverse bacterial species. (A) Time-kill curves of *Escherichia coli* MG1655 (left panel) and *Salmonella*. Typhimurium ATCC14028 (right panel) exposed to polymyxin B (PMXB) (4 μg/ml for *E. coli* strains, 20 μg/ml for *S.* Typhimurium) following acute Pi-starvation treatment (Pi-, 0 mM Pi) compared to the group without treatment (Pi+, 1.32 mM Pi). The dotted line labelled “0” denotes the limit of detection; values below this threshold were not detectable by colony counting. (B) Survival rates for various *Enterobacteriaceae* strains (*E. coli* MG1655, O157, UTI-1, UTI-3, UTI-4, UTI-5, *S.* Typhimurium ATCC14028) and *Pseudomonas aeruginosa* PAO1 exposed 1-hour to polymyxin B (4 μg/ml for *E. coli* strains, 20 μg/ml for *S.* Typhimurium and 8 μg/ml for *P. aeruginosa*). For all tested strains, phosphate depletion (Pi-, purple bars) significantly increases polymyxin B tolerance compared to phosphate-replete (Pi+, teal bars) conditions. Error bars represent the standard deviation of at least three independent experiments. Statistical significance was determined using unpaired two-tailed Student’s t-test. Significance levels are indicated as follows: \**p* < 0.05; \*\**p* < 0.01; \*\*\**p* < 0.001.

### Pi depletion induces 4-amino-4-deoxy-L-arabinose (L-Ara-4N) modification of lipid A in *Enterobacteriaceae*

Since these species lack the MGDG/GADG and OL pathways described in *P. aeruginosa* and *A. baumannii*, we employed *E. coli* K-12 MG1655 as a model to comprehensively investigate the regulatory mechanisms underlying this stress-induced polymyxin tolerance. We performed quantitative proteomic analysis on *E. coli* MG1655 cells subjected to the identical growth and treatment conditions as for the polymyxin B challenging assay (Figure S1). Among the 2990 proteins detected which cover 69% of the 4358 proteins annotated in the UniProt *E. coli* MG1655 database, abundance of 225 proteins was found to be altered following acute Pi depletion, with 142 proteins upregulated and 83 downregulated (Figure S3) (Table S2). In addition to proteins belonging to the Pho regulon, such as PhoB, PstB, PhoU, and PhnB (Figure 2A) in the COG functional group of transportation and metabolism of inorganic ions (13% of upregulated proteins) (Figure S3), the most abundant functional groups among the significantly upregulated proteins include those involved in cell wall/membrane/envelope biogenesis (13%) and energy production and conversion (10%) (Figure S3). Conversely, proteins involved in translation, ribosomal structure and biogenesis (33%) and amino acid transport and metabolism (17%) were significantly downregulated (Figure S3).

**Figure 2.**
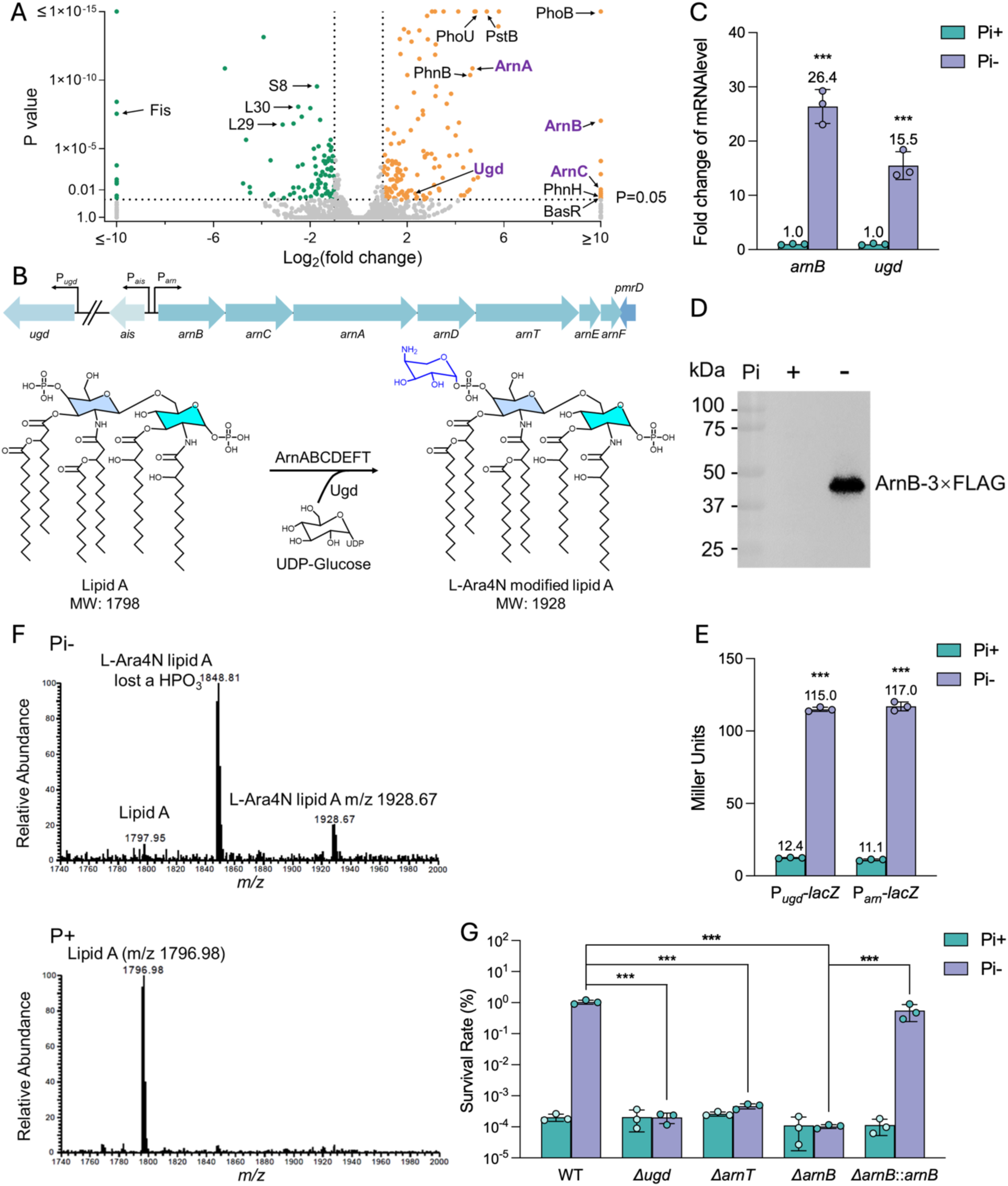
Pi depletion induces the expression of *ugd-arn* system to modify lipid A with L-Ara-4N, conferring stress-induced polymyxin tolerance. (A) Volcano plot depicting differentially expressed proteins in cells following acute Pi-starvation treatment (Pi-, 0 mM) compared to the group without treatment (Pi+, 1.32 mM). Orange (right) and green (left) dots represent significantly up-regulated and down-regulated proteins, respectively. The horizontal dashed line depicts a p-value cutoff (0.05), and vertical dashed lines depict 1/2 and 2-fold ratio cutoffs. Selected proteins are labeled. (B) Genetic organization of the *arn* operon and *ugd* gene in MG1655. Ugd and ArnABCDEFT catalyze the biosynthesis of 4-amino-4-deoxy-L-arabinose (L-Ara4N) and its modification of lipid A. (C) mRNA levels of *arnB* and *ugd* in cells following acute Pi-starvation treatment (Pi-, 0 mM) compared to the group without treatment (Pi+, 1.32 mM). (D) Production of ArnB-3×FLAG in cells following acute Pi-starvation treatment (Pi-, 0 mM) compared to the group without treatment (Pi+, 1.32 mM). The protein band corresponding to ArnB-3×FLAG is marked. (E) β-galactosidase activities of P*_arn_*-*lacZ* and P*_ugd_*-*lacZ* in MG1655 Δ*lacZ* cells following acute Pi-starvation treatment (Pi-, 0 mM) compared to the group without treatment (Pi+, 1.32 mM). (F) ESI-MS spectra of lipid A molecules extracted from *E. coli* MG1655 cells following acute Pi-starvation treatment (Pi-, 0 mM) compared to the group without treatment (Pi+, 1.32 mM). Peaks corresponding to native lipid A and L-Ara-4N modified lipid A were shown. (G) Survival rates for *E. coli* MG1655 parent and isogenic Δ*ugd,* Δ*arnT*, Δ*arnB* and Δ*arnB*::*arnB* mutants exposed 1-hour to polymyxin B (4 μg/ml) following acute Pi-starvation treatment (Pi-, 0 mM) compared to the group without treatment (Pi+, 1.32 mM). Error bars represent the standard deviation of at least three independent experiments. Statistical significance was determined using two-tailed Student’s t-test for **C** and **E**, and one-way ANOVA followed by Dunnett’s multiple comparisons test for **G**. Significance levels are indicated as follows: \**p* < 0.05; \*\**p* < 0.01; \*\*\**p* < 0.001.

Specifically, among the most upregulated proteins (log_2_FC>10) induced by Pi depletion (Table S2), two proteins encoded in the *arnBCADTEF* operon: ArnB and ArnC, were identified (Figures 2A and 2B). Additionally, two other proteins in the Arn pathway, ArnA and Ugd, were also up-regulated with more than five-fold in response to Pi depletion treatment (Figures 2A and 2B). This finding is particularly intriguing, because Ugd and Arn proteins are known to catalyze the biosynthesis of 4-amino-4-deoxy-L-arabinose (L-Ara4N) and its modification of lipid A (Figure 2B), which confers resistance to the cationic antibiotic polymyxins due to simultaneous neutralization of a negative charge and introduction of a positive charge to lipid A.^37^ To verify the upregulation of these proteins, we first examined the mRNA level of *ugd* and *arnB* genes by RT-qPCR and found their levels were 16- and 27-fold higher, respectively, under Pi-than under Pi+ conditions (Figure 2C). Next, we constructed ArnB-3×FLAG in MG1655 chromosome and observed a significant production of the ArnB protein under Pi-conditions, while its level was undetectable under Pi+ conditions (Figure 2D). We also constructed promoter-*lacZ* fusions and observed more than 10-fold higher promoter activities for both *ugd* and *arn* genes under Pi-conditions than Pi+ conditions (Figure 2E), confirming the upregulation of the Ugd-Arn system under Pi depletion.

We then extracted lipid A molecules from *E. coli* MG1655 cells cultured under identical conditions as in the proteomic analysis, and analyzed the extract using ESI-MS. Native lipid A, a hexacetylated glucosamine disaccharide with two phosphate groups at position 1 and 4’ (m/z 1796.98), was detected in *E. coli* cells cultured under Pi sufficient conditions (Figure 2F). In contrast, the predominant lipid A species in *E. coli* cells subjected to Pi depletion were found to be modified with L-Ara4N (m/z 1928.67) (Figure 2F). Next, we deleted *arnT*, *arnB*, and *ugd* and found that deletion of these genes abolished Pi depletion-induced polymyxin B tolerance in *E. coli* (Figure 2G). To confirm the linkage between the observed phenotypes and the targeted genes, we complemented the Δ*arnB* mutant by integrating a native promoter-driven copy of *arnB* at the attTn7 site. This complementation restored the phenotype to wild-type levels, demonstrating that the loss of *arnB* was responsible for the observed defect (Figure 2G).

The induction of L-Ara4N modified lipid A, along with decrease in the unmodified form, was also observed in *E. coli* O157 (Figure S4A), a representative UTI strain UTI-3 (Figure S4B), and *S.* Typhimurium ATCC14028 (Figure S4C), each exhibiting varying ratios. These findings indicate that L-Ara4N modification of lipid A is widespread in *Enterobacteriaceae* and correlates with increased polymyxin B tolerance. We also analyzed lipid A species in *P. aeruginosa* PAO1 and found that instead of the L-Ara4N modification, a 2-hydroxymyristate modification of lipid A, which is catalyzed by the aspartyl beta-hydroxylase LpxO, was detected in Pi-depletion treated cells (Figure S4D). Altogether, these results indicated that unlike the strategy observed in PAO1 where Pi stress induces tolerance by replacing the negatively charged phospholipids with MGDG/GADG, in *E. coli*, tolerance is mediated by lipid A modification with L-Ara4N catalyzed by Ugd and Arn proteins.

To delineate the sensitivity of the Ugd-Arn pathway to varying Pi levels, we measured the promoter activities of P*_ugd_-lacZ* and P*_arn_-lacZ* across a gradient of phosphate concentrations. We observed that the activities of both P*_ugd_-lacZ* and P*_arn_-lacZ* remained robustly activated, at levels comparable to complete Pi depletion (0 mM), even when 0.040 mM or 0.132 mM Pi was supplemented to the medium (Figure S5). When Pi supplementation reached 0.4 mM, their upregulation was attenuated by 60%. These findings collectively suggest that moderate Pi limitation (below 0.4mM) is effective in sustaining the activation of *ugd-arn* genes (Figure S5). To determine whether activation of the Ugd-Arn pathway under phosphate limitation translates into increased polymyxin resistance, we measured polymyxin B MICs under phosphate-replete (1.32 mM Pi) and phosphate-limited (0.132 mM Pi) conditions. Despite strong induction of the pathway at 0.132 mM Pi (Figure S5), no significant change in MIC was observed for the wild-type, mutant, or complemented strains (Table S3). Together with the enhanced survival observed in polymyxin challenging assays (Figure 1), these results further confirm that phosphate limitation promotes polymyxin tolerance rather than classical resistance.

### The PmrAB two component system is activated to up-regulate the *arn* operon during Pi depletion

To elucidate the regulatory mechanisms underlying *ugd-arn* upregulation during Pi limitation, we first investigated the involvement of the global phosphate regulators PhoB and PhoR. We found that deleting either PhoB or PhoR did not affect the enhanced polymyxin B tolerance or the upregulation of the *arn* operon observed in Pi-depleted *E. coli* (Figures 3A and S6). This observation suggested that other regulatory systems were primarily responsible for upregulating the *arn* gene expression under Pi depletion.

**Figure 3.**
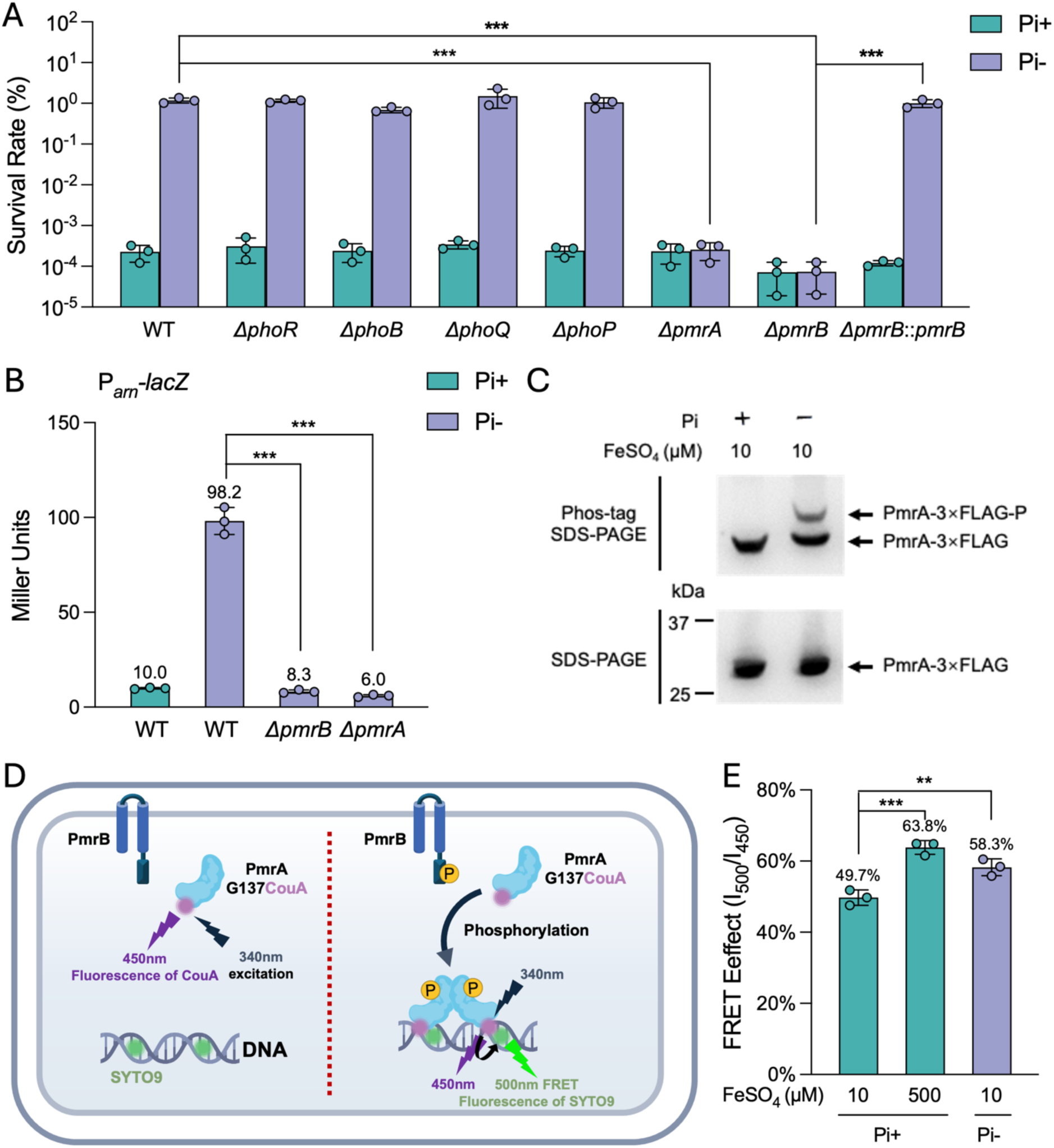
The PmrAB two-component system is activated during Pi depletion and is responsible for the up-regulation of *arn* genes under this condition. (A) Survival rates for *E. coli* MG1655 parent and isogenic, Δ*phoR,* Δ*phoB,* Δ*phoQ,* Δ*phoP,* Δ*pmrA,* Δ*pmrB* and Δ*pmrB*::*pmrB* mutants exposed 1-hour to polymyxin B (4 μg/ml) following acute Pi-starvation treatment (Pi-, 0 mM) compared to the group without treatment (Pi+, 1.32 mM). (B) β-galactosidase activities of P*_arn_-lacZ* in MG1655 Δ*lacZ* parent and isogenic Δ*pmrB*, Δ*pmrA,* Δ*phoB,* and Δ*phoR* mutants following acute Pi-starvation treatment (Pi-, 0 mM) compared to the group without treatment (Pi+, 1.32 mM). (C) Phosphorylation of PmrA-3×FLAG detected by Phos-Tag SDS–PAGE following immunoblotting with anti-FLAG antibody in cells following acute Pi-starvation treatment (Pi-, 0 mM) compared to the group without treatment (Pi+, 1.32 mM). Protein bands representing PmrA-3×FLAG and PmrA-3×FLAG-P are indicated. (D) Schematic diagram of the whole-cell FRET measurement of PmrA-promoter binding assay. A bright fluorescent unnatural amino acid donor CouA (red spheres) is incorporated into G137 of PmrA in DNA-binding domain. SYTO9 (green spheres) is employed to stain chromosome DNA in living *E. coli* MG1655. (E) Quantification of the FRET effect between the CouA-SYTO9 pair in *E. coli* cells following acute Pi-starvation treatment (Pi-, 0 mM Pi, 10 μM Fe) or iron excess treatment (Pi+, 1.32 mM Pi, 500 μM FeSO_4_) compared to the group without treatment (Pi+, 1.32 mM Pi, 10 μM FeSO_4_). Error bars represent the standard deviation of at least three independent experiments. Statistical significance was determined using one-way ANOVA followed by Dunnett’s multiple comparisons test for **A**, **B**, and **E**. Significance levels are indicated as follows: \**p* < 0.05; \*\**p* < 0.01; \*\*\**p* < 0.001.

The *arn* operon and *ugd* are known targets of the PmrAB two-component system (TCS) in *E. coli.* ^38^ (Note the *E. coli* homolog of this system is BasSR; we adopt the PmrAB nomenclature in this study to reflect its direct role in polymyxin tolerance and its wider usage in the literature.)^39^ To determine if this regulation also extended to Pi depletion, we constructed Δ*pmrB* and Δ*pmrA* mutants. Both the enhanced polymyxin B tolerance and the upregulation of the *arn* operon under Pi depletion were abolished in these mutants (Figures 3A and 3B). To confirm the linkage between the observed phenotypes and the targeted genetic lesions, we complemented the Δ*pmrB* mutant by introducing a native promoter-driven copy of *pmrB* at the attTn7 site. This complementation restored polymyxin B tolerance to near wild-type levels (Figure 3A). Together, these results indicate a critical role for PmrAB in *arn* regulation and polymyxin B tolerance during Pi depletion. The promoter activity of *ugd* remained unaffected in Δ*pmrB* and Δ*pmrA* mutants (Figure S7), suggesting a distinct regulatory input for *ugd*. Given that the PhoPQ TCS can indirectly activate the PmrAB regulon in *S*. Typhimurium via PmrD (though this indirect regulation appears absent in *E. coli*), we also examined the contribution of PhoPQ.^40^ Deletion of *phoP* or *phoQ* did not alter the enhanced polymyxin B tolerance or *arn* operon upregulation under Pi depletion (Figures 3A and S6). These findings suggest that the *arn* system is activated by PmrAB in response to an as-yet-unidentified secondary signal induced by Pi depletion, rather than by direct Pi depletion sensing or through the PhoPQ system.

Given the distinct regulation we observed for *ugd* promoter activity, we further investigated its response to the PhoBR system. We measured *ugd* promoter activity in Δ*phoB*, Δ*phoR*, Δ*phoP*, and Δ*phoQ* mutants and found P*_ugd_-lacZ* activity decreased by approximately 30% in both Δ*phoB* and Δ*phoR* mutants, but not in Δ*phoP* and Δ*phoQ* mutants (Figure S7). This indicates that *ugd* is regulated by the PhoBR system, potentially alongside other pathways, during Pi depletion.

Given the central role of the PmrAB system in mediating Pi depletion-induced polymyxin tolerance, we next investigated whether *eptA*, another PmrAB-regulated gene, contributes to this phenotype. We constructed a Δ*eptA* mutant and assessed bacterial survival in a polymyxin B killing assay. Deletion of *eptA* had no impact on the enhanced polymyxin B tolerance (Figure S8), indicating that *eptA* is not required for the tolerance phenotype induced by Pi depletion. This observation was further supported by ESI-MS analysis (Figure 2F), which did not detect phosphoethanolamine (pEtN) modification of lipid A, a hallmark of EptA activity, in Pi-depleted cells. Collectively, these results strongly suggest that the activation of the Arn system via PmrAB is the primary mechanism responsible for Pi depletion-induced polymyxin tolerance.

To assess whether PmrAB activation occurs under Pi depletion, we examined the phosphorylation status of the response regulator PmrA using Phos-tag SDS-PAGE, which reliably detects phosphorylated PmrA in cells treated with known activator Fe^3+^.^41^ Results showed that phosphorylated PmrA was detected in cells subjected to Pi depletion (Figure 3C), demonstrating that PmrAB TCS is indeed activated under Pi depletion condition. To further confirm this in vivo, we performed a FRET-based whole-cell transcription factor-promoter binding assay.^42^ We incorporated a bright fluorescent unnatural amino acid, L-(7-hydroxycoumarin-4-yl) ethylglycine (CouA)^43^ at position G137 within the DNA-binding domain of PmrA through genetic encoding, allowing it to serve as the FRET donor (Figure 3D).^42^ The nucleic acid dye SYTO9 was utilized to stain promoter DNA of PmrA regulon (Figure 3D),^42^ acting as the FRET acceptor. The whole-cell FRET assay revealed that, similar to the treatment with excessive FeSO_4_ (500μM),^42^ a FRET signal indicative of PmrA binding to its target promoters was observed in cells subjected to Pi depletion without excessive FeSO_4_ (Figure 3E). These results confirmed that Pi depletion induced activation of the PmrAB signal transduction system in *E. coli*.

### Pi depletion drives Fe^3+^ elevation in *E. coli* cells to activate the PmrAB TCS

PmrAB has been previously reported to be activated by ferric ion (Fe^3+^), zinc ions (Zn²⁺), aluminum ions (Al^3+^), or acidic pH.^44,45^ Since no Al^3+^ was added in the culture, and the pH remained largely unchanged (7.2-7.0) across the Pi depletion treatments (Figure S9), we focused on the two metal ions and examined cellular metal profiles including iron (Fe), zinc (Zn), magnesium (Mg), copper (Cu), and calcium (Ca). ICP-MS analysis revealed a significant, greater than 5-fold increase in cellular Fe content under Pi depletion, compared to Pi-sufficient conditions (Figure 4A). Conversely, cellular Mg levels decreased, while Cu, Zn, and Ca levels remained largely unchanged (Figure 4A). Corroborating this cellular Fe accumulation, the Fe content in the spent medium decreased after 3 hours of *E. coli* incubation under Pi depletion (Figure 4B), indicating active Fe mobilization from the medium into *E. coli* cells. These findings strongly implicated Fe^3+^ as the activating signal for PmrAB under Pi depletion.

**Figure 4.**
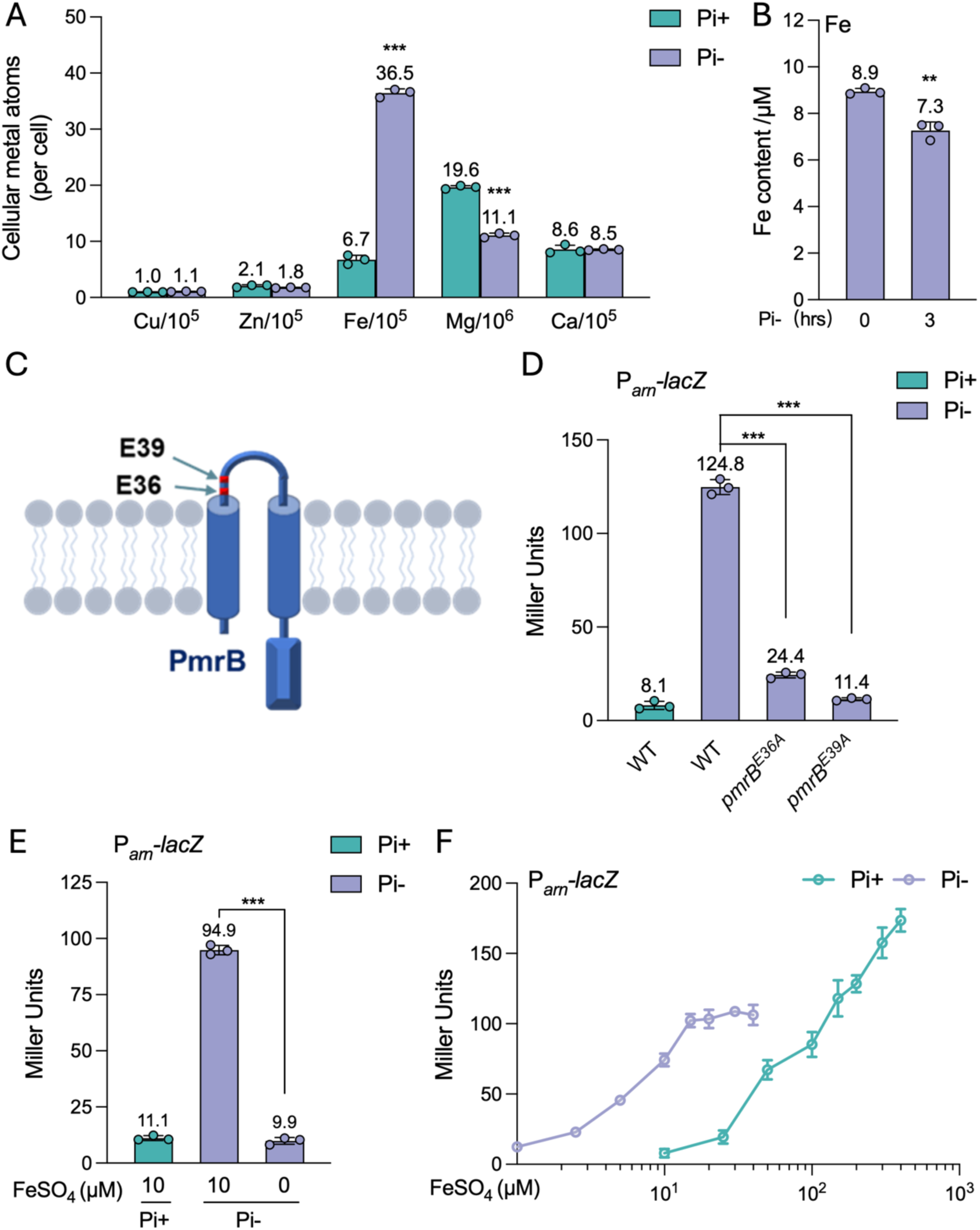
Activation of PmrAB during Pi depletion is mediated by the secondary stress signal iron (Fe^3+^) induced under this condition. (A) Cellular metal contents determined by ICP-MS in *E. coli* MG1655 cells following acute Pi-starvation treatment (Pi-, 0 mM) compared to the group without treatment (Pi+, 1.32 mM). (B) Iron contents in the spent medium of *E. coli* MG1655 at the indicated time points during Pi depletion treatment (Pi-, 0 mM). (C) Schematic diagram of the sensor kinase PmrB with two important residues for sensing Fe^3+^, E36A and E39A, are shown. (D) β-galactosidase activities of P*_arn_-lacZ* in MG1655 Δ*lacZ* parent and isogenic *pmrB*^E36A^ and *pmrB*^E36A^ mutants following acute Pi-starvation treatment (Pi-, 0 mM) compared to the group without treatment (Pi+, 1.32 mM). (E) β-galactosidase activities of P*_arn_-lacZ* in MG1655 Δ*lacZ* cells following Pi depletion treatment (Pi-, 0 mM Pi, 10 μM Fe) or Fe, Pi depletion treatment (Pi-Fe-, 0 mM Pi, 0 μM Fe) compared to the group without treatment (Pi+, 1.32 mM Pi, 10 μM FeSO_4_). (F) β-galactosidase activities of P*_arn_-lacZ* in MG1655 Δ*lacZ* grown under Pi sufficient MOPS medium in the presence of different concentrations of FeSO_4_ and that subjected to Pi depletion conditions in the presence of different concentrations of FeSO_4_. Error bars represent the standard deviation of at least three independent experiments. Statistical significance was determined using one-way ANOVA followed by Dunnett’s multiple comparisons test for **B**, and two-tailed Student’s t-test for **C**, **D**, and **E**. Significance levels are indicated as follows: \**p* < 0.05; \*\**p* < 0.01; \*\*\**p* < 0.001.

Given that *S.* Typhimurium PmrB (StyPmrB) senses Fe^3+^ through conserved glutamic acid residues (E36, E39, E61, E64) in its periplasmic domain,^41^ we hypothesized a similar mechanism for *E. coli* PmrB (EcoPmrB). To test this, we generated chromosomal *pmrB*^E36A^ and *pmrB*^E39A^ mutations in *E. coli* MG1655 (Figure 4C). Consistent with their proposed role in Fe^3+^ sensing, these mutations abolished the P*_arn_*-*lacZ* reporter response to exogenous Fe^3+^ (100 µM) (Figure S10). Crucially, both mutations also substantially reduced *arn* operon upregulation under Pi depletion (Figure 4D). Western blot analysis showed that the two PmrB variants were produced to levels comparable to the WT PmrB, indicating that their loss of Pi depletion-induced signaling in these mutants results from impaired Fe³⁺ sensing rather than reduced protein abundance or stability (Figure S11). Together, these findings demonstrate that Pi depletion-induced PmrAB activation requires the Fe^3+^ sensing capability EcoPmrB.

To confirm the necessity of Fe mobilization, we observed that *arn* operon upregulation was abolished when cells were grown in Fe-free, Pi-depleted medium (Figure 4E). Although Pi depletion significantly impaired bacterial growth (doubling time increased from ∼1 hr to ∼6 hrs), co-depletion of Fe did not further exacerbate this effect (doubling time ∼6 hrs), confirming that the observed PmrAB activation specifically responds to Pi-related Fe mobilization rather than general growth inhibition. Investigating the sensitivity of this response, we found that reintroduction of Fe to Pi-depleted cells restored P*_arn_*-*lacZ* activity in a dose-dependent manner (Figure 4F), further confirming the role Fe mobilization in activating PmrAB and *arn* genes during Pi depletion. Importantly, *arn* activation in Pi-starved cells occurs at significantly lower exogenous Fe concentrations (∼10-fold less) than in Pi-sufficient conditions (Figure 4F), suggesting an enhanced cellular capacity for Fe mobilization or sensing under Pi depletion.

### Pi depletion triggers Mg^2+^ loss, leading to Fe^3+^ mobilization to membrane and PmrAB activation

To elucidate the mechanism of Fe mobilization during Pi depletion, we first investigated the role of canonical iron uptake pathways. Surprisingly, deletion of genes encoding outer membrane iron-chelator complex receptors and enterobactin synthetase (*fecA*, *fepA*, *fiu*, *cirA*, *entE*), individually or as a penta-deletion mutant (Δ5Fe), did not abolish P*_arn_*-*lacZ* induction during Pi depletion (Figure S12A). ICP-MS analysis further confirmed that cell-associated Fe still increased by over 5-fold in Δ5Fe cells during Pi depletion (Figure S12B). Proteomic analysis revealed that the major iron-storage proteins Bfr, Dps, and FtnA were also upregulated modestly during Pi depletion, with abundances increasing 2- to 4-fold relative to Pi-sufficient conditions (Table S3). We therefore tested a triple mutant lacking all three proteins (Δ*bfr* Δ*dps* Δ*ftnA*) to determine whether they contribute to Fe accumulation or Fe-dependent PmrAB activation. Neither cellular Fe accumulation nor P*_arn_–lacZ* induction during Pi depletion was significantly altered in this strain (Figures S13A and S13B), indicating that Bfr, Dps, and FtnA are dispensable for both processes. Together, these results suggest that Fe-mediated PmrAB activation during Pi depletion does not depend on enhanced cellular Fe uptake or sequestration by the major iron-storage proteins. We therefore investigated whether Fe is instead mobilized to the cell envelope, consistent with the ability of the PmrB sensor kinase to respond to periplasmic or surface-associated Fe³⁺.^41^

The outer leaflet of the Gram-negative outer membrane (OM) primarily consists of lipid A, whose negatively charged phosphate groups are typically neutralized by cell-associated Mg^2+^ and Ca^2+^ to maintain membrane stability.^46^ Our ICP-MS analysis revealed a 44% reduction in cellular Mg content, while Ca^2+^ levels remained unchanged, in *E. coli* cells following 3 hours of Pi depletion treatment (Figure 4A). Monitoring the dynamics revealed that cellular Mg progressively decreased upon transferring *E. coli* cells to Pi-depleted medium, continuing over a 5-hour period (Figure 5A). This Mg^2+^ loss was independent of Fe, as a similar degree of loss occurred in Fe-free, Pi-depleted medium (Figure S14). Conversely, cellular Fe increased, peaking after two hours of Pi depletion before slightly declining (Figure 5A). The temporal correlation between Fe accumulation and P*_arn_*-*lacZ* activity (Figure 5B) suggested active Fe mobilization to the cell surface during Pi depletion, potentially to compensate for Mg^2+^ loss and stabilize the membrane. The subsequent activation of the PmrAB-Arn system that remodel lipid A by decreasing negative charges, might thus represent an adaptive response to physiological changes, such as OM instability, caused by Mg^2+^ dissociation. Supporting this, deletion of *pmrA* led to an even greater and sustained increase in Fe mobilization during Pi depletion (Figure 5C), which could be toxic to *E. coli*.^41^

**Figure 5.**
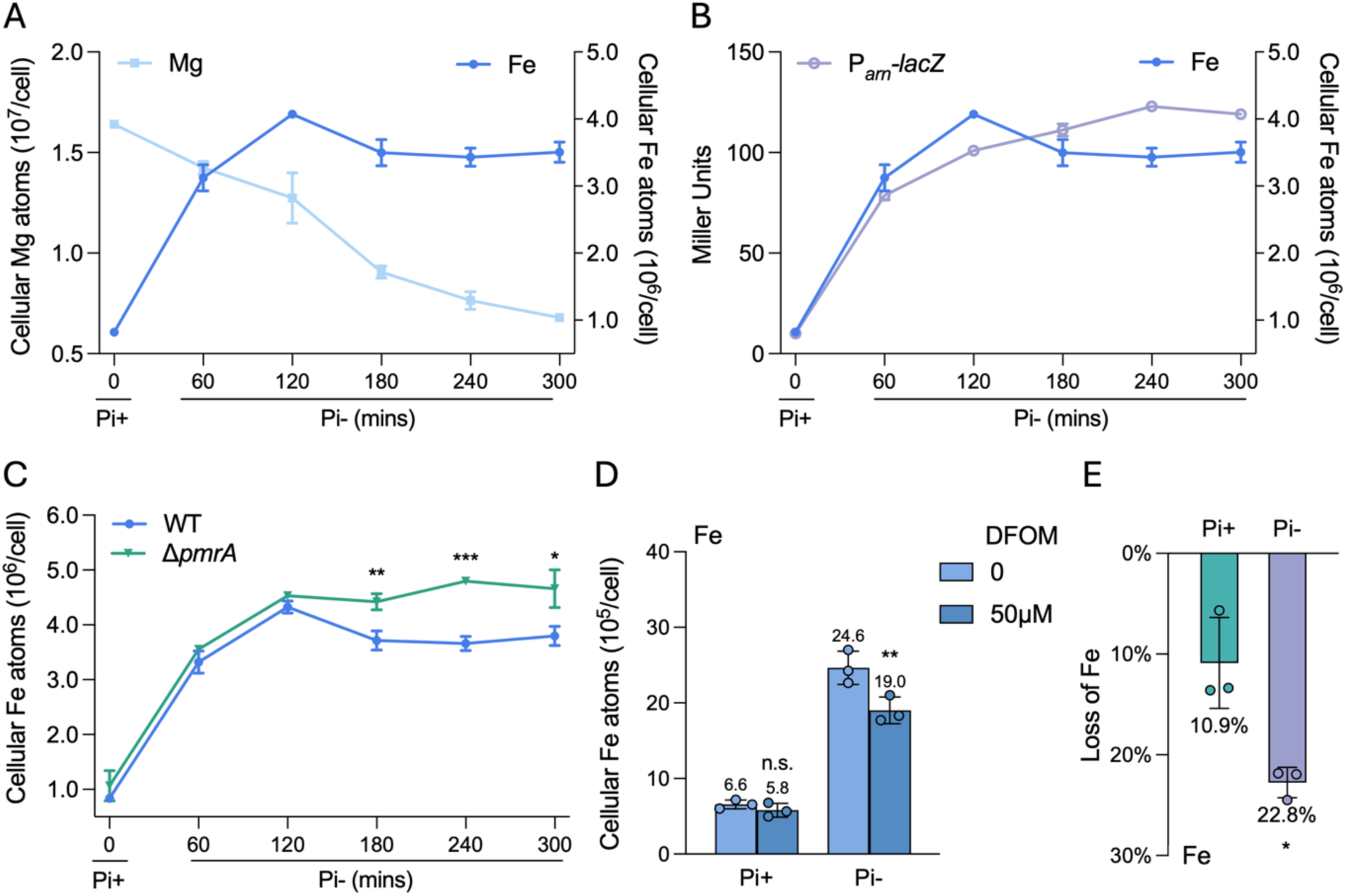
Response to Pi depletion induced iron mobilization to cell surfaces. (A) Iron (Fe) and magnesium (Mg) contents in *E. coli* MG1655 cells following Pi depletion treatment (Pi-, 0 mM Pi) at the indicated time points. (B) β-galactosidase activities of P*_arn_-lacZ* in MG1655 Δl*acZ* cells following Pi depletion treatment at the indicated time points. Fe content data from panel A are reproduced here for direct comparison with P*_arn_-lacZ* activity. (C) Fe contents in *E. coli* MG1655 parent and isogenic Δ*pmrA* mutant cells following Pi depletion treatment (Pi-, 0 mM Pi) at the indicated time points. (D) Fe contents in *E. coli* MG1655 cells with Pi depletion treatment (Pi-, 0 mM) and without Pi depletion treatment (Pi+, 1.32 mM) following washing by HEPES or DFOM-supplemented (50μM) HEPES buffer. (E) Reduction of cellular Fe content following washing with DFOM-supplemented (50μM) HEPES buffer. Error bars represent the standard deviation of at least three independent experiments. Statistical significance was determined using two-tailed Student’s t-test for **C** and **E**, and two-way ANOVA followed by Sidak’s multiple comparisons test for **D**. Significance levels are indicated as follows: \**p* < 0.05; \*\**p* < 0.01; \*\*\**p* < 0.001.

To directly assess Fe association with the cell surface, we washed cell pellets with deferoxamine (DFOM)-containing buffer, a membrane-impermeable Fe^3+^-specific chelator.^47^ ICP-MS analysis showed that washing with DFOM-supplemented buffer significantly reduced Fe content in cells treated with Pi depletion but had no significant effect on cells harvested from Pi-sufficient conditions (Figure 5D). Quantification revealed that Pi-starved cells exhibited a two-fold greater loss of Fe upon DFOM washing compared to Pi-sufficient conditions (Figure 5E), demonstrating that Pi depletion promoted Fe mobilization to cell surfaces, where it remained susceptible to removal by a membrane-impermeable chelator.

To identify the mechanism driving enhanced Fe mobilization, we supplemented Pi-depleted cultures with excess Mg²⁺ (10 mM) to compensate for cellular Mg²⁺ loss. Mg²⁺ supplementation markedly reduced Fe association with MG1655 cells (Figure 6A), and P*_arn_–lacZ* activity decreased in a concentration-dependent manner as Mg²⁺ levels increased (Figure 6B). We next asked whether this effect was specific to Mg²⁺ or instead reflected restoration of divalent-cation-mediated envelope charge neutralization. Supplementation with Ca²⁺, another divalent cation associated with the bacterial cell envelope, similarly suppressed P*_arn_–lacZ* activity in a concentration-dependent manner under Pi-depleted conditions (Figure S15). Although ICP–MS detected depletion of cellular Mg²⁺ but not Ca²⁺ (Figure 4A), Mg²⁺ is approximately three orders of magnitude more abundant than Ca²⁺ in the culture medium; therefore, changes in cellular Ca²⁺ content may have been too small to detect under our experimental conditions. Together, these findings support a model in which Pi depletion disrupts cellular Pi–Mg homeostasis, leading to Mg²⁺ loss and a consequent deficit in divalent-cation-mediated envelope charge neutralization; Fe is then mobilized to the cell envelope to compensate for this charge imbalance, whereas exogenous Ca²⁺ can substitute for Mg²⁺ and diminish the requirement for Fe mobilization. Consistent with this model, Mg²⁺ concentrations in the spent medium increased continuously during Pi depletion (Figure 6C), indicating net Mg²⁺ release from the cells. Mg²⁺ release also occurred when both Mg²⁺ and Pi were limited (Figure S16), suggesting that it is associated with cellular Pi–Mg imbalance during Pi starvation rather than simply reflecting extracellular Mg²⁺ abundance. A comparable loss of cellular Mg²⁺ (∼45%) was observed in *P. aeruginosa* PAO1 under the same conditions (Figure S17). However, Fe mobilization was less pronounced in PAO1 than in *E. coli* (2.4-fold versus 5.4-fold), consistent with species-specific adaptations to Pi limitation^11^.

**Figure 6.**
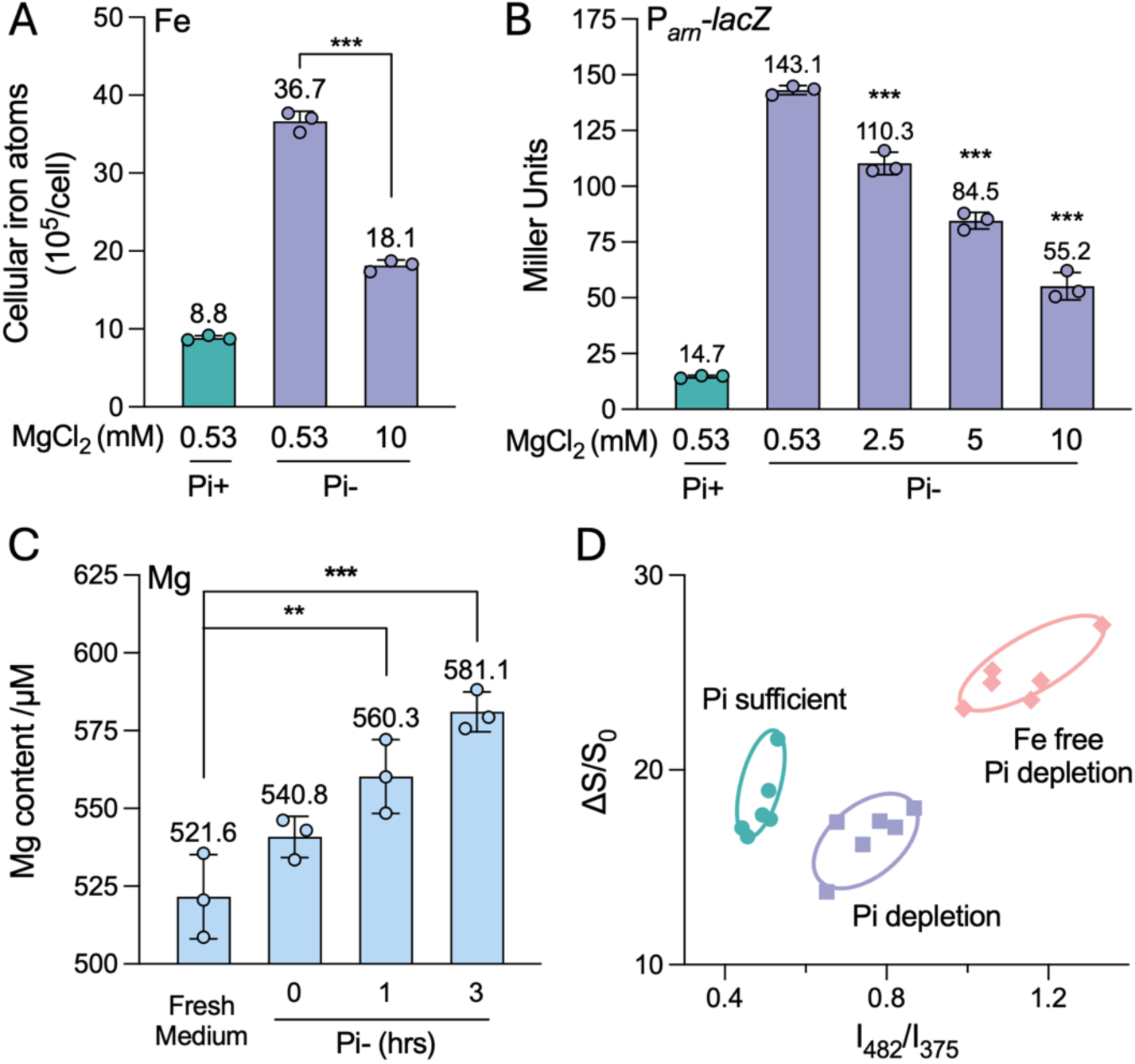
Iron mobilization is driven by Mg^2+^ dissociation during Pi depletion. (A) Fe contents in *E. coli* MG1655 cells following acute Pi-starvation treatment (Pi-, 0 mM Pi, 0.53 mM Mg) or Mg excess Pi depletion treatment (Pi-, 0 mM Pi, 10 mM Mg) compared to the group without treatment (Pi+, 1.32 mM, 0.53 mM Mg). (B) β-galactosidase activities of P*_arn_-lacZ* in MG1655 Δ*lacZ* cells following acute Pi-starvation treatment (Pi-, 0 mM Pi, 0.53 mM Mg) or Mg excess Pi depletion treatment (Pi-, 0 mM Pi, 0.53-10 mM Mg) compared to the group without treatment (Pi+, 1.32 mM, 0.53 mM Mg). (C) Mg content in the spent medium during Pi depletion treatment at the indicated time points. (D) 2D graph plotted from output signals of two channels of the PI-BactD sensor: fluorescence increase (ΔS/S0) and fluorescence radiometric changes (I482/I375) in each of the treatments indicated. Data are presented as mean ± s.d. of three biological replicates for **A**, **B**, and **C**, and six biological replicates for **D**. Error bars represent the standard deviation of at least three independent experiments. Statistical significance was determined using two-tailed Student’s t-test for **a**, and one-way ANOVA followed by Dunnett’s multiple comparisons test for **B** and **C**. Significance levels are indicated as follows: \**p* < 0.05; \*\**p* < 0.01; \*\*\**p* < 0.001.

Given that membrane-associated Mg^2+^ primarily neutralizes the negative charges of lipid A to stabilize the OM, we hypothesized that Mg^2+^ loss during Pi depletion alters OM properties, and Fe mobilization may serve to stabilize the OM. To test this, we employed an OM-sensitive probe.^48^ The probe reports membrane surface properties through two orthogonal readouts: an overall fluorescence increase, which reflects probe disassembly upon binding to the anionic bacterial surface, and the pyrene excimer-to-monomer emission ratio (I_482_/I_375_), which captures differences in local surface interactions. The 2D sensor output revealed distinct signals between cells grown under Pi-sufficient and Pi-depleted conditions (Figure 6D), indicating altered OM characteristics. Notably, cells grown in Fe-free, Pi-depleted medium exhibited even greater shifts (Figure 6D), suggesting that Fe mobilization and subsequent activation of the PmrAB-Arn pathway served as an adaptation strategy to maintain OM integrity during Pi stress. This strategy likely prevents uncontrolled Fe accumulation and toxicity,^41^ ultimately resulting in stress-induced polymyxin tolerance. A schematic model illustrating this process is presented (Figure 7).

**Figure 7.**
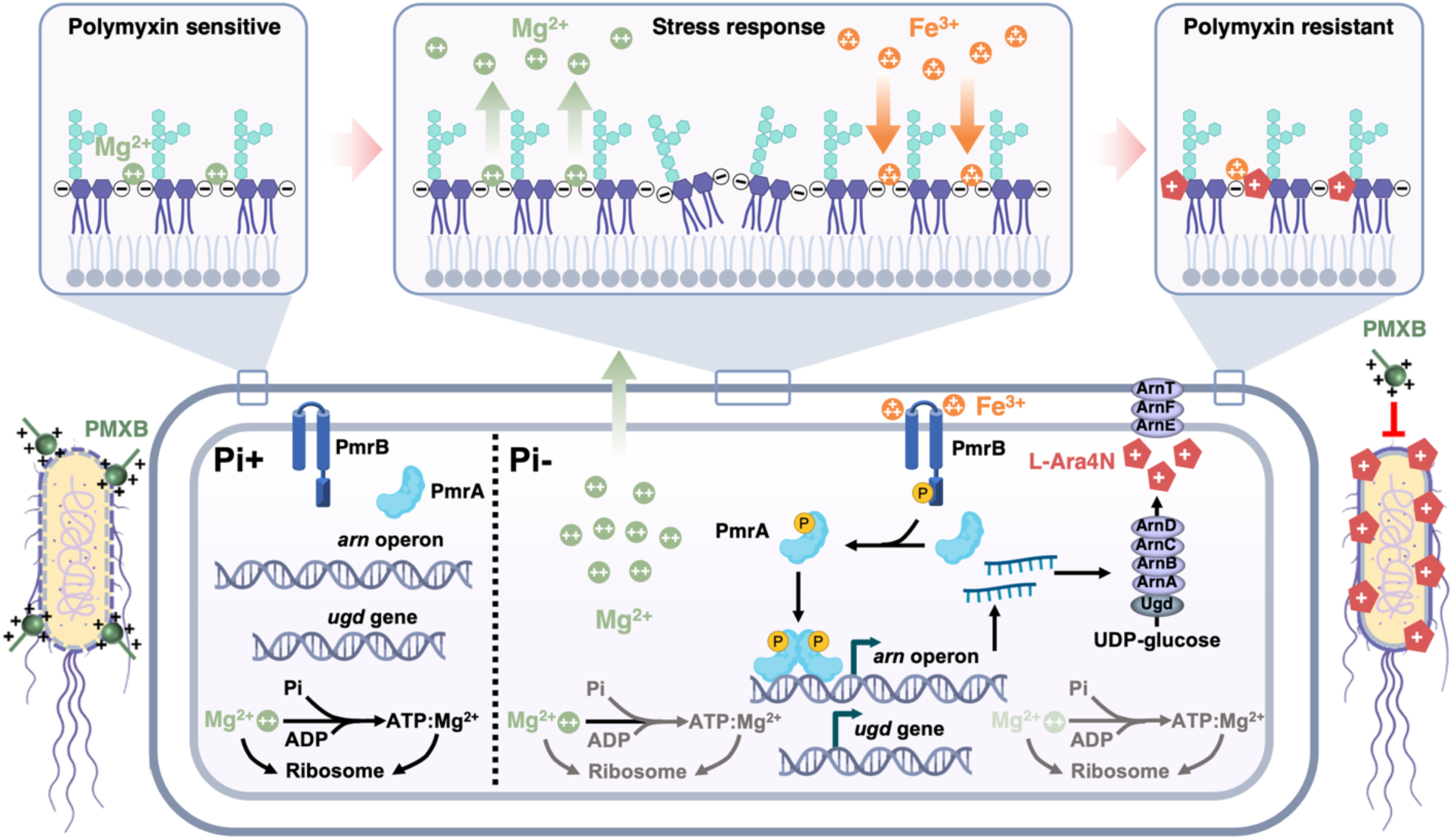
A Mg-Fe-PmrAB regulatory circuit underpins the Pi-depletion induced polymyxin tolerance through activating the lipid A L-Ara4N modification system. In cell grown under Pi sufficient condition, ribosome and ATP serve as the major intracellular Mg^2+^ and Pi reservoir. Mg^2+^ neutralizes negative charge of lipid A and stabilizes outer membrane (OM). The PmrAB two-component system (TCS) is inactive, and the cell is sensitive to polymyxin. During Pi stress, ATP biosynthesis and ribosome biogenesis are halted due to Pi depletion, disrupting the balance between Pi and Mg^2+^. Mg^2+^ disassociates from the cell which destabilizes the OM and derives compensatory Fe^3+^ mobilization to cell surface. Mobilized Fe^3+^ activates the PmrAB TCS which in turn upregulates the *arn* operon. Arn and Ugd proteins catalyze the biosynthesis of L-Arn4N and modify lipid A with L-Arn4N to stabilize OM under stressed condition and prevent further Fe mobilization. *E. coli* cells with L-Arn4N modified lipid A become resistant to polymyxin.

### Disrupting the Mg-Fe-PmrAB-Arn circuit counters Pi depletion-induced polymyxin tolerance in *Enterobacteriaceae*

Our findings above identified a Mg-Fe-PmrAB-Arn circuit underpinning the polymyxin tolerance phenotype induced by stress adaptation to Pi depletion. To evaluate whether interference with this pathway could mitigate the stress-induced polymyxin tolerance, we supplemented Pi depletion medium with excess Mg^2+^ (10mM) or applied a membrane impermeable chelator DFOM (10μM). Both approaches effectively suppressed the Pi depletion-induced polymyxin tolerance phenotype in *E. coli* and *S.* Typhimurium (Figure 8A). In contrast, neither intervention reversed the Pi depletion-associated polymyxin tolerance phenotype in *P. aeruginosa* despite various combinations of Mg^2+^ supplementation and Fe chelation (Figure S18). This highlights species-specific adaptative strategies and aligns with several observations from our study as well as several previous research.^11,49^ Specifically, in *P. aeruginosa,* despite a 45% Mg loss during Pi depletion, Fe levels increased only two-fold compared to over five-fold in *E. coli*, consistent with its reliance on MGDG/GADG lipids synthesis regulated by the PhoBR system (Figure 8B).^11^ Additionally, *P. aeruginosa* exhibits a distinct lipid A modification, 2-hydroxy myristate (Figure S4D), that does not confer polymyxin tolerance.^50^ Phylogenetic analysis of 250 PmrB homologs revealed that EcoPmrB and PaePmrB cluster in separate clades, with PaePmrB lacking the conserved “ExxE” motif essential for Fe³⁺ sensing (Figure 8C). Together, these findings underscore the diverse stress response strategies employed by Gram-negative bacteria to adapt to Pi limitation and highlight the importance of species-specific regulatory mechanisms in shaping these responses. While the present study establishes the mechanistic basis of this pathway in *E. coli* and demonstrates similar phenotypic responses in *S. Typhimurium*, the extent to which related mechanisms operate in other members of the *Enterobacteriaceae* remains to be determined. Nevertheless, the broad conservation of the PmrAB signaling system and key Fe-sensing residues among many *Enterobacteriaceae* species raises the possibility that analogous regulatory mechanisms may exist beyond *E. coli*.

**Figure 8.**
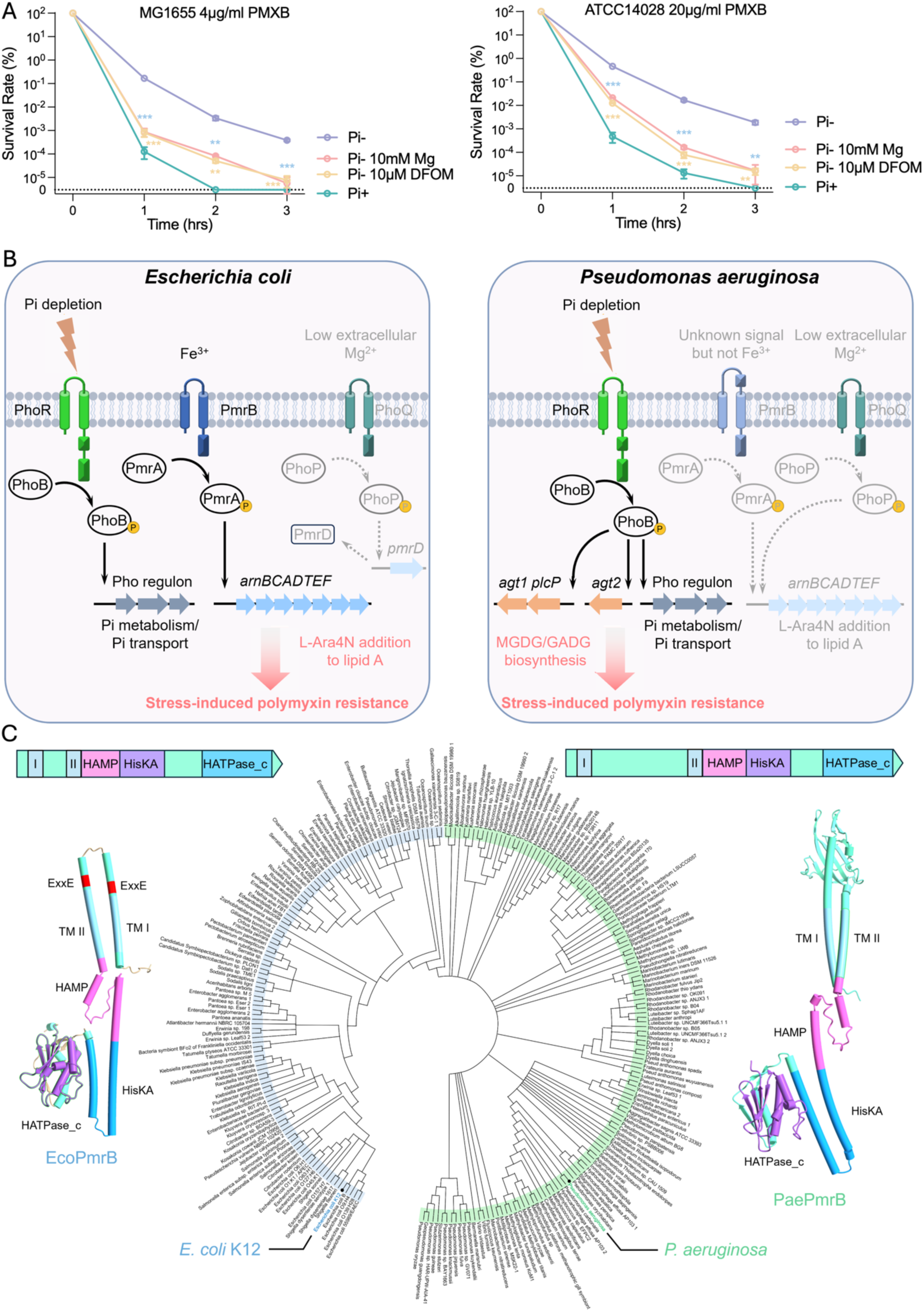
Disrupting the Mg-Fe-PmrAB signaling circuit attenuated Pi-depletion induced polymyxin tolerance in *Enterobacteriaceae* (A) Time-kill curves of *Escherichia coli* MG1655 (left panel) and *Salmonella*. Typhimurium ATCC14028 (right panel) exposed to polymyxin B (PMXB) (4 μg/ml for *E. coli* strains, 20 μg/ml for *S.* Typhimurium) following acute Pi-starvation treatment (Pi-, 0 mM Pi), Pi depletion treatment with excess Mg (Pi-, 0 mM Pi, 10mM MgCl_2_) or Pi depletion treatment with iron chelator (Pi-, 0 mM Pi, 10 μM DFOM) compared to the group without treatment (Pi+, 1.32 mM Pi). The dotted line labelled “0” denotes the limit of detection; values below this threshold were not detectable by colony counting. (B) Distinct stress adaptation strategies in *E. coli* and *P. aeruginosa* under Pi depletion conditions. Genes and regulatory systems activated in the two species are illustrated. Inactive regulatory systems are depicted in faded effects. The regulatory mechanisms for *P. aeruginosa* are based on findings from Jones et. al.^11^ (C) A phylogenetic tree of 250 PmrB homologous. A predominant clade is indicated by a blue circle, comprising sequences from bacteria such as *Escherichia, Shigella, Citrobacter, Salmonella, Klebsiella, Yersinia*, and *Enterobacter* species. A green circle marks several clades distinct from the predominant one. The domain architecture and AlphaFold3-predicted structure of EcoPmrB and PaePmrB are shown. Error bars represent the standard deviation of at least three independent experiments. Statistical significance was determined using one-way ANOVA followed by Dunnett’s multiple comparisons test. Significance levels are indicated as follows: \**p* < 0.05; \*\**p* < 0.01; \*\*\**p* < 0.001.

## Discussion

Bacterial adaptation to environmental stressors is crucial for survival and pathogenesis. Phosphate (Pi) scarcity, a pervasive condition in diverse environments including host-associated niches such as macrophage phagosomes, salmonella-containing vacuoles, and cystic fibrosis sputum, is a potent cue altering antimicrobial susceptibility against cationic antibiotics such as polymyxins.^9-11^ Adaptive mechanisms previously characterized for Pi depletion-induced antimicrobial tolerance often involve remodeling bacterial membranes by substituting phosphorus-containing phospholipids with Pi-free, neutral, or positively-charged lipids.^11,26–28^ However, the specific mechanisms in clinically important *Enterobacteriaceae*, which largely lack these canonical phospholipid remodeling pathways, have remained largely unknown. In this study, we unveil a conserved, yet distinct, polymyxin B tolerance mechanism in *Enterobacteriaceae*: Pi limitation triggers a cascade of cationic imbalance and Fe^3+^ signaling, leading to L-Ara4N modification of lipid A and outer membrane remodeling, a process distinct from phospholipid-based tolerance pathways (Figure 7). This intricate Mg-Fe-PmrAB regulatory circuit represents a previously unappreciated ecophysiological vulnerability, offering a novel strategy to mitigate stress-induced polymyxin tolerance.

Maintaining a balanced P/Mg ratio is paramount for bacterial cellular physiology, with ATP and rRNA serving as primary cytoplasmic reservoirs for both Pi and Mg^2+^.^51–54^ Previous work by Bruna et al. demonstrated that bacteria suppress Pi uptake under Mg^2+^ limitation to preserve cellular function.^52^ Herein, our study extends this understanding by showing that Pi scarcity itself actively triggers Mg^2+^ release from *E. coli* cells. This observation underscores the tightly coupled nature of Pi and Mg^2+^ homeostasis. Our comprehensive proteomic analysis further supports this, revealing that proteins involved in translation, ribosomal structure, and biogenesis were among the most significantly downregulated groups during Pi depletion (Figures 1A and S3). Concurrently, we observed a steady decline in cellular ATP levels under Pi-limited conditions (Figure S19), suggesting a profound arrest in ribosome biogenesis and protein synthesis that likely contributes to the observed Mg^2+^ release and cellular loss.

In *Enterobacteriaceae*, Mg^2+^ homeostasis is primarily governed by the PhoPQ two-component system, which is well-characterized in *S.* Typhimurium to be activated under low extracellular Mg^2+^ conditions.^55,56^ Given the observed Mg^2+^ release during Pi-depletion, we investigated the involvement of this classical Mg^2+^-sensing PhoPQ system. Our results showed that despite cellular Mg^2+^ loss during Pi depletion, the transcription of key PhoPQ regulon genes, such as *phoQ* and *mgtA*, was not upregulated (Figure S20). This indicates that while cellular Mg*^2+^*levels are perturbed, they do not drop sufficiently to activate the canonical PhoPQ response. Instead, our data compellingly demonstrate that under acute Pi depletion, *Enterobacteriaceae* preferentially engage an alternative, Fe^3+^-mediated membrane remodeling pathway. This pathway activates the *arn* operon and subsequent L-Ara4N modification independently of the classical Mg^2+^-sensing PhoPQ system, highlighting a distinct adaptive strategy.

This Fe^3+^-mediated lipid A modification strategy in *Enterobacteriaceae* offers a stark contrast to Pi depletion-induced polymyxin tolerance mechanisms described in other Gram-negative bacteria, notably *P. aeruginosa*. In *P. aeruginosa*, Pi depletion primarily activates the PhoBR regulon, orchestrating the biosynthesis of phosphorus-free glycolipids (MGDG and GADG) to replace negatively charged phospholipids and maintain membrane integrity (Figure 8B).^11^ While *P. aeruginosa* possesses a PmrAB homolog, its PmrB sensor kinase lacks the critical Fe^3+^-sensing “ExxE” motifs found in *Enterobacteriaceae* PmrB, precluding direct Fe^3+^ response. Instead, PmrAB activation in *P. aeruginosa* is typically coupled to the PhoPQ system, which senses low extracellular Mg^2+^. Thus, while *P. aeruginosa* employs PhoBR-regulated phospholipid remodeling as its primary Pi depletion response, *Enterobacteriaceae*, largely lacking these glycolipid pathways, have evolved to utilize a secondary, Fe^3+^-sensing mechanism. The physiological basis for this divergence remains unclear. One possible contributing factor is the difference in central carbon metabolism between *Enterobacteriaceae* and *Pseudomonads*.^57^ *Enterobacteriaceae* primarily catabolize glucose through the Embden–Meyerhof–Parnas (EMP) pathway, whereas *P. aeruginosa* lacks phosphofructokinase and consequently relies on the Entner–Doudoroff (ED) pathway.^58^ These pathways differ in their organization of phosphorylated carbon intermediates, ATP yield, and redox output; therefore, Pi limitation may impose distinct metabolic constraints on the two bacterial groups. Such differences could influence the availability of precursors for lipid synthesis or the coupling of Pi stress to cell-envelope adaptation, potentially contributing to the distinct responses observed: PhoBR-associated phospholipid-to-glycolipid remodeling in *P. aeruginosa* and activation of the Mg–Fe–PmrAB–Arn pathway in *Enterobacteriaceae*. However, this hypothesis remains speculative and warrants further investigation. In *Enterobacteriaceae*, PmrAB senses Fe³⁺ through the Fe³⁺-coordinating ExxE motifs of PmrB and activates the Arn pathway, resulting in L-Ara4N modification of lipid A as an adaptive response to Pi depletion.^49^ By contrast, Pi depletion in *P. aeruginosa* also induces LpxO-mediated 2-hydroxylation of the secondary laurate chains of lipid A. However, this modification does not contribute to polymyxin tolerance and is regulated independently of PhoBR, PhoPQ, and PmrAB, further highlighting the distinct physiological responses of these organisms to Pi limitation.^50^

The biosynthesis of L-Ara4N and its transfer to lipid A are mediated by Ugd and ArnABCDEFT proteins, encoded by the *ugd* gene and the *arn* operon, respectively.^59–62^ Our data reveals a sophisticated regulatory separation: during Pi depletion, *arn* transcription is tightly controlled by the PmrAB system, while ugd expression is not PmrAB-dependent and exhibits partial regulation by the PhoBR system (Figure S6). This bifunctional control is not unexpected, as *ugd* encodes UDP-glucose dehydrogenase, an enzyme with dual roles in L-Ara4N biosynthesis and in the production of exopolysaccharides like colanic acid.^63,64^ Therefore, while PmrAB activation of *arn* is absolutely critical for L-Ara4N modification of lipid A during Pi limitation, *ugd* regulation reflects its broader involvement in diverse cellular metabolic and stress responses.

While preparing this manuscript, a recent study by Baijal et al. (2024) reported that the *ppk* gene modulates *arn* operon upregulation under combined nutrient limitations (transfer from rich LB to low-Pi, no-amino-acid MOPS medium) in *E. coli.*^65^ Inspired by this, we investigated the role of *ppk* in polymyxin tolerance specifically under Pi depletion, our primary focus. We found that *ppk* deletion did not affect *arn* or *ugd* promoter activity under sole Pi-depleted conditions (Figure S21), indicating its dispensability when Pi is the only limiting factor. To reconcile these findings, we replicated the Baijal et al. experimental conditions, confirming *ppk*-dependent *arn* upregulation under combined Pi and amino acid scarcity (Figure S22A). Our subsequent experiments, systematically disentangling these stresses, revealed that while both Pi and amino acid depletion induce *arn* upregulation, Pi depletion is the dominant driver, and *ppk*-dependent regulation emerges only under combined Pi and amino acid scarcity (Figure S22B). These results clarify that while *ppk* influences *arn* expression under specific multifactorial stresses, our identified PmrAB-arn pathway is independently and dominantly activated by Pi depletion alone, providing precise mechanistic insight into this fundamental environmental stressor.

In both natural habitats and host-associated niches, Pi scarcity imposes dual challenges: nutrient acquisition and defense against antimicrobial threats. The Pi-Mg-Fe-PmrAB regulatory circuit we have uncovered likely represents a sophisticated adaptive strategy for *Enterobacteriaceae*, facilitating their survival in highly heterogeneous environments with fluctuating Pi levels, such as during host colonization, transmission, or environmental persistence.^66^ This PmrAB-mediated membrane remodeling not only confers protection to polymyxins but may also enable bacteria to preserve membrane integrity and withstand other cationic antimicrobial peptides (CAMPs) produced by competing microbes or host defenses.

In summary, our findings elucidate a conserved yet distinct regulatory circuit in *Enterobacteriaceae* that intricately links Pi and Mg^2+^ homeostasis to Fe^3+^-mediated lipid A modification, driving stress-induced antibiotic tolerance. This mechanism is notably distinct from those characterized in *P. aeruginosa* or *A. baumannii*. Crucially, we demonstrate that targeting this circuit effectively mitigates stress-induced tolerance. Given that polymyxins remain indispensable as last-line antibiotics for treating multidrug-resistant Gram-negative bacterial infections, our study not only offers critical insights into the fundamental molecular mechanisms underlying stress-induced tolerance but also highlights a promising avenue for informing novel tolerance management strategies.

## Materials and methods

### Bacterial strains and growth conditions

Strains, plasmids, and primers used in this study are listed in the S2 and S3 Tables. Bacteria were cultured in lysogeny broth (LB) broth/plates or MOPS minimal medium. MOPS medium was freshly prepared from 1× MOPS stock by supplementing 0.1 % (w/v) glucose and 1.32 mM K_2_HPO_4._ Acute Pi depletion was applied by transferring bacteria culture into the non-Pi MOPS medium. 1× MOPS consists of 40 mM MOPS, 4 mM Tricine, 10 μM FeSO_4_·7H_2_O, 9.5 mM NH_4_Cl, 0.276 mM K_2_SO_4_, 0.5 μM CaCl_2_·2H_2_O, 525 μM MgCl_2_, 50 mM NaCl, 0.003 μM (NH_4_)_6_Mo_7_O_24_, 0.4 μM H_3_BO_3_, 0.03 μM CoCl_2_, 0.01 μM CuSO_4_, 0.08 μM MnCl_2_, and 0.01 μM ZnSO_4_. Antibiotics applied to select and maintain the construct were chloramphenicol (25 μg/ml), kanamycin (20 μg/ml) and ampicillin (100 μg/ml).

### Lipid A extraction and ESI-MS analysis

Lipid A was extracted from designated *E. coli* cells using Bligh-Dyer method as described with minor modifications.^67^ *E. coli* MG1655 cultured in MOPS medium to OD_600_ of 0.5 was harvested as cells under Pi sufficient condition. The same volume of cells was washed with Pi depletion medium for 3 times and resuspended in Pi depletion medium and shake for 3 hrs at 37°C as cells subjected to Pi depletion. 200 ml of each type of cell pellets with three biological replicates were harvested by centrifugation at 4500 rpm for 30 min and washed with H_2_O. The cell pellets were suspended in 76 ml of a single-phase Bligh-Dyer mixture (chloroform: methanol: water=1:2:0.8; v/v/v) and stirred at room temperature for 1 h. Insoluble debris was collected by centrifugation at 4500 rpm for 30 min, washed with 60 ml single-phase Bligh-Dyer mixture, and suspended in 27 ml of 12.5 mM sodium acetate (pH 4.5). The mixture was heated at 100°C for 30 min and cooled to room temperature. The suspension was mixed with 30 ml chloroform and 30 ml methanol. Following vigorously shaken for extraction of lipid A and centrifugation for 30 min, the lower phase containing lipid A was collected and dried with a rotary evaporator.

All the mass spectra of lipid A samples were acquired using Waters SYNAPT quadrupole time-of-flight mass spectrometer (MS) equipped with an electrospray ionization source^68^. Lipid A samples were dissolved in a mixture of chloroform and methanol (4: 1; v/v) and subjected to ESI/MS in the negative ion mode. Data acquisition and analysis were performed using MASS LYNX V4.1 software, Waters, Milford, MA, USA.

### ICP-MS and DFOM treatment

Total cellular metal content was quantified using inductively coupled plasma mass spectrometry (ICP-MS) following previous description with minor modifications^69^. *E. coli* cells equivalent to OD_600_ as 5 were harvested by centrifugation at 4500 rpm for 15 mins and washed three times with ice-cold PBS buffer. Cell pellets were subjected to acid digestion with 100 μl 65.0 % HNO_3_ (Fluka) at 65°C for 16hrs. Samples were diluted to a final volume of 15 ml with 1% HNO_3_ and subjected to ICP-MS detection on Agilent 7900 system (Agilent Technologies, CA). Samples were further diluted if the signals exceeded the liner range of standard curve. The standard curves of tested metals were generated from multi-element calibration standard-2A for ICP-MS (Agilent). Metal contents in bacteria were calculated based on the standard curve and results were normalized to original cell number.

Deferoxamine mesylate salt (sigma) were applied to examine the degree of iron association to *E. coli* cell surface. *E. coli* cells equivalent to OD_600_ as 5 were harvested by centrifugation at 4500 rpm for 15 mins. Cell pellets were resuspended in HEPES buffered saline with and without 50μM DFOM and incubated in 37°C for 30 mins with 220rpm agitation. Total cellular metal content of these treated cells was quantified using inductively coupled plasma mass spectrometry (ICP-MS) as described above.

### OM feature monitoring using PI-BactD

OM feature of *E. coli* was monitored by an OM-sensitive probe PI-BactD following previous description with minor modifications.^48^ *E. coli* cells grown under designated conditions were harvested by centrifugation at 13000 rpm for 2 mins and washed with PBS buffer (10 mM, pH 7.2) twice. Cell pellets were resuspended in PBS buffer to an OD_600_ of 0.5. Fluorescence probe PI-BactD (gift from prof. Xu Z, Dalian) was added into 1mL cell suspension to a final concentration of 2 mM and incubated at room temperature for 1 min. Output signals of two channels: fluorescence increase (ΔS/S_0_) and fluorescence ratio metric changes (I_482_/I_375_) were recorded to plot the 2D graph (Thermo Scientific Varioskan^®^ Flash spectral scanning multimode reader). Six replicates were conducted for each strain and treatment.

### Construction of Ppromoter-*lacZ* (P*arn-lacZ* as example) fusion in the single-copy plasmid pNN387

A DNA fragment corresponding to the −200 to −1 region relative to the ATG start codon of *arnB* was amplified using *E. coli* MG1655 genomic DNA as the template and primers specific to this fragment flanked with NotI and HindIII restriction sites at the 5’- and 3’-end, respectively. Following digestion with NotI and HindIII and purification using the MiniBEST agarose gel DNA extraction kit (TaKaRa), the fragment was ligated into the plasmid pNN387 which was treated with the same restriction enzymes using the quick ligation kit (NEB). The resulting construct was selected on LB agar plates supplemented with chloramphenicol (25 μg/ml) and was verified by DNA sequencing (BGI, Shenzhen).

### Beta-galactosidase activity assay

β-gal activity assay was performed based on a previous description with modifications.^69^ Cells were grown under designated conditions until OD_600_ reached 0.4. 10 μg/ml tetracycline was added to stop cell growth and protein synthesis, and cells were left on ice until being assayed. 100μl of cell culture and 400 μl of Z buffer (60 mM Na_2_HPO_4_, 40 mM NaH_2_PO_4_, 10 mM KCl, 1 mM MgCl_2_, 50 mM β-mercaptoethanol, pH 7.0, 4 °C) were mixed. One drop of 0.1% SDS and two drops of chloroform were added to lyse the cells. The reactions were initiated by adding 100 μl of 4 mg/ml ortho-nitrophenyl-β-galactoside into cell suspension and were incubated at 28 °C. Color development in the mixtures was monitored. Finally, 250μl of 1M Na_2_CO_3_ was added to stop the reaction. Color of the mixtures were measured using spectrophotometer at OD_420_ and OD_550_. The expression value in Miller units was calculated, and results are presented as the mean of three biological replicates.

### RNA extraction and RT-qPCR

Bacterial cells grown under designated conditions was subjected to RNA extraction when the OD_600_ reached 0.4. Cells were harvested by centrifugation at 4,500 rpm for 10 min at 4 °C. After removing the supernatant, the cell pellet was stored at −80 °C to aid cell lysis. Total RNA was extracted using the MiniBEST Universal RNA Extraction Kit (Takara, Japan) according to the manufacturer’s instruction. Reverse transcription (RT) was performed using PrimeScript^TM^ RT reagent Kit (Takara, Japan). Quantitative PCR was performed using specific primers and the TB Green® Premix Ex Taq™ (Takara, Japan) in a 20 μL reaction system. The reaction was performed in the ABI StepOnePlus real time PCR system with *rrsB* as reference gene to normalize the relative expression of the target genes. The results were presented as fold change expression of the target genes, and results were presented as the mean of three biological isolates.

### Construction of clean gene deletion, point mutation and tag insertion in chromosome of *E. coli* MG1655

Gene deletion, mutation, and tag insertion in chromosomal were achieved employing a CRISPR/Cas9-mediated approach developed by Zhao et al. with modifications.^70^ For gene deletion (deletion of *lacZ* as example), *E. coli* MG1655 was electrotransformated with pCAGO plasmid which contains both SpCas9 and λ-RED to obtain MG1655-pCAGO. A single colony of the resulting transformant was inoculated into LB medium supplemented with ampicillin (100 mg/ml) and grew overnight at 30°C. 100 μl of the overnight culture was inoculated into 10 ml fresh culture medium and cells were grown at 30°C with IPTG (0.1 mM) induction to OD600 as 0.6. The cells were collected and washed with cold ddH_2_O for three times, generating competent cells for electroporation. An aliquot comprising 100 μl competent cells was mixed with 800 ng of the editing-cassette which encompasses 40 bp of sequence among the CDS of *lacZ* gene followed by the chloramphenicol resistant gene, the N20PAM sequence (GTCCATCGAACCGAAGTAAGG), 40 bp upstream-homologous arm of *lacZ* gene, and 40bp downstream-homologous arm of *lacZ* gene, in a 2 mm Gene Pulser cuvette (Bio-Rad) and was subjected to electroporation at 2.3 kV. Following recovering the transformants by cultivation in LB medium for 2 hrs at 30°C, the mixture was spread onto LB agar plate supplemented with ampicillin (100mg/ml), chloramphenicol (25 mg/ml), and 1% glucose (to avoid the background expression of the Cas9 protein). Single colonies were collected and verified by colony PCR to obtain an authentic clone in which the editing-cassette was embedded in chromosomal. The confirmed colony was inoculated into LB medium supplemented with ampicillin (100 mg/ml), IPTG (0.1 mM), and L-arabinose (0.2%) at 30°C overnight to enable expression of SpCas9 and the λ-RED recombinase. The cells were then streaked on LB agar plates supplemented with ampicillin (100mg/ml). Following colony PCR, desired colonies containing deletion of the *lacZ* gene were verified by DNA sequencing (BGI, Shenzhen). For point mutation, (mutation of *pmrB* E36 to A as example) *E. coli* MG1655 was conducted following the same procedures except for the editing cassette sequence which encompasses 40 bp of upstream-homologous arm of GAA (Glu36) followed by the chloramphenicol resistant gene, the N20PAM sequence, the 40 bp upstream-homologous arm of GAA (Glu36), GCA (Ala) and 40 bp downstream-homologous arm of GAA (Glu36). For tag insertion, (insertion 3xFLAG to C-terminal of *arnB* gene as example) *E. coli* MG1655 was conducted following the same procedures except for the editing cassette sequence which encompasses 40 bp of upstream-homologous arm of stop codon of *arnB* gene followed by the chloramphenicol resistant gene, the N20PAM sequence, the 40 bp upstream-homologous arm of stop codon of *arnB* gene, tag sequence and 40 bp downstream-homologous arm of stop codon of *arnB* gene (include stop codon). After obtaining all desired genome modifications, the editing plasmid was cured by growing the resulting cells at 42°C for 48 hrs.

### Construction of pET28A-pBAD-*pmrA* for PmrA protein expression

A DNA fragment encoding the PmrA protein was amplified using genomic DNA of *E. coli* MG1655 as the template and primers specific to this fragment flanked with NcoI and XhoI restriction sites at the 5’- and 3’-end, respectively. After digestion with NcoI and XhoI, and purification using the MiniBEST agarose gel DNA extraction kit (TaKaRa), the purified fragment was ligated into the plasmid pET28a-pBAD and pET28a-pBAD-3xFLAG ^42^ which was treated with the same restriction enzymes using the quick ligation kit (NEB). The resulting construct was selected on LB agar plates supplemented with kanamycin (20 μg/ml) and was verified by DNA sequencing (BGI, Shenzhen). All plasmids constructed are listed in the supplementary S3 Table.

### Immunoblot analysis

Protein levels of ArnB and PmrA were analyzed by SDS-PAGE following Western Blot. 1ml of cells with OD_600_ of 0.5 was collected and resuspended in 200 μl sample buffer (13 mM Tris, 0.2% SDS, 5% glycerol, 0.004% bromophenol blue, pH 6.8) before being subjected to cell lysis by sonication. Following centrifugation at 13000 rpm for 10 min at 4°C, 20 μl of the sample was separated on SDS-PAGE gels (TGX FastCast Acrylamide Kit, 10%, BIO-RAD). Proteins were then transferred onto a polyvinylidene difluoride (PVDF) membrane (BIO-RAD) using a Trans-Blot SD Semi-Dry Transfer Cell (BioRad) at constant voltage (25 V) for 7 min. Membranes were blocked for 1 h with 5% non-fat milk solution in TBST (20 mM Tris, 150 mM NaCl, 0.1% Tween-20) at room temperature followed by incubation with 1:10,000 primary antibody (monoclonal or polyclonal anti-His antibody) in 1% non-fat milk in TBST at room temperature with shaking for 1 hour. After washing with TBST, the membrane was incubated with the secondary antibody (Goat anti-mouse IgG HRP conjugate) in 1% non-fat milk in TBST at room temperature with shaking for 1 hour. Chemiluminescence of protein bands was detected by ECL Plus Western Blotting Detection Reagents (GE Healthcare, USA) for 2 min and then subjected to the imaging system (UVITEC Cambridge) to record signals.

### Phos-tag SDS-PAGE

In vivo phosphorylation of PmrA was examined using Phos-Tag SDS-PAGE following our previous description with minor modifications.^42^ *E. coli* MG1655Δ*pmrA* cell harboring pET28a-pBAD-*pmrA*-3×FLAG (AY7068) was inoculated into MOPS minimum medium supplemented with kanamycin (20 μg/ml) and grew overnight at 37°C. 100 μl of the overnight culture was inoculated into 5 ml fresh MOPS medium and grew at 37°C for 4 hrs. Expression of PmrA was induced with 1mM L-arabinose for 1 hr. Cells were collected by centrifugation at 13000 rpm and were washed with MOPS medium supplemented with 0.1, 0.5, or 1 mM FeSO_4_ or Pi-MOPS medium twice and shake for 1 hr at 37°C in the corresponding medium. 1ml of cells with OD_600_ of 0.5 was collected for the subsequent analysis. Cell pellet was washed with cold PBS twice and was resuspended in 200 μl sample buffer (13 mM Tris, 0.2% SDS, 5% glycerol, 0.004% bromophenol blue, pH 6.8) before being subjected to cell lysis by sonication. Following centrifugation at 13000 rpm for 10 min at 4°C, 20 μl of the sample was loaded onto a 10% Tris-Gly polyacrylamide gel containing 25 mM Phos-tag Acrylamide (Wako) and 50 mM MnCl_2_. Samples were separated by electrophoresis for 2 hrs at 4°C, following washing with 50 ml transfer buffer containing 10 mM EDTA for 20 min to remove residual Mn^2+^. Proteins on the gel were transferred onto a PVDF membrane. The membrane was blocked with 5% non-fat milk in TBST buffer and was incubated subsequently with monoclonal ANTI-FLAG^®^ M2 antibody (Sigma) and goat anti-mouse IgG (H+L)-HRP conjugate (Bio-rad). To detect FLAG-tagged protein bands, the membrane was submerged in the ECL Plus Western Blotting Detection Reagents (GE Healthcare) for 2 min and then subjected to the imaging system (UVITEC Cambridge) to record signals.

### Amber codon mutation of selected CouA incorporation sites in pET28A-pBAD-*pmrA*

Selective amino acid located in the DNA binding domain of PmrA was subjected to replacement by CouA, each of these CouA incorporation sites were mutated to a TAG amber codon by site-directed mutagenesis PCR. PCR was conducted using a pair of primers for the desired mutagenesis and pET28A-pBAD-*pmrA* plasmid as the template. PCR product was purified using MiniBEST agarose gel DNA extraction kit (TaKaRa). Following DpnI digestion of the template for 16 hrs, the PCR product was transformed into *E. coli* DH5α. Transformants were recovered on LB agar plates supplemented with kanamycin (20 μg/ml) at 37°C overnight. Colonies recovered were verified by DNA sequencing (BGI, Shenzhen).

### In vivo PmrACouA - promoterSYTO 9 binding assay

Binding of PmrA_CouA_ and promoter_SYTO 9_ was monitored by FRET assay following our previous description with minor modifications^42^. Plasmid pET28a-pBAD-*pmrA*-G137TAG was co-transformed with pEVOL-CouRS into *E. coli* MG1655 *ΔpmrA* strain, resulting in AY7065 constructs. A fresh single colony of AY7065 was inoculated into MOPS medium supplemented with kanamycin (20 μg/ml) and chloramphenicol (25 μg/ml) and grew overnight at 37°C. 75 μl of the overnight culture was inoculated to 7.5 ml fresh medium to subculture the bacteria. Following culturing at 37°C for 4 hrs, CouA incorporation and expression of PmrA-G137CouA was induced mildly by supplementing 0.9% arabinose and 1mM CouA at 37°C for 5 hrs. Approximately 3.5×10^10^ cells were harvested and washed twice with fresh MOPS medium. Cells were resuspended in MOPS medium as control, or MOPS medium supplemented with 0.5 mM FeSO_4_, or Pi-MOPS medium and shake for 1 hr at 37°C. After centrifugation at 4500 rpm for 15 min at 4°C, cell pellet was resuspended in 600 μl of PBS. 85 μl of the suspension was added in each of 6 wells (3 wells for control and 3 wells for experimental group) on a 96-well black plate preloaded with 15 μl of 50 μM SYTO 9 and was incubated for 15 min at room temperature in the dark before being subjected to fluorescence recording (Thermo Scientific Varioskan® Flash spectral scanning multimode reader).

### Comparative proteomic analysis

*E. coli* MG1655 cultured in MOPS medium to OD_600_ of 0.5 was harvested as cells under Pi sufficient condition. The same volume of cells was washed with Pi depletion medium for 3times and resuspended in Pi depletion medium and shake for 3 hrs at 37°C as cells subjected to Pi depletion. Each type of cell pellets with three biological replicates were lysed by resuspending in 20 μl BugBuster reagent (Novagen, USA) (containing 25 mg/ml lysozyme, 1 unit DNase I) at room temperature for 30 mins following centrifugation at 13000 rpm at 4°C for 20 min. Supernatant was transferred to a clean microcentrifuge tube and mixed with 4×SDS sample buffer (40% glycerol, 8% SDS, 240 mM Tris/HCl pH 6.8, 0.04% bromophenol blue) supplemented with 5% β-mercaptoethanol. Sample were incubated at 55 °C for 25 min. 20 µL of each sample were loaded onto a polyacrylamide gel composed of a 10% resolving (lower) gel and a 6% stacking (upper) gel following staining with Coomassie blue staining and fractionated into eight fractions. Gel slices were cut into 1 mm^3^ cubes and destained with 50% acetonitrile (ACN) in 50 mm triethyl ammonium bicarbonate (TEAB) and dehydrated with pure CAN and dried in a vacuum concentrator. Sequencing grade modified trypsin (Promega) diluted by an ice-cold buffer containing 50 mM NH4HCO3 and 10% (v/v) ACN was added to cover the dehydrated gel pieces on ice. After 30 min, more trypsin-containing buffer was added to cover the expanded gel pieces followed by overnight incubation at 37 °C. Finally, the resulting tryptic peptides were extracted from the gel by equilibrating the samples with 50% ACN and 5% formic acid (FA) for 20 min in a 37 °C shaker. The extraction was repeated once, and all fractions were combined and vacuum dried. The protein digest was resuspended in 100 μl of 100 mm TEAB, and then 4 μl of 0.6 m sodium cyanoborohydride (NaBH_3_CN) were added. Peptides from PI+ and Pi-were labeled with 4 μl of 4% formaldehyde (CH_2_O) and deuterated formaldehyde (CD_2_O), respectively. The reaction mixture was vortexed and incubated at room temperature for 1 h. The reaction was quenched by sequential addition of 16 μl of 1% (v/v) ammonia solution and 8 μl of formic acid. Finally, light- and heavy-labeled peptides were mixed, and vacuum dried immediately before further mass spectrometric analyses. Extracted peptides were analyzed by Nanoflow LC-MS/MS analyses using the EASY-nLC 1000 System (Thermo Scientific, USA) and LTQ Velos (Thermo Scientific, USA).

Data processing was performed following a previous description^71^. For protein identification, MS/MS spectra were extracted from RAW files using MaxQuant (http://maxquant.org/, version 1.5.3.30). MASCOT (Matrix Science, version 2.3.02) was used to perform database search against an *E. coli* database (strain MG1655, with sequence files downloaded from UniProt in FASTA format, 4,358 entries. Average mass was selected with precursor mass tolerance of 1.5 Da and fragment mass tolerance of 0.8 Da. Methionine oxidation was set as a dynamic modification. The resulting assignments were filtered to achieve a false-discovery rate (FDR) of <1% at both the peptide and protein levels using the target-decoy method. Relative abundances of protein were represented by spectral counts, which are the number of repeated identification of peptides during the entire analysis.

To identify proteins that were differentially expressed in the sets of samples, the spectral counts data were subjected to statistic by DESeq package (Anders and Huber 2010) in R (Team 2014). Briefly, to eliminate the variance caused by sample loading and machine reading depth, the total counts data of two biological conditions was normalized to the same size. The following statistics were performed: *foldChange* (fold change from condition A to B), *pval* (p value for the statistical significance of this change), and *padj* (p value adjusted for multiple testing with the Benjamini-Hochberg procedure, which controls false discovery rate). Data set with *padj* <0.05 were considered to be significant (Anders and Huber 2010), and corresponding proteins were listed in S1 Table.

### Polymyxin challenging assay

Challenging *E. coli* cells with designated concentration of polymyxin B (Sigma. USA) were performed as described with modifications.^72^ *E. coli* cells grown under designated conditions were harvested by centrifugation at 13000 rpm for 1min. Cell pellets were re-suspended in MOPS medium to a final concentration of 10^8^ cells/ml. The cell culture was treated with polymyxin B with designated concentration for 1-3 hours at 37 °C without agitation. Cell culture prior to and following polymyxin treatment was plated on LB plate, CFU was counted after 12-16 hours incubation at 37°C. Survival of polymyxin challenging was determined by the survival rate of *E. coli* cells after drug treatment. Results are presented as the mean of three biological replicates.

### Phylogenetic analysis of EcoPmrB homologs

The PmrB sequences from *E. coli* K-12 MG1655 were used as a query against γ-proteobacteria via BLASTP searches in the UniProt database, with an E-threshold of 10 and the BLOSUM62 substitution matrix. The top 250 retrieved sequences were aligned using the MUSCLE algorithm implemented in MEGA12 software. A phylogenetic tree of EcoPmrB homologous was constructed using the Maximum Likelihood method, with branch support assessed via adaptive bootstrap analysis using 562 bootstrap replicates in MEGA12. The JTT model was applied as the evolutionary model for tree construction.

### Statistical analysis

Statistical analysis was performed using GraphPad Prism. Group comparisons between two groups were performed using two-tailed Student’s *t* test. Group comparisons between multiple groups were performed using one-way ANOVA or two-way ANOVA with post hoc tests.

## Acknowledgments

We thank Ms. Yining Chen in Prof. A. Yan’s lab for proofreading the manuscript. We thank Prof. Changhao Bi (Tianjin Institute of Industrial Biotechnology, Chinese Academy of Sciences) for the generous gift of pCAGO plasmid. We thank Prof. Zhaochao Xu and Prof. Lu Miao (CAS Key Laboratory of Separation Science for Analytical Chemistry, Dalian Institute of Chemical Physics, Chinese Academy of Sciences) for the generous gift of OM-sensitive probe PI-BactD. We thank Prof. Patrick C. Y. Woo (Department of Microbiology, University of Hong Kong) for the generous gift of clinical *E. coli* strains. We thank Prof. Patricia J. Kiley (Department of Biomolecular Chemistry, University of Wisconsin-Madison) for sharing the Δ*bfr* Δ*dps* Δ*ftnA* strain and for stimulating discussions)

## Author contributions

**Conceptualization:** Guangming Zhang, Ziqing Deng, Aixin Yan.

**Investigation** Guangming Zhang, Ziqing Deng, Jiezhang Jiang, Minji Wang, Xiaoyuan Wang, Xiaoyun Liu, Aixin Yan.

**Formal Analysis** Guangming Zhang, Ziqing Deng, Xiaoyuan Wang, Xiaoyun Liu, Aixin Yan.

**Funding Acquisition:** Xiaoyun Liu, Aixin Yan.

**Resources:** Xiaoyuan Wang, Xiaoyun Liu, Aixin Yan.

**Supervision:** Xiaoyuan Wang, Xiaoyun Liu, Aixin Yan.

**Visualization:** Guangming Zhang.

**Writing – original draft:** Guangming Zhang, Aixin Yan.

**Writing – review & editing:** Guangming Zhang, Xiaoyun Liu, Aixin Yan.

## Data and materials availability

All data are available in the main text or the supplementary materials.

## Supporting information

**Figure S1.**
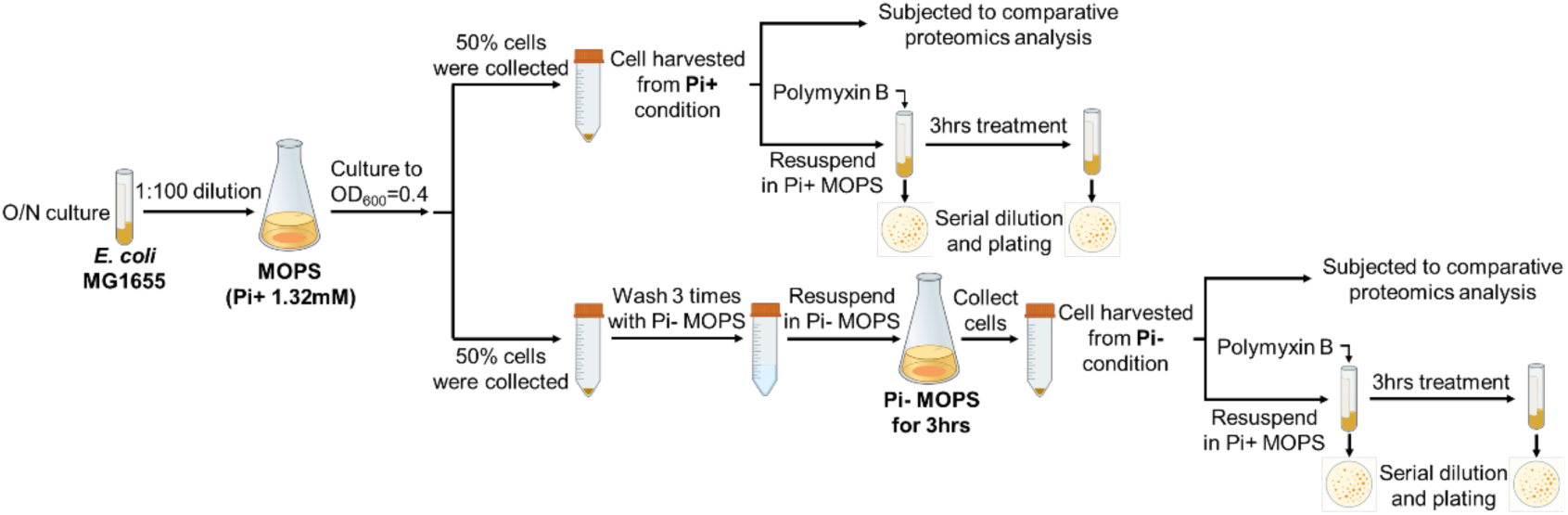
Workflow of acute Pi depletion treatment and following proteomic sample preparation and polymyxin B challenging assay. Bacteria were cultured to an OD_600_ of 0.4 in standard MOPS synthetic medium containing 1.32 mM inorganic phosphate (Pi), representing Pi-sufficient conditions. For Pi-depletion conditions, bacteria were first grown to OD_600_ = 0.4 in the same medium, then transferred to Pi-depleted MOPS medium (0 mM Pi). For comparative proteomic analysis, the collected bacterial cells were subjected to sample preparation and proteomic analysis. For the polymyxin challenging assay, the collected bacterial cells were resuspended in standard MOPS medium and treated with the designated concentration of polymyxin B. Cell viability was assessed at the beginning and end of treatment by serial dilution and plating.

**Figure S2.**
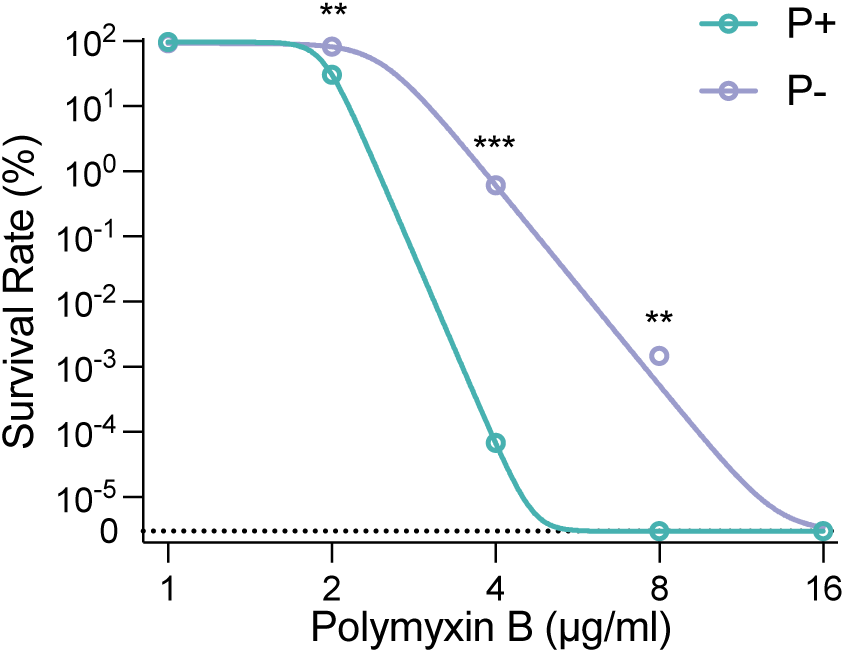
Killing of *E. coli* MG1655 with different concentration of polymyxin. **B**. Survival rates for *E. coli* MG1655 cells exposed 1-hour to polymyxin B (1-16 μg/ml) following acute Pi-starvation treatment (Pi-, 0 mM) compared to the group without treatment (Pi+, 1.32 mM). The dotted line labelled “0” denotes the limit of detection; values below this threshold were not detectable by colony counting. Error bars represent the standard deviation of at least three independent experiments. Statistical significance was determined using unpaired two-tailed Student’s t-test. Significance levels are indicated as follows: \**p* < 0.05; \*\**p* < 0.01; \*\*\**p* < 0.001.

**Figure S3.**
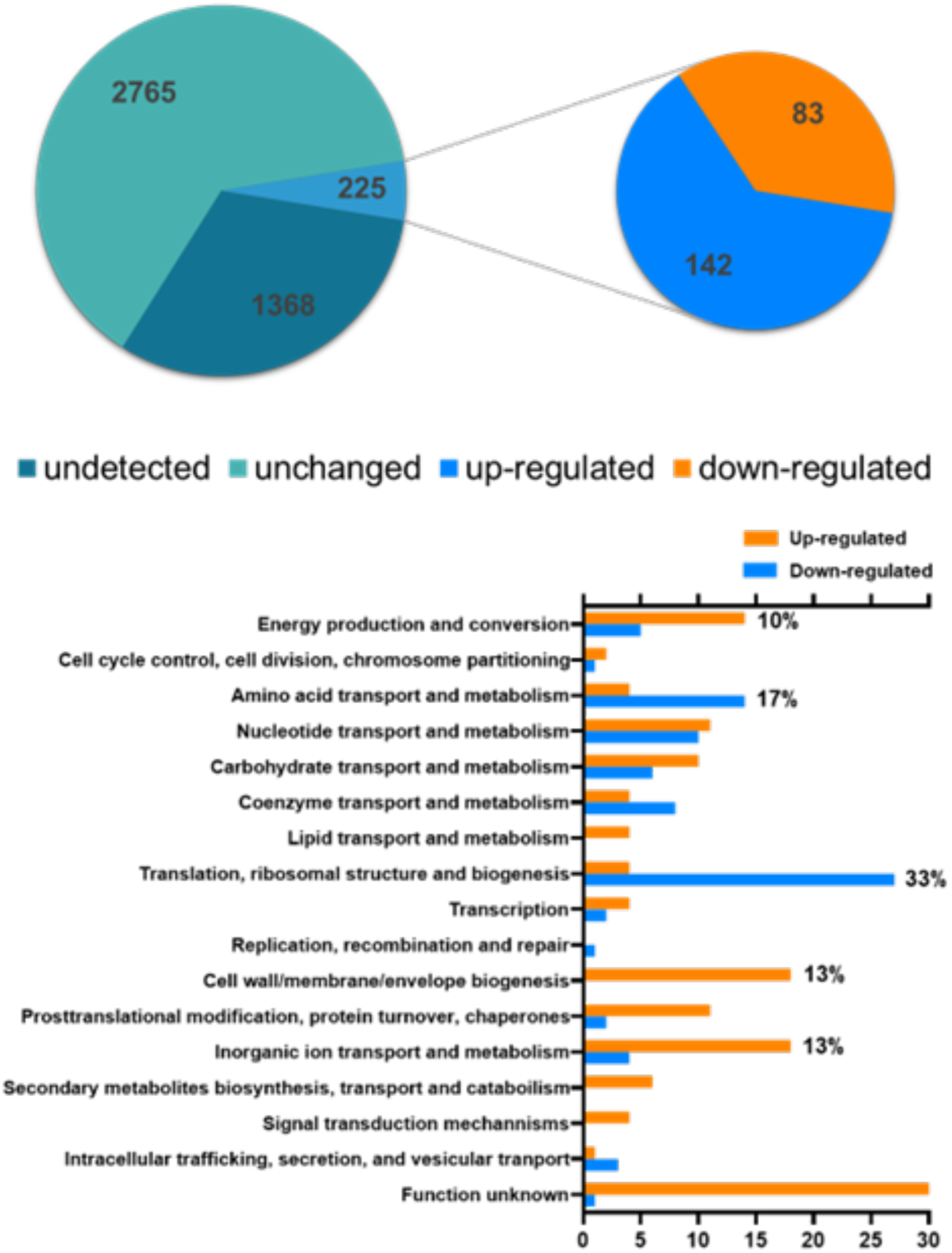
Quantitative proteomic analysis of *E. coli* MG1655 in response to acute Pi depletion. Pie chart of total and differentially expressed proteins detected by LC-MS/MS analysis in *E. coli* MG1655 cells subjected to Pi depletion for 3 hours, relative to cells under Pi sufficient conditions. Red and blue colors represent significantly upregulated and downregulated proteins, respectively. Clusters of Orthologous Groups (COG) functional categories of differentially expressed proteins in *E. coli* MG1655 in response to Pi depletion. Orange and blue colors represent significantly upregulated and downregulated proteins, respectively.

**Figure S4.**
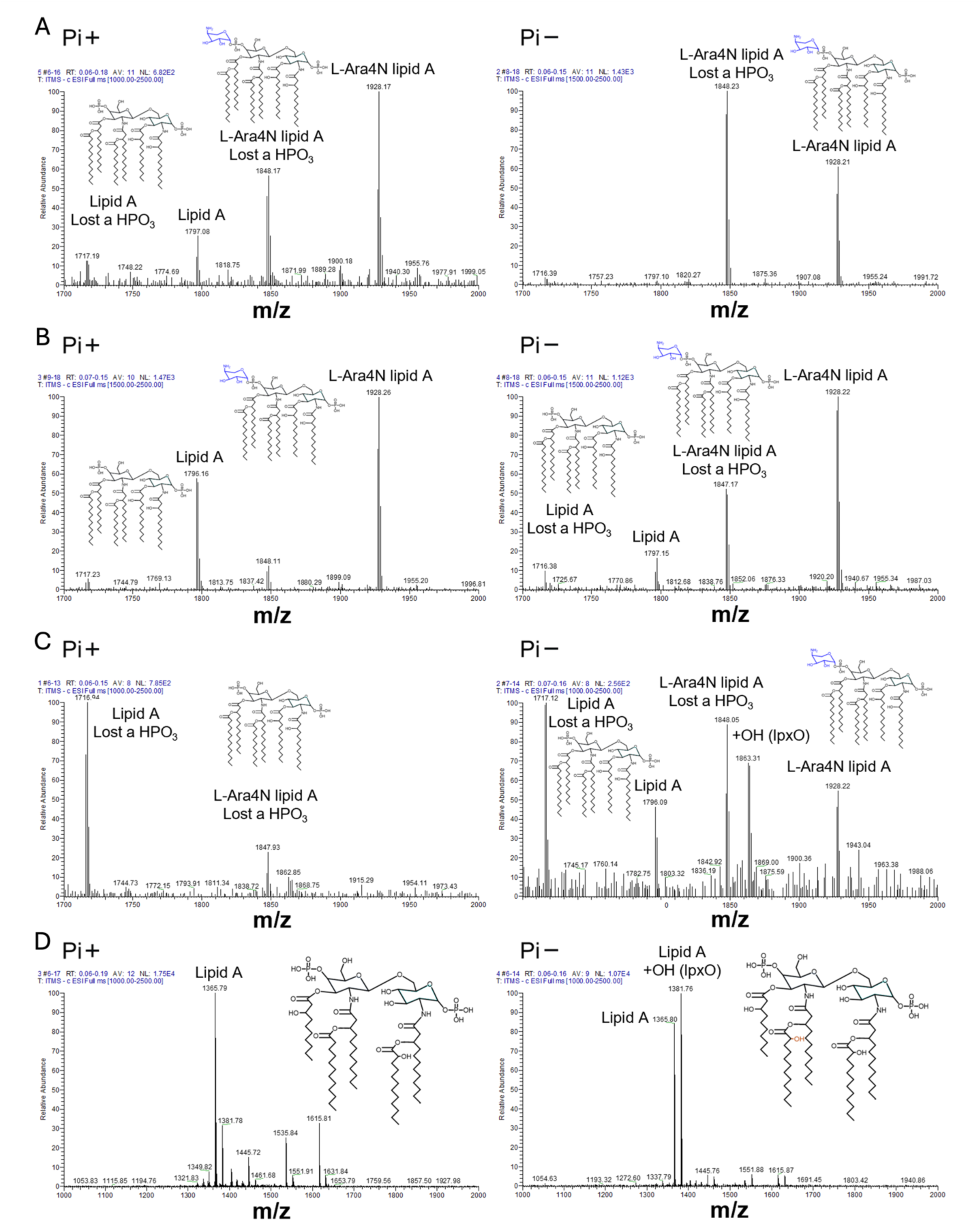
The ratio of L-Ara-4N modified Lipid A molecule increases in *E. coli* O157, *E. coli* UTI-3, *S.* Typhimurium ATCC14028 and *P. aeruginosa* PAO1 cells subjected to Pi depletion. (**A**) ESI-MS spectra of lipid A molecules extracted from *E. coli* O157 cells following acute Pi-starvation treatment (Pi-, 0 mM) (right) compared to the group without treatment (Pi+, 1.32 mM) (left). Peaks correspond to native lipid A lost a HPO_3_ (1718), native lipid A (1798), L-Ara-4N modified lipid A lost a HPO_3_ (1848) and L-Ara-4N modified lipid A (1928) are shown. (**B**) ESI-MS spectra of lipid A molecules extracted from *E. coli* UTI-3 cells following acute Pi-starvation treatment (Pi-, 0 mM) (right) compared to the group without treatment (Pi+, 1.32 mM) (left). Peaks correspond to native lipid A lost a HPO_3_ (1718), native lipid A (1798), L-Ara-4N modified lipid A lost a HPO_3_ (1848) and L-Ara-4N modified lipid A (1928) are shown. (**C**) ESI-MS spectra of lipid A molecules extracted from *S.* Typhimurium ATCC14028 cells following acute Pi-starvation treatment (Pi-, 0 mM) (right) compared to the group without treatment (Pi+, 1.32 mM) (left). Peaks correspond to native lipid A lost a HPO_3_ (1718), native lipid A (1798), L-Ara-4N modified lipid A lost a HPO_3_ (1848), L-Ara-4N and -OH modified lipid A lost a HPO_3_ (1863) and L-Ara-4N modified lipid A (1928) are shown. (**D**) ESI-MS spectra of lipid A molecules extracted from PAO1 cells following acute Pi-starvation treatment (Pi-, 0 mM) (right) compared to the group without treatment (Pi+, 1.32 mM) (left). Peaks correspond to hydroxylated lipid A lost a HPO_3_ (1366), di-hydroxylated lipid A lost a HPO_3_ (1382) are shown.

**Figure S5.**
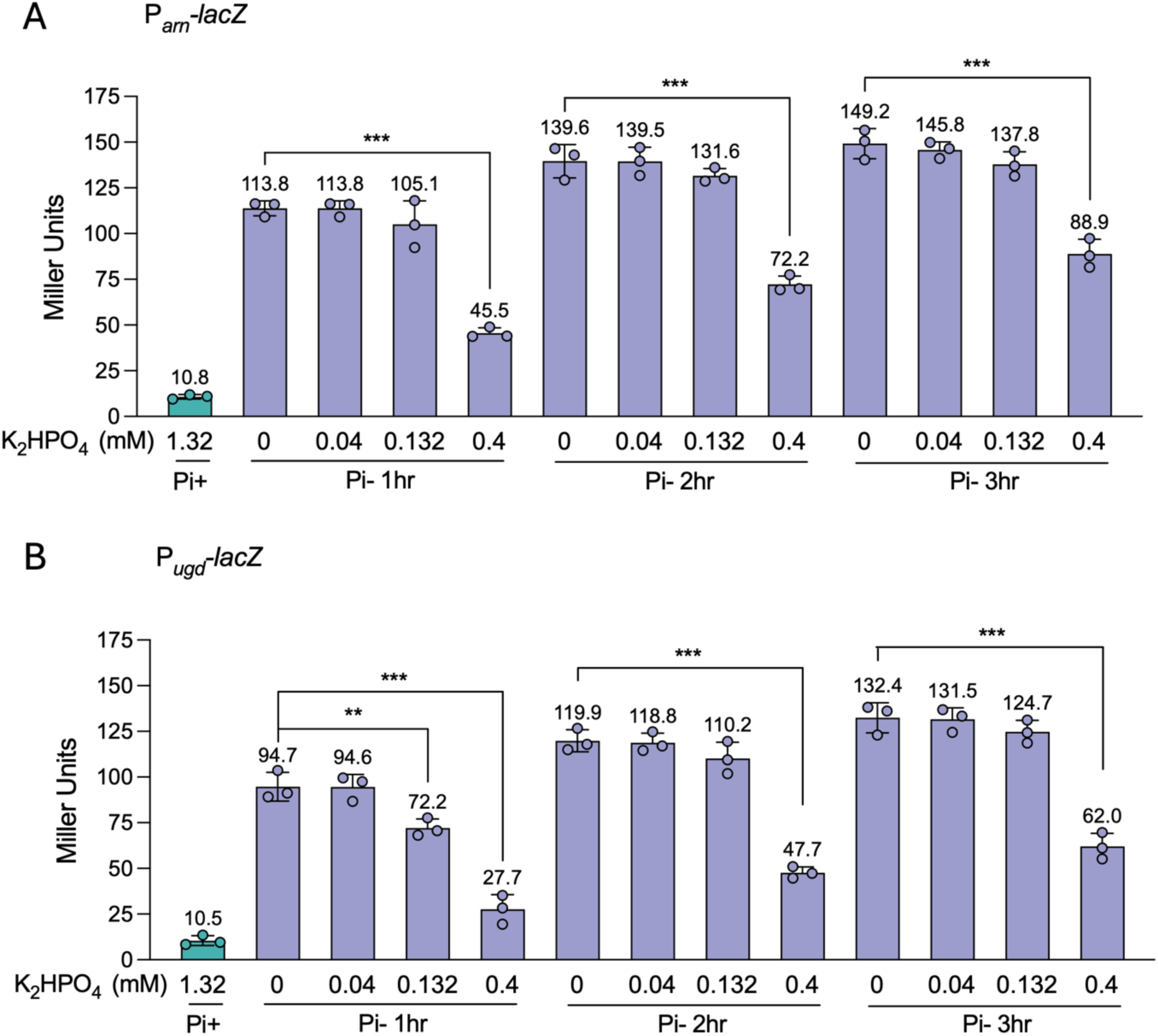
P*arn* and P*ugd* promoter activities in *E. coli* MG1655 during Pi depletion supplemented with different concentration of Pi. (A) β-galactosidase activities of P*_arn_*-*lacZ* in MG1655 Δ*lacZ* cells following Pi-starvation treatment with different concentration of Pi (Pi-, 0-0.4 mM) compared to the group without treatment (Pi+, 1.32 mM). (B) β-galactosidase activities of P*_ugd_*-*lacZ* in MG1655 Δ*lacZ* cells following Pi-starvation treatment with different concentration of Pi (Pi-, 0-0.4 mM) compared to the group without treatment (Pi+, 1.32 mM). Error bars represent the standard deviation of at least three independent experiments. Statistical significance was determined using one-way ANOVA followed by Dunnett’s multiple comparisons test. Significance levels are indicated as follows: \**p* < 0.05; \*\**p* < 0.01; \*\*\**p* < 0.001.

**Figure S6.**
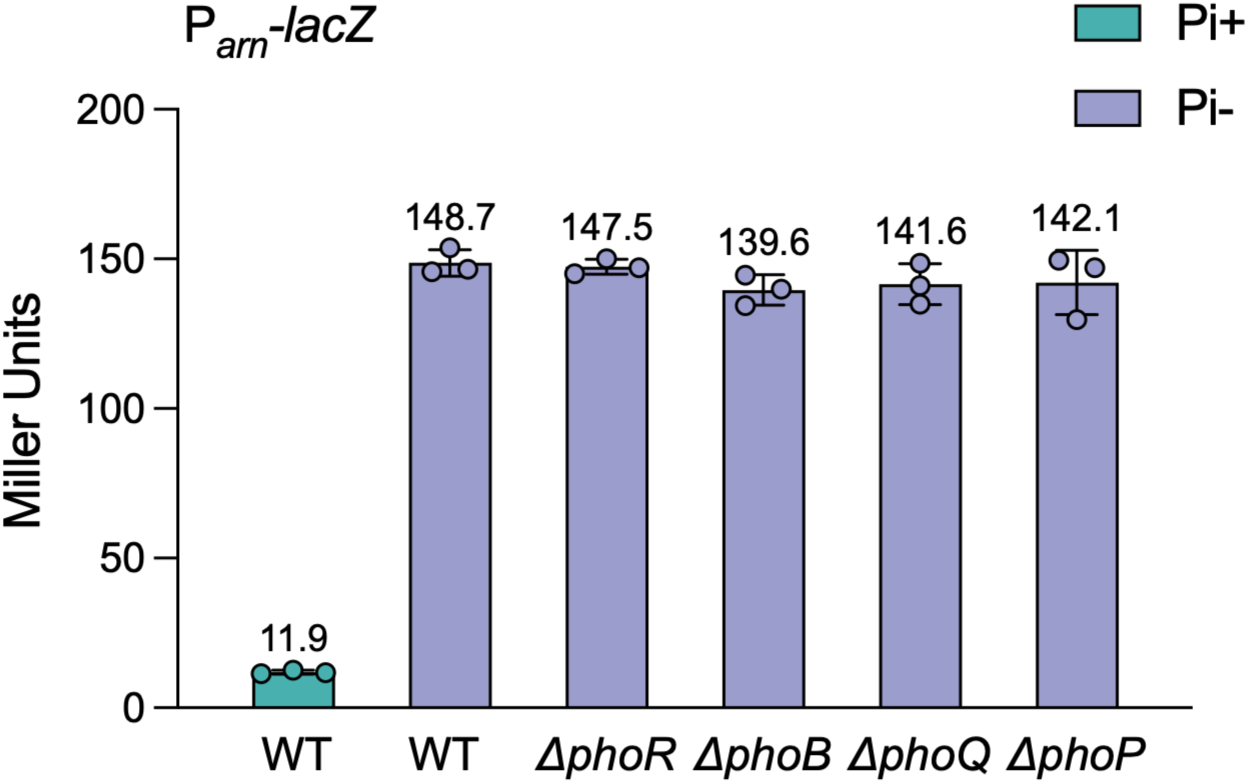
Up-regulation of *arn* genes during Pi depletion is independent of PhoBR and PhoPQ TCS in *E. coli* MG1655. β-galactosidase activities of P*_ugd_-lacZ* in MG1655 Δ*lacZ* parent and isogenic Δ*pmrB,* Δ*pmrA,* Δ*phoB* and Δ*phoR* mutants following acute Pi-starvation treatment (Pi-, 0 mM) compared to the group without treatment (Pi+, 1.32 mM). Error bars represent the standard deviation of at least three independent experiments. Statistical significance was determined using one-way ANOVA followed by Dunnett’s multiple comparisons test. Significance levels are indicated as follows: \**p* < 0.05; \*\**p* < 0.01; \*\*\**p* < 0.001.

**Figure S7.**
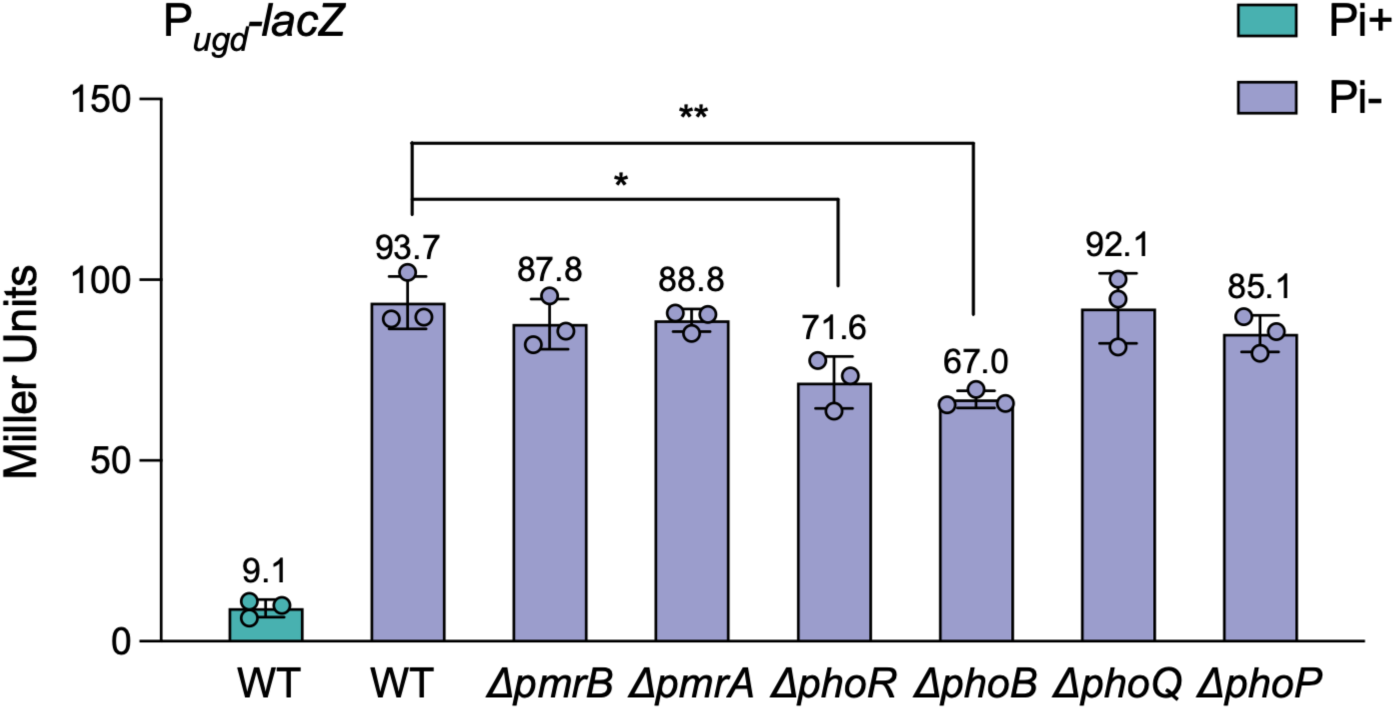
Up-regulation of *ugd* gene during Pi depletion is independent of PmrAB and PhoPQ TCS, but partially dependent on PhoBR TCS in *E. coli* MG1655. β-galactosidase activities of P*_arn_-lacZ* in MG1655 Δ*lacZ* parent and isogenic Δ*phoR,* Δ*phoB,* Δ*phoP* and Δ*phoQ* mutants following acute Pi-starvation treatment (Pi-, 0 mM) compared to the group without treatment (Pi+, 1.32 mM). Error bars represent the standard deviation of at least three independent experiments. Statistical significance was determined using one-way ANOVA followed by Dunnett’s multiple comparisons test. Significance levels are indicated as follows: \**p* < 0.05; \*\**p* < 0.01; \*\*\**p* < 0.001.

**Figure S8.**
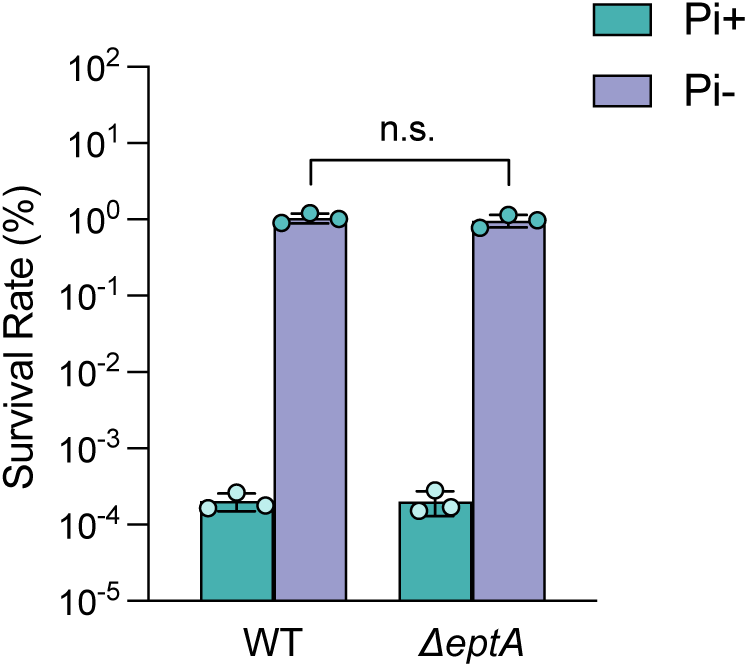
Pi depletion induced polymyxin tolerance is not affected in *eptA* deletion strain. Survival of *E. coli* MG1655 parent and isogenic ΔeptA mutant exposed 1-hour to polymyxin B (4 μg/ml) following acute Pi-starvation treatment (Pi-, 0 mM) compared to the group without treatment (Pi+, 1.32 mM). Error bars represent the standard deviation of at least three independent experiments. Statistical significance was determined using two-tailed Student’s t-test. Significance levels are indicated as follows: \**p* < 0.05; \*\**p* < 0.01; \*\*\**p* < 0.001.

**Figure S9.**
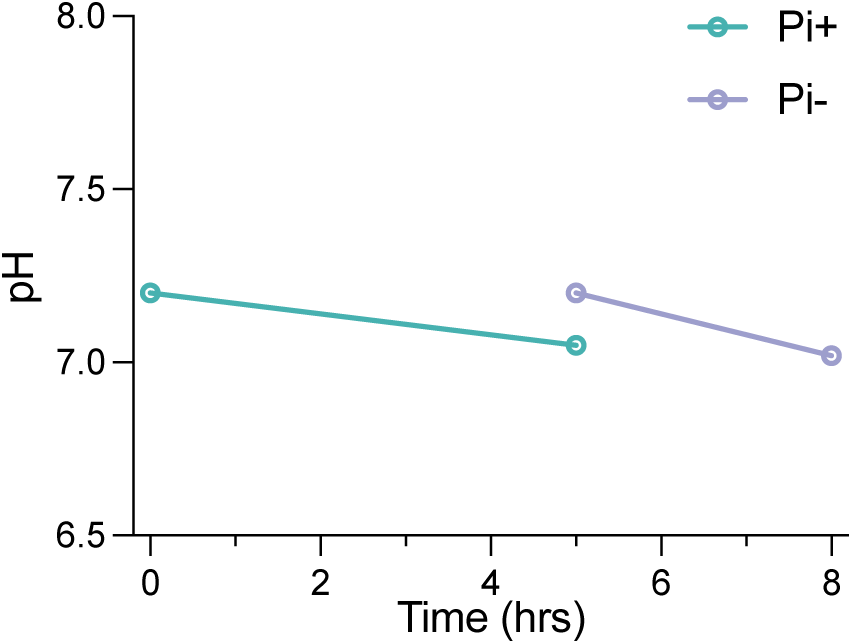
pH of spent medium of *E. coli* MG1655 after 5hrs Pi sufficient growth and after 3hrs Pi depletion treatment. pH of the medium before bacteria inoculating and pH of the spent medium after 5hrs Pi sufficient growth or 3hrs Pi depletion treatment were measured. Error bars represent the standard deviation of at least three independent experiments.

**Figure S10.**
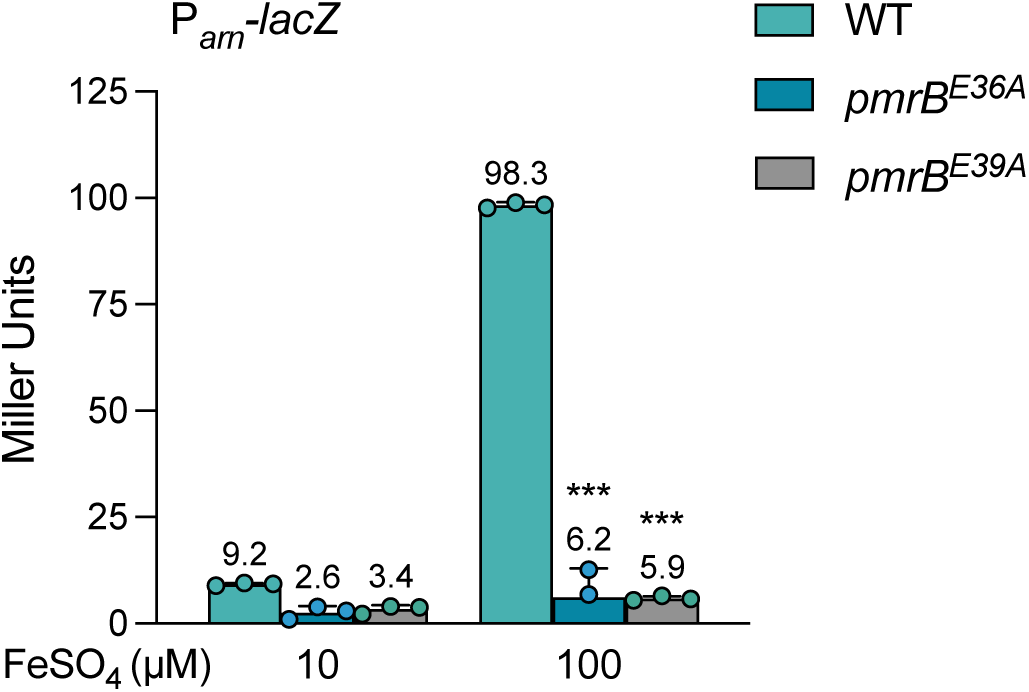
E36A and E39A mutation abolish the Fe^3+^ signaling capacity of PmrA. β-galactosidase activities of P*_arn_-lacZ* in MG1655 Δ*lacZ* parent and isogenic *pmrB*^E36A^*, pmrB*^E39A^mutants growth under MOPS minimal medium supplemented with different concentration of FeSO_4_. Error bars represent the standard deviation of at least three independent experiments. Statistical significance was determined using two-way ANOVA followed by Dunnett’s multiple comparisons test. Significance levels are indicated as follows: \**p* < 0.05; \*\**p* < 0.01; \*\*\**p* < 0.001.

**Figure S11.**
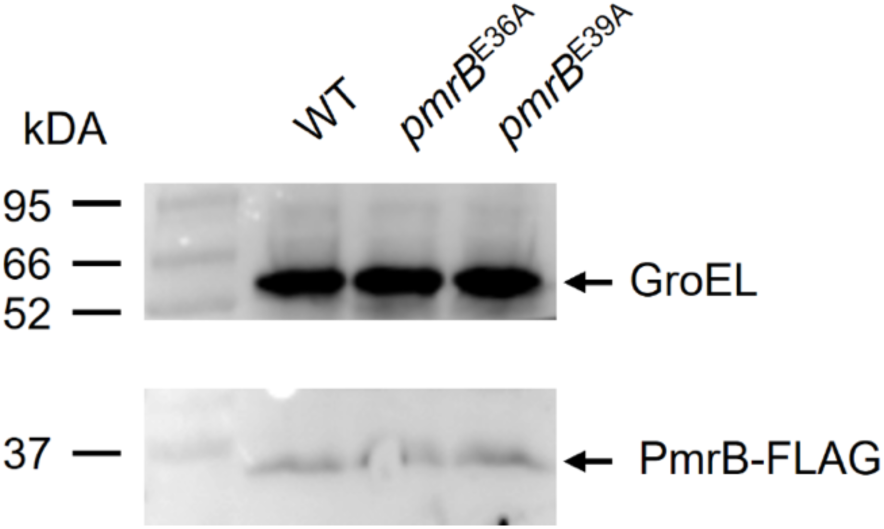
PmrB^E36A^ and PmrB^E39A^ exhibit expression levels comparable to wild-type PmrB. Western blot analysis of PmrB-FLAG expression in strains expressing wild-type PmrB, PmrB^E36A^ or PmrB^E39A^. GroEL was used as loading control.

**Figure S12.**
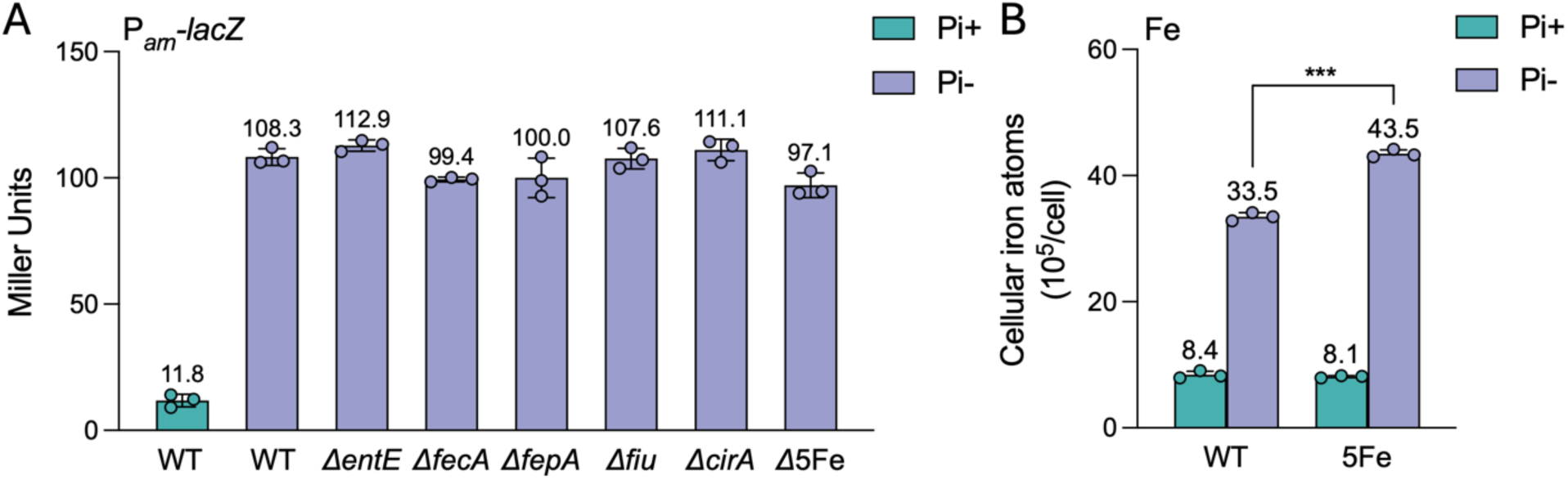
Deletion of canonical Fe^3+^ up-take genes does not affect the activation of PmrAB system and increase of Fe mobilization to *E. coli* cell during Pi depletion. (A) β-galactosidase activities of P*_arn_-lacZ* in MG1655 Δ*lacZ* parent and isogenic Δ*fecA*, Δ*fepA*, Δ*entE*, Δ*fiu*, Δ*cirA*, and Δ5Fe (Δ*fecA*Δ*fepA*Δ*entE*Δ*fiu*Δ*cirA*) mutants following acute Pi-starvation treatment (Pi-, 0 mM) compared to the group without treatment (Pi+, 1.32 mM). (B) Fe contents in *E. coli* MG1655 parent and isogenic Δ*fecA* Δ*fepA* Δ*entE* Δ*fiu* Δ*cirA* mutant following acute Pi-starvation treatment (Pi-, 0 mM) compared to the group without treatment (Pi+, 1.32 mM). Error bars represent the standard deviation of at least three independent experiments. Statistical significance was determined using one-way ANOVA followed by Dunnett’s multiple comparisons test for **A** and two-way ANOVA followed by Sidak’s multiple comparisons test for **B**. Significance levels are indicated as follows: \**p* < 0.05; \*\**p* < 0.01; \*\*\**p* < 0.001.

**Figure S13.**
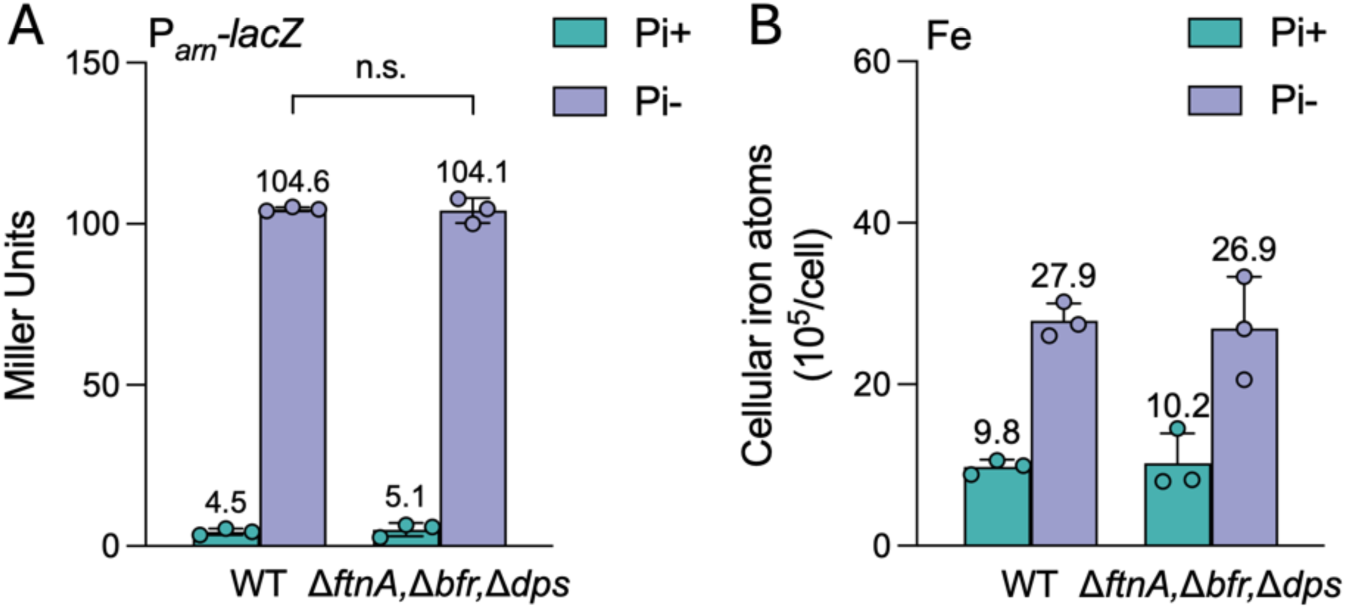
Deletion of Fe storage genes does not affect the activation of PmrAB system and increase of Fe mobilization to *E. coli* cell during Pi depletion. (A) β-galactosidase activities of P*_arn_-lacZ* in MG165ss5 Δ*lacZ* parent and isogenic Δ*ftnA* Δ*bfr* Δ*dps* mutant following acute Pi-starvation treatment (Pi-, 0 mM) compared to the group without treatment (Pi+, 1.32 mM). (B) Fe contents in *E. coli* MG1655 parent and isogenic Δ*ftnA* Δ*bfr* Δ*dps* mutant following acute Pi-starvation treatment (Pi-, 0 mM) compared to the group without treatment (Pi+, 1.32 mM). Error bars represent the standard deviation of at least three independent experiments. Statistical significance was determined using one-way ANOVA followed by Dunnett’s multiple comparisons test for **A** and two-way ANOVA followed by Sidak’s multiple comparisons test for **B**. Significance levels are indicated as follows: \**p* < 0.05; \*\**p* < 0.01; \*\*\**p* < 0.001.

**Figure S14.**
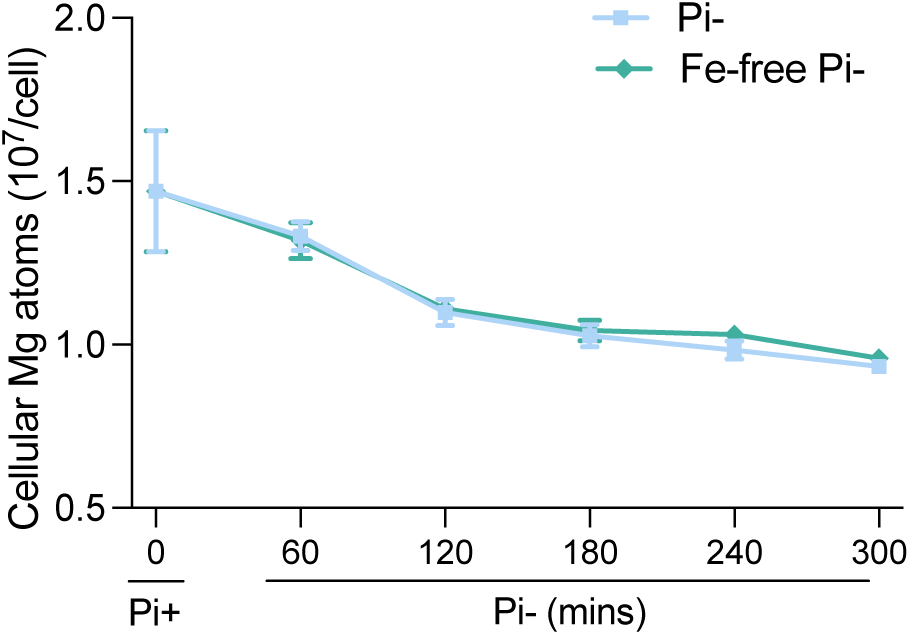
Decrease of cellular magnesium during Pi depletion is independent of Fe availability in medium. Mg contents in *E. coli* MG1655 cell following Pi depletion treatment (0 Pi, 10 μM FeSO_4_) or Fe-free Pi depletion treatment (0 Pi, 0 FeSO_4_) at the indicated time points. Error bars represent the standard deviation of at least three independent experiments.

**Figure S15.**
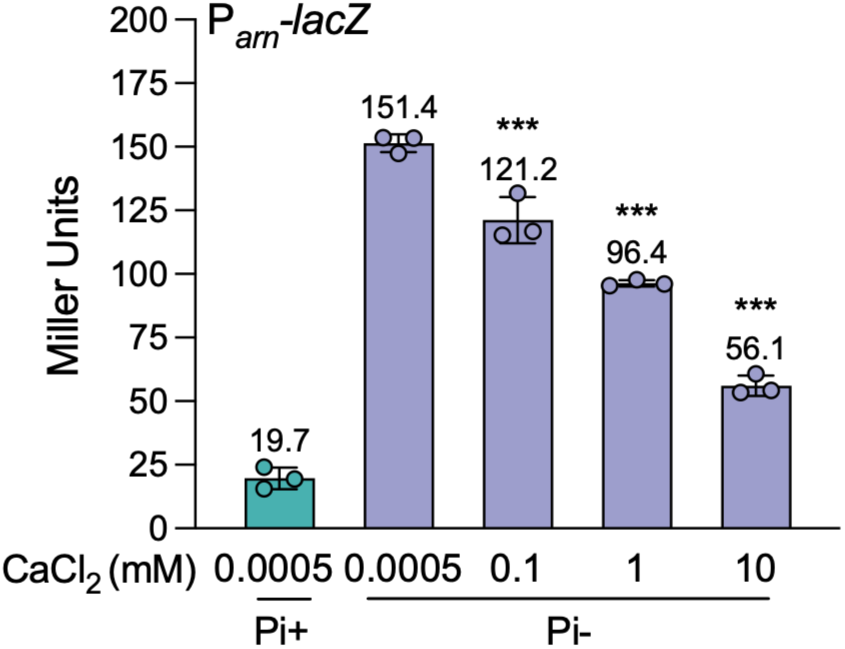
Excess Ca supplementation could reverse the activation of PmrAB system during Pi depletion. β-galactosidase activities of P*_arn_-lacZ* in MG1655 Δ*lacZ* cells following acute Pi-starvation treatment (Pi-, 0 mM Pi, 0.0005 mM Ca) or Ca excess Pi depletion treatment (Pi-, 0 mM Pi, 0.0005-10 mM Ca) compared to the group without treatment (Pi+, 1.32 mM, 0.0005 mM Ca). Error bars represent the standard deviation of at least three independent experiments. Statistical significance was determined using one-way ANOVA followed by Dunnett’s multiple comparisons test. Significance levels are indicated as follows: \**p* < 0.05; \*\**p* < 0.01; \*\*\**p* < 0.001.

**Figure S16.**
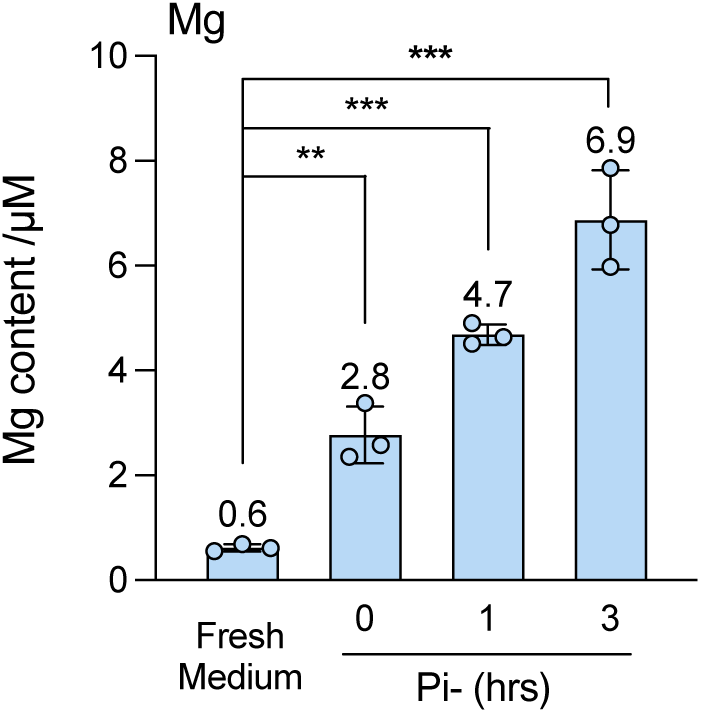
Magnesium disassociates from *E. coli* cells when subjected to Mg-free Pi-depleted medium. Mg content in the spent medium during Mg-free Pi depletion treatment (0 Pi, 0 MgCl_2_) at the indicated time points. Error bars represent the standard deviation of at least three independent experiments. Statistical significance was determined using one-way ANOVA followed by Dunnett’s multiple comparisons test. Significance levels are indicated as follows: \**p* < 0.05; \*\**p* < 0.01; \*\*\**p* < 0.001.

**Figure S17.**
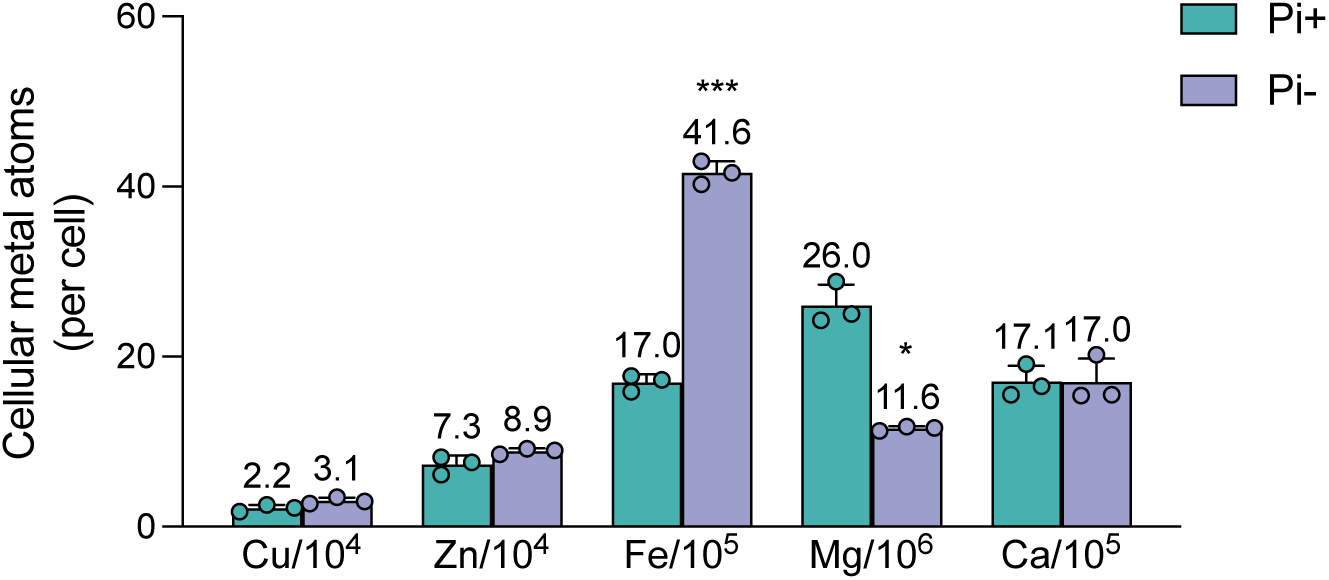
Cellular metal content in *P. aeruginosa* PAO1 cells with Pi depletion treatment and without Pi depletion treatment. Cellular metal contents determined by ICP-MS in *P. aeruginosa* PAO1 cells following acute Pi-starvation treatment (Pi-, 0 mM) compared to the group without treatment (Pi+, 1.32 mM). Error bars represent the standard deviation of at least three independent experiments. Statistical significance was determined using two-tailed Student’s t-test. Significance levels are indicated as follows: \**p* < 0.05; \*\**p* < 0.01; \*\*\**p* < 0.001.

**Figure S18.**
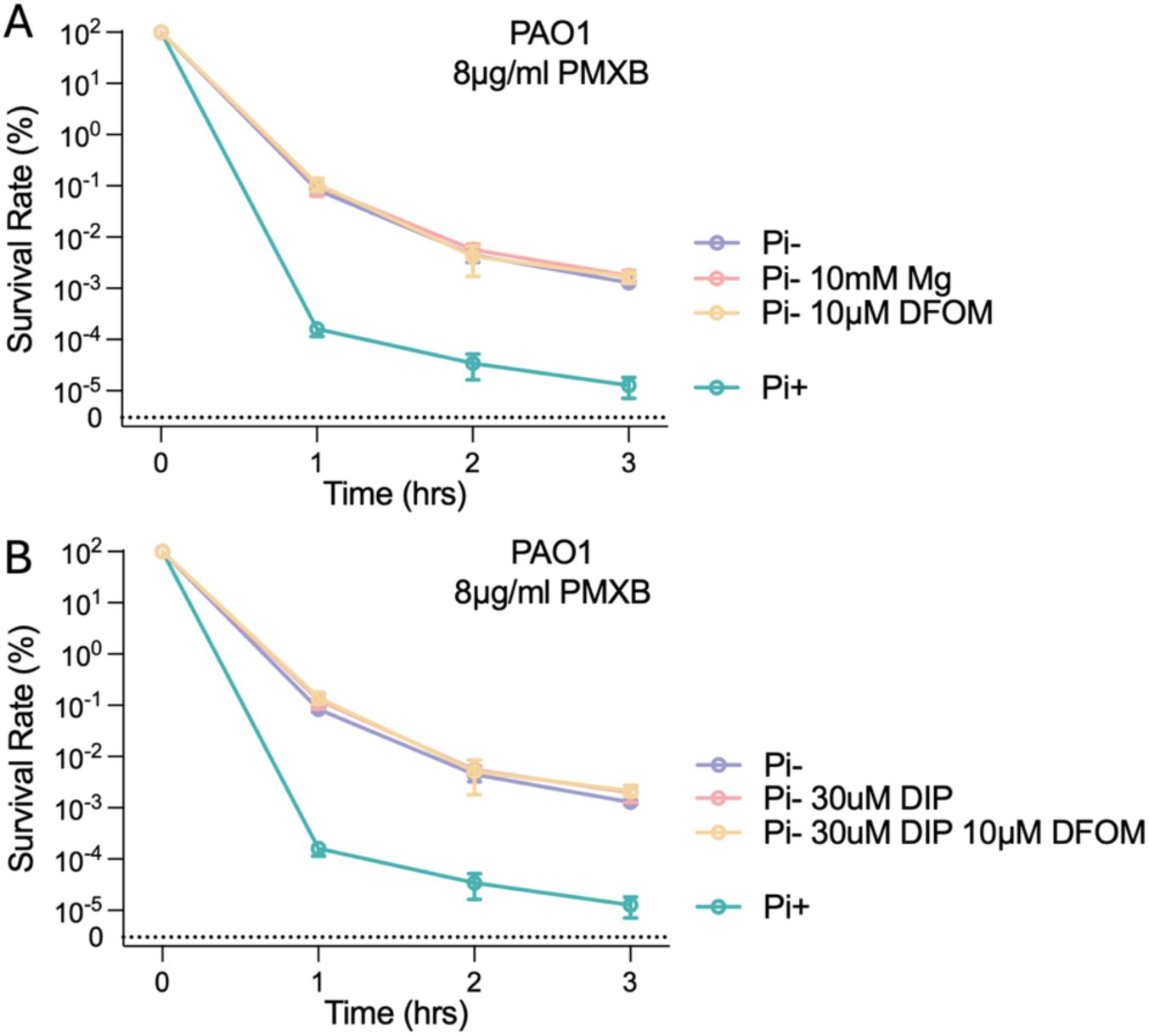
Disruption of Mg and Fe signaling showed no effect on Pi depletion induced polymyxin tolerance in *P. aeruginosa.* (A) Time-kill curves of *P. aeruginosa* PAO1 exposed to PMXB (8 μg/ml) following acute Pi-starvation treatment (Pi-, 0 mM Pi), Pi depletion treatment with excess Mg (Pi-, 0 mM Pi, 10mM MgCl_2_) or Pi depletion treatment with iron chelator (Pi-, 0 mM Pi, 10 μM DFOM) compared to the group without treatment (Pi+, 1.32 mM Pi). The dotted line labelled “0” denotes the limit of detection; values below this threshold were not detectable by colony counting. (B) Time-kill curves of *P. aeruginosa* PAO1 exposed to PMXB (8 μg/ml) following acute Pi-starvation treatment (Pi-, 0 mM Pi), Pi depletion treatment with Fe^2+^ chelator (Pi-, 0 mM Pi, 30 μM DIP) or Pi depletion treatment with both Fe^2+^ and Fe^3+^chelator (Pi-, 0 mM Pi, 30 μM DIP, 10 μM DFOM) compared to the group without treatment (Pi+, 1.32 mM Pi). The dotted line labelled “0” denotes the limit of detection; values below this threshold were not detectable by colony counting. Error bars represent the standard deviation of at least three independent experiments. Statistical significance was determined using one-way ANOVA followed by Dunnett’s multiple comparisons test. Significance levels are indicated as follows: \**p* < 0.05; \*\**p* < 0.01; \*\*\**p* < 0.001.

**Figure S19.**
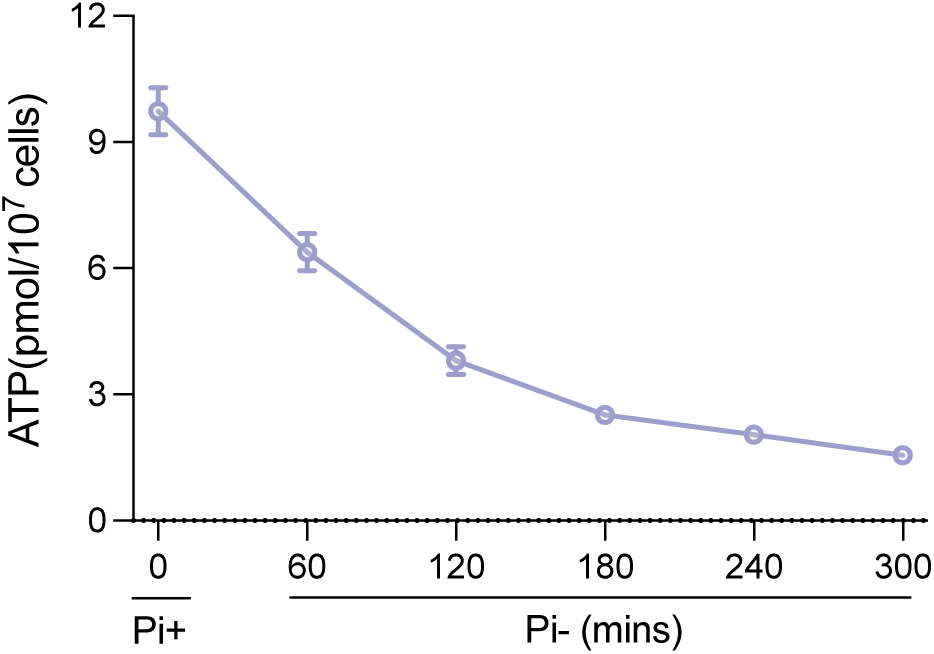
Cellular ATP level in *E. coli* MG1655 cells continuously decrease during Pi depletion. Cellular ATP levels in *E. coli* MG1655 cells following Pi depletion treatment at the specified time points. Error bars represent the standard deviation of at least three independent experiments.

**Figure S20.**
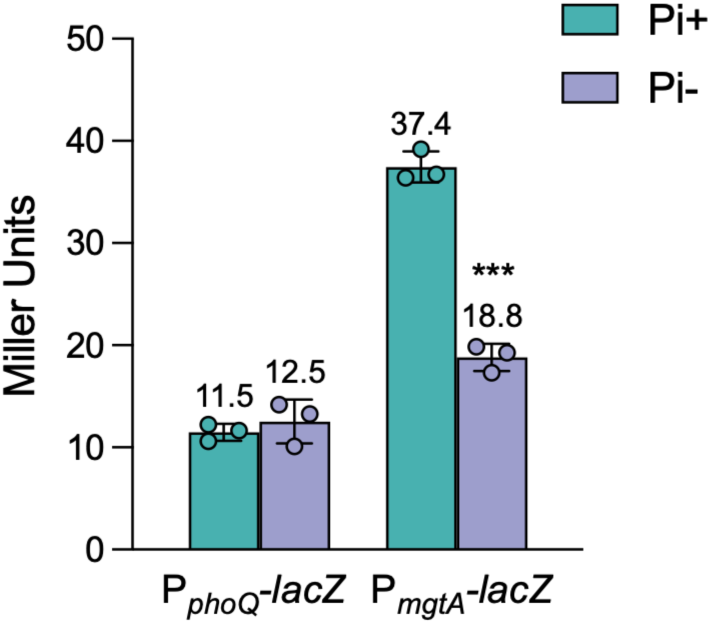
Lack of upregulation of P*phoQ* and P*mgtA* promoter activities indicates that the PhoPQ TCS is not activated during Pi depletion in *E. coli* MG1655. β-galactosidase activities of P*_phoQ_-lacZ* and P*_mgtA_-lacZ* in MG1655 Δ*lacZ* parent following acute Pi-starvation treatment (Pi-, 0 mM) compared to the group without treatment (Pi+, 1.32 mM). Error bars represent the standard deviation of at least three independent experiments. Statistical significance was determined using two-tailed Student’s t-test. Significance levels are indicated as follows: \**p* < 0.05; \*\**p* < 0.01; \*\*\**p* < 0.001.

**Figure S21.**
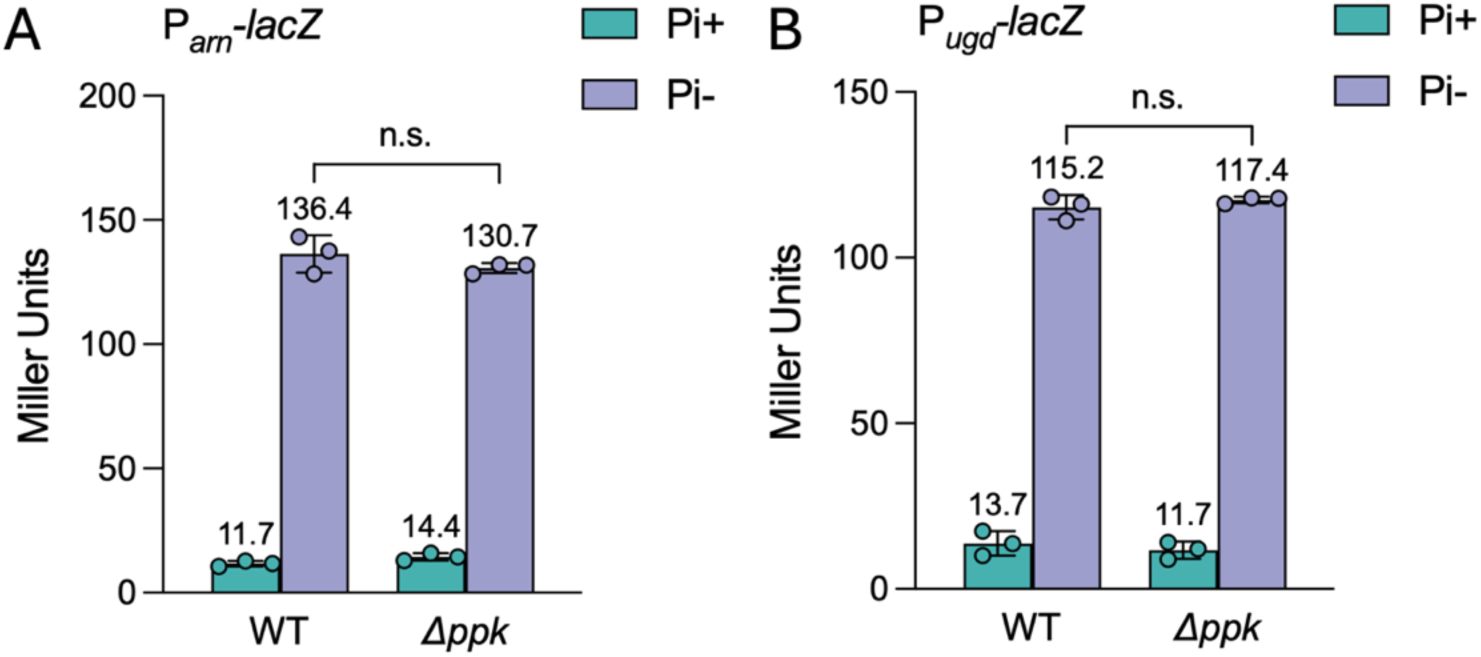
Up-regulation of both P*arn* and P*ugd* promoter activities during Pi depletion are not affected in *ppk* deletion strain. (A) β-galactosidase activities of P*_arn_-lacZ* in MG1655 Δ*lacZ* parent and isogenic Δ*ppk* mutant following acute Pi-starvation treatment (Pi-, 0 mM) compared to the group without treatment (Pi+, 1.32 mM). (B) β-galactosidase activities of P*_ugd_-lacZ* in MG1655 Δ*lacZ* parent and isogenic Δ*ppk* mutant following acute Pi-starvation treatment (Pi-, 0 mM) compared to the group without treatment (Pi+, 1.32 mM). Error bars represent the standard deviation of at least three independent experiments. Statistical significance was determined using two-way ANOVA followed by Tukey’s multiple comparisons test. Significance levels are indicated as follows: \**p* < 0.05; \*\**p* < 0.01; \*\*\**p* < 0.001.

**Figure S22.**
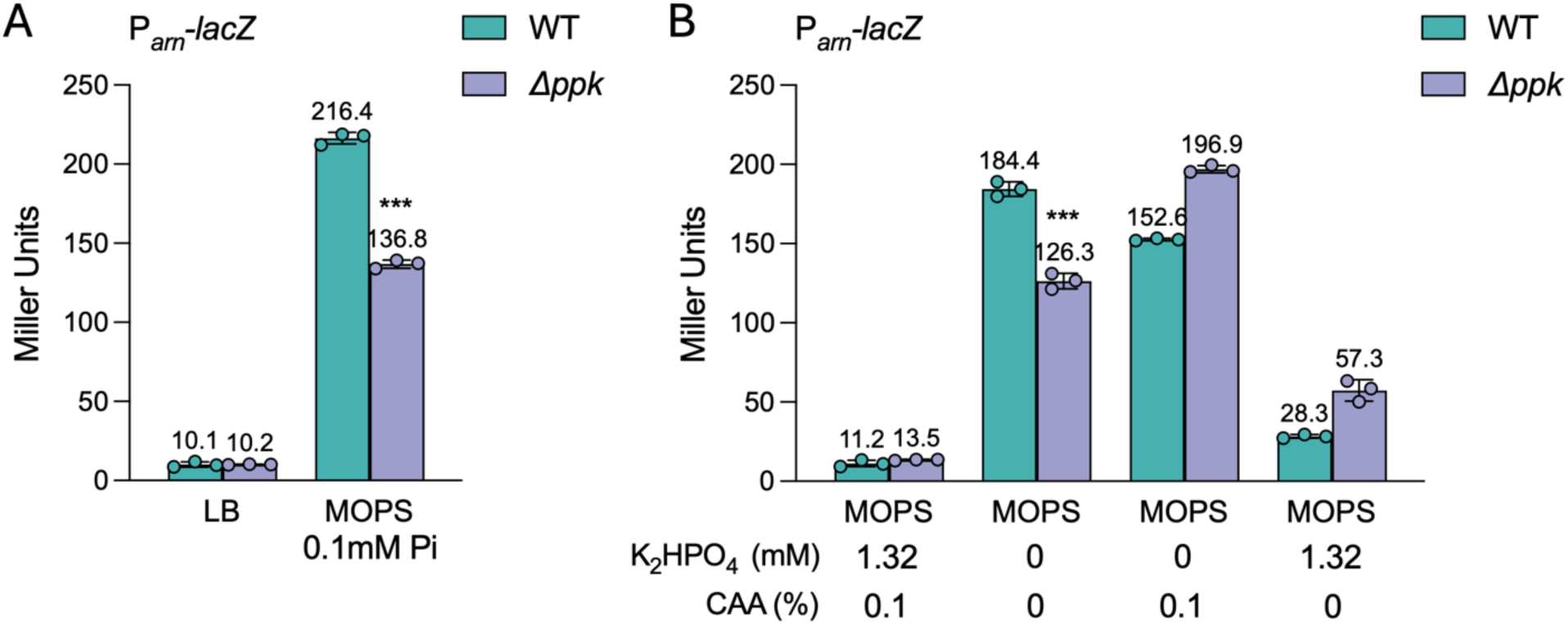
Pi depletion, amino acid depletion, and combined Pi–amino acid stress upregulate P*arn* promoter activity, while *ppk* deletion only attenuates upregulation under combined stress. (A) β-galactosidase activities of P*_arn_-lacZ* in MG1655 Δ*lacZ* parent and isogenic Δ*ppk* mutant following acute nutrients downshift (MOPS 0.1 mM Pi) compared to the group without treatment (LB). Error bars represent the standard deviation of at least three independent experiments. Statistical significance was determined using two-way ANOVA followed by Tukey’s multiple comparisons test. (B) β-galactosidase activities of P*_arn_-lacZ* in MG1655 Δ*lacZ* parent and isogenic Δ*ppk* mutant following Pi depletion treatment (MOPS 0 Pi, 0.1% CAA), amino acid depletion treatment (MOPS 1.32 mM Pi, 0 CAA) and combined Pi–amino acid stress treatment (MOPS 0 Pi, 0 CAA) compared to the group without treatment (MOPS 1.32 mM Pi, 0.1% CAA). Error bars represent the standard deviation of at least three independent experiments. Statistical significance was determined using two-way ANOVA followed by Tukey’s multiple comparisons test. Significance levels are indicated as follows: \**p* < 0.05; \*\**p* < 0.01; \*\*\**p* < 0.001.

---

**Table S1.** Minimum inhibitory concentrations (MICs) of strains in MOPS minimal medium.

| **Strain** | **PMXB (ug/ml)** |
|---|---|
| MG1655 | 0.5 |
| O157 | 1 |
| UTI-1 | 0.5 |
| UTI-3 | 1 |
| UTI-4 | 1 |
| UTI-5 | 1 |
| PAO1 | 0.5 |
| ATCC 14028 | 2 |

**Table S2.** Proteins identified to be different expressed in *E. coli* MG1655 growing under Pi sufficient and depletion conditions.

| **prot_desc** | **id** | **log2FoldChange** | **padj** |
|---|---|---|---|
| argF | sp\|P06960\|OTC2_ECOLI | -Inf | 3.42E-18 |
| gatZ | sp\|P0C8J8\|GATZ_ECOLI | -Inf | 3.77E-09 |
| fis | sp\|P0A6R3\|FIS_ECOLI | -Inf | 2.78E-08 |
| secY | sp\|P0AGA2\|SECY_ECOLI | -Inf | 0.000159211 |
| kgtP | sp\|P0AEX3\|KGTP_ECOLI | -Inf | 0.001647533 |
| gatC | sp\|P69831\|PTKC_ECOLI | -Inf | 0.002435201 |
| cysC | sp\|P0A6J1\|CYSC_ECOLI | -Inf | 0.003341414 |
| znuC | sp\|P0A9X1\|ZNUC_ECOLI | -Inf | 0.025826417 |
| rffH | sp\|P61887\|RMLA2_ECOLI | -Inf | 0.036760697 |
| cysD | sp\|P21156\|CYSD_ECOLI | -5.525585778 | 1.38E-11 |
| mnmG | sp\|P0A6U3\|MNMG_ECOLI | -4.77187114 | 0.003253389 |
| argA | sp\|P0A6C5\|ARGA_ECOLI | -4.649638457 | 2.18E-06 |
| yhjE | sp\|P37643\|YHJE_ECOLI | -4.500028599 | 0.005992816 |
| yegQ | sp\|P76403\|YEGQ_ECOLI | -4.482584177 | 0.015352456 |
| cysN | sp\|P23845\|CYSN_ECOLI | -3.927062955 | 7.22E-14 |
| pncB | sp\|P18133\|PNCB_ECOLI | -3.639270989 | 6.73E-05 |
| fbp | sp\|P0A993\|F16PA_ECOLI | -3.600157407 | 0.023396164 |
| moaA | sp\|P30745\|MOAA_ECOLI | -3.529794204 | 0.018342621 |
| rpmC | sp\|P0A7M6\|RL29_ECOLI | -3.142935813 | 1.68E-07 |
| ilvN | sp\|P0ADF8\|ILVN_ECOLI | -3.08821322 | 0.03765939 |
| aroP | sp\|P15993\|AROP_ECOLI | -2.833642328 | 0.021648588 |
| deaD | sp\|P0A9P6\|DEAD_ECOLI | -2.695068638 | 0.006647318 |
| ilvB | sp\|P08142\|ILVB_ECOLI | -2.690257592 | 1.41E-07 |
| asnA | sp\|P00963\|ASNA_ECOLI | -2.66746299 | 0.015909254 |
| ptsG | sp\|P69786\|PTGCB_ECOLI | -2.510708646 | 0.000188954 |
| rpmD | sp\|P0AG51\|RL30_ECOLI | -2.495454148 | 8.47E-09 |
| ykgM | sp\|P0A7N1\|RL31B_ECOLI | -2.483011177 | 0.00644327 |
| gatB | sp\|P37188\|PTKB_ECOLI | -2.418194429 | 0.035274735 |
| cs | sp\|P0A972\|CSPE_ECOLI | -2.352007644 | 4.54E-08 |
| metA | sp\|P07623\|META_ECOLI | -2.317452293 | 0.000139049 |
| der | sp\|P0A6P5\|DER_ECOLI | -2.267771633 | 0.0102137 |
| potA | sp\|P69874\|POTA_ECOLI | -2.183307337 | 0.03275735 |
| leuA | sp\|P09151\|LEU1_ECOLI | -1.998105409 | 1.07E-08 |
| glf | sp\|P37747\|GLF_ECOLI | -1.97691323 | 0.029751 |
| tr | sp\|P00895\|TRPE_ECOLI | -1.943830992 | 0.005678046 |
| miaB | sp\|P0AEI1\|MIAB_ECOLI | -1.92205685 | 0.004244044 |
| pntB | sp\|P0AB67\|PNTB_ECOLI | -1.852085802 | 9.85E-06 |
| purF | sp\|P0AG16\|PUR1_ECOLI | -1.826541229 | 4.76E-05 |
| lepA | sp\|P60785\|LEPA_ECOLI | -1.791966706 | 0.01657399 |
| gatY | sp\|P0C8J6\|GATY_ECOLI | -1.776699008 | 0.000182727 |
| purT | sp\|P33221\|PURT_ECOLI | -1.764435997 | 0.000643699 |
| prfC | sp\|P0A7I4\|RF3_ECOLI | -1.76312872 | 0.012506321 |
| ybiC | sp\|P30178\|YBIC_ECOLI | -1.727397747 | 0.01941163 |
| rpsH | sp\|P0A7W7\|RS8_ECOLI | -1.723379737 | 2.80E-10 |
| purM | sp\|P08178\|PUR5_ECOLI | -1.678278829 | 0.003764 |
| cyoA | sp\|P0ABJ1\|CYOA_ECOLI | -1.663437618 | 0.006838543 |
| metL | sp\|P00562\|AK2H_ECOLI | -1.661017127 | 0.00181135 |
| leuC | sp\|P0A6A6\|LEUC_ECOLI | -1.653771358 | 6.45E-06 |
| pntA | sp\|P07001\|PNTA_ECOLI | -1.595004247 | 7.61E-08 |
| trxB | sp\|P0A9P4\|TRXB_ECOLI | -1.571960722 | 0.002958769 |
| metH | sp\|P13009\|METH_ECOLI | -1.565084962 | 0.000869601 |
| metC | sp\|P06721\|METC_ECOLI | -1.510217214 | 0.025144896 |
| rpmF | sp\|P0A7N4\|RL32_ECOLI | -1.498744569 | 0.000469522 |
| gyrA | sp\|P0AES4\|GYRA_ECOLI | -1.448436432 | 7.72E-05 |
| rpsR | sp\|P0A7T7\|RS18_ECOLI | -1.418043759 | 3.03E-05 |
| rpmG | sp\|P0A7N9\|RL33_ECOLI | -1.381024188 | 0.006747605 |
| purH | sp\|P15639\|PUR9_ECOLI | -1.376402854 | 0.000114658 |
| argI | sp\|P04391\|OTC1_ECOLI | -1.37321011 | 0.000772873 |
| rnb | sp\|P30850\|RNB_ECOLI | -1.364231063 | 0.003951903 |
| rplT | sp\|P0A7L3\|RL20_ECOLI | -1.361561823 | 7.98781E-05 |
| cysS | sp\|P21888\|SYC_ECOLI | -1.352510933 | 0.018153582 |
| metK | sp\|P0A817\|METK_ECOLI | -1.28929395 | 0.000291382 |
| sdhA | sp\|P0AC41\|SDHA_ECOLI | -1.284366661 | 8.12E-05 |
| ydjN | sp\|P77529\|YDJN_ECOLI | -1.282130778 | 0.012801159 |
| pyrD | sp\|P0A7E1\|PYRD_ECOLI | -1.24642368 | 0.017272295 |
| rpsM | sp\|P0A7S9\|RS13_ECOLI | -1.231500265 | 4.82E-05 |
| rpsC | sp\|P0A7V3\|RS3_ECOLI | -1.228519333 | 5.14E-06 |
| rplU | sp\|P0AG48\|RL21_ECOLI | -1.200353979 | 0.000935979 |
| pliG | sp\|P76002\|PLIG_ECOLI | -1.192616084 | 0.023769411 |
| rplB | sp\|P60422\|RL2_ECOLI | -1.160209361 | 1.29E-05 |
| rpsA | sp\|P0AG67\|RS1_ECOLI | -1.154439475 | 2.19E-06 |
| rplS | sp\|P0A7K6\|RL19_ECOLI | -1.140490911 | 0.042048579 |
| rpsT | sp\|P0A7U7\|RS20_ECOLI | -1.136468494 | 0.000373058 |
| thrA | sp\|P00561\|AK1H_ECOLI | -1.131400346 | 7.55E-06 |
| thrS | sp\|P0A8M3\|SYT_ECOLI | -1.130800143 | 0.017957548 |
| rplR | sp\|P0C018\|RL18_ECOLI | -1.127973054 | 0.000234204 |
| rplX | sp\|P60624\|RL24_ECOLI | -1.108686244 | 0.000297915 |
| aceB | sp\|P08997\|MASY_ECOLI | -1.103226647 | 7.72E-05 |
| aroA | sp\|P0A6D3\|AROA_ECOLI | -1.074448834 | 0.048313271 |
| rplI | sp\|P0A7R1\|RL9_ECOLI | -1.074352178 | 3.31E-05 |
| rplK | sp\|P0A7J7\|RL11_ECOLI | -1.030031069 | 0.000940436 |
| argH | sp\|P11447\|ARLY_ECOLI | -1.029621386 | 0.010350803 |
| cspC | sp\|P0A9Y6\|CSPC_ECOLI | -1.010776163 | 0.00524746 |
| gpmA | sp\|P62707\|GPMA_ECOLI | 1.007836346 | 0.000278112 |
| gmhA | sp\|P63224\|GMHA_ECOLI | 1.02030343 | 0.027607596 |
| luxS | sp\|P45578\|LUXS_ECOLI | 1.049588806 | 0.000506408 |
| rpiA | sp\|P0A7Z0\|RPIA_ECOLI | 1.052561101 | 0.01318169 |
| pgi | sp\|P0A6T1\|G6PI_ECOLI | 1.065800857 | 0.000188954 |
| pykF | sp\|P0AD61\|KPYK1_ECOLI | 1.081019653 | 2.73E-05 |
| mlaC | sp\|P0ADV7\|MLAC_ECOLI | 1.117643099 | 0.006345313 |
| yfiF | sp\|P0AGJ5\|YFIF_ECOLI | 1.128896995 | 0.03275735 |
| uspG | sp\|P39177\|USPG_ECOLI | 1.143746641 | 0.03765939 |
| pgl | sp\|P52697\|6PGL_ECOLI | 1.167828441 | 0.01318169 |
| deoC | sp\|P0A6L0\|DEOC_ECOLI | 1.174516518 | 0.018153582 |
| grxB | sp\|P0AC59\|GLRX2_ECOLI | 1.178432467 | 0.000732045 |
| fbaB | sp\|P0A991\|ALF1_ECOLI | 1.197658464 | 7.99E-05 |
| adhE | sp\|P0A9Q7\|ADHE_ECOLI | 1.201151692 | 3.88E-07 |
| degP | sp\|P0C0V0\|DEGP_ECOLI | 1.206061871 | 0.000278112 |
| tas | sp\|P0A9T4\|TAS_ECOLI | 1.219282681 | 0.036014767 |
| katG | sp\|P13029\|KATG_ECOLI | 1.220074858 | 7.01E-05 |
| ribC | sp\|P0AFU8\|RISA_ECOLI | 1.272415628 | 0.030285161 |
| pmbA | sp\|P0AFK0\|PMBA_ECOLI | 1.273270833 | 0.007784421 |
| ftnA | sp\|P0A998\|FTNA_ECOLI | 1.301974436 | 0.010044367 |
| moaB | sp\|P0AEZ9\|MOAB_ECOLI | 1.314241161 | 0.001746554 |
| wrbA | sp\|P0A8G6\|NQOR_ECOLI | 1.34050881 | 4.33E-06 |
| hdhA | sp\|P0AET8\|HDHA_ECOLI | 1.365397414 | 0.014224302 |
| yahK | sp\|P75691\|YAHK_ECOLI | 1.380128931 | 0.000137547 |
| acnA | sp\|P25516\|ACNA_ECOLI | 1.381383706 | 7.69E-05 |
| sbp | sp\|P0AG78\|SUBI_ECOLI | 1.410886097 | 0.001647533 |
| degQ | sp\|P39099\|DEGQ_ECOLI | 1.437968753 | 0.005431624 |
| tktB | sp\|P33570\|TKT2_ECOLI | 1.45986506 | 7.28E-07 |
| yjgA | sp\|P0A8X0\|YJGA_ECOLI | 1.470361111 | 0.0375636 |
| glgP | sp\|P0AC86\|PHSG_ECOLI | 1.489480956 | 0.004703968 |
| yggE | sp\|P0ADS6\|YGGE_ECOLI | 1.497996423 | 9.00E-05 |
| mtlD | sp\|P09424\|MTLD_ECOLI | 1.508297143 | 0.007422869 |
| cbpA | sp\|P36659\|CBPA_ECOLI | 1.520821408 | 0.040987772 |
| deoB | sp\|P0A6K6\|DEOB_ECOLI | 1.533048098 | 8.38E-05 |
| ybjP | sp\|P75818\|YBJP_ECOLI | 1.536131134 | 0.007427731 |
| ybaY | sp\|P77717\|YBAY_ECOLI | 1.541925932 | 0.00084693 |
| tdcE | sp\|P42632\|TDCE_ECOLI | 1.549713435 | 0.000108916 |
| thiF | sp\|P30138\|THIF_ECOLI | 1.58507647 | 0.00235383 |
| hdeB | sp\|P0AET2\|HDEB_ECOLI | 1.585194293 | 0.001902541 |
| tpiA | sp\|P0A858\|TPIS_ECOLI | 1.585411082 | 9.28E-09 |
| uspE | sp\|P0AAC0\|USPE_ECOLI | 1.605682754 | 0.000278112 |
| rraA | sp\|P0A8R0\|RRAA_ECOLI | 1.607904489 | 0.000417551 |
| yeaY | sp\|P0AA91\|YEAY_ECOLI | 1.609759679 | 0.001461256 |
| nagD | sp\|P0AF24\|NAGD_ECOLI | 1.651426571 | 0.020565443 |
| slp | sp\|P37194\|SLP_ECOLI | 1.661057589 | 0.000127411 |
| osmC | sp\|P0C0L2\|OSMC_ECOLI | 1.671087683 | 3.90E-06 |
| ycfP | sp\|P0A8E1\|YCFP_ECOLI | 1.676895218 | 0.000511934 |
| acuI | sp\|P26646\|ACUI_ECOLI | 1.680553338 | 0.001851227 |
| pflB | sp\|P09373\|PFLB_ECOLI | 1.700478434 | 9.87E-14 |
| lpoB | sp\|P0AB38\|LPOB_ECOLI | 1.721279546 | 0.000376072 |
| clpB | sp\|P63284\|CLPB_ECOLI | 1.727347175 | 1.23E-12 |
| gstB | sp\|P0ACA7\|GSTB_ECOLI | 1.748580068 | 0.001647533 |
| slt | sp\|P0AGC3\|SLT_ECOLI | 1.751322485 | 0.015251337 |
| yggN | sp\|P0ADS9\|YGGN_ECOLI | 1.768724665 | 0.049702565 |
| loiP | sp\|P25894\|LOIP_ECOLI | 1.795763405 | 0.012616436 |
| pspA | sp\|P0AFM6\|PSPA_ECOLI | 1.803047994 | 0.000834302 |
| chrR | sp\|P0AGE6\|CHRR_ECOLI | 1.829520642 | 0.00084693 |
| gstA | sp\|P0A9D2\|GSTA_ECOLI | 1.843282701 | 2.82E-06 |
| msrA | sp\|P0A744\|MSRA_ECOLI | 1.871059945 | 0.023254637 |
| dps | sp\|P0ABT2\|DPS_ECOLI | 1.8711205 | 1.92E-14 |
| yghU | sp\|Q46845\|YGHU_ECOLI | 1.947282377 | 0.000291495 |
| lldD | sp\|P33232\|LLDD_ECOLI | 1.949858867 | 0.035748827 |
| yfcH | sp\|P77775\|YFCH_ECOLI | 1.988861376 | 0.015909254 |
| cdd | sp\|P0ABF6\|CDD_ECOLI | 1.993269202 | 0.042932089 |
| yeaG | sp\|P0ACY3\|YEAG_ECOLI | 2.011516734 | 4.21E-11 |
| pqiB | sp\|P43671\|PQIB_ECOLI | 2.013507475 | 0.028681191 |
| ygaU | sp\|P0ADE6\|YGAU_ECOLI | 2.059465726 | 3.23E-14 |
| blc | sp\|P0A901\|BLC_ECOLI | 2.064974471 | 0.011100578 |
| qorA | sp\|P28304\|QOR1_ECOLI | 2.155671369 | 4.33E-07 |
| curA | sp\|P76113\|CURA_ECOLI | 2.175460559 | 0.010474873 |
| bfr | sp\|P0ABD3\|BFR_ECOLI | 2.180562547 | 1.56E-13 |
| yegP | sp\|P76402\|YEGP_ECOLI | 2.284730203 | 7.89E-10 |
| **ugd** | sp\|P76373\|UDG_ECOLI | 2.340441236 | 0.018153582 |
| yceF | sp\|P0A729\|YCEF_ECOLI | 2.382621181 | 0.040987772 |
| yecA | sp\|P0AD05\|YECA_ECOLI | 2.405409463 | 0.03275735 |
| bepA | sp\|P66948\|BEPA_ECOLI | 2.421711425 | 0.000774393 |
| otsA | sp\|P31677\|OTSA_ECOLI | 2.448868257 | 6.65E-08 |
| katE | sp\|P21179\|CATE_ECOLI | 2.50821371 | 2.35E-14 |
| hchA | sp\|P31658\|HCHA_ECOLI | 2.50879706 | 2.75E-12 |
| dkgA | sp\|Q46857\|DKGA_ECOLI | 2.649063918 | 1.64E-08 |
| ybeL | sp\|P0AAT9\|YBEL_ECOLI | 2.658797186 | 3.16E-08 |
| ymdB | sp\|P0A8D6\|YMDB_ECOLI | 2.671235984 | 0.024568779 |
| fruB | sp\|P69811\|PTFAH_ECOLI | 2.672541661 | 1.42E-05 |
| ydbC | sp\|P25906\|YDBC_ECOLI | 2.711914425 | 0.046758232 |
| ynhG | sp\|P76193\|YNHG_ECOLI | 2.733690087 | 3.31E-05 |
| yciE | sp\|P21363\|YCIE_ECOLI | 2.735728026 | 4.14E-05 |
| ymbA | sp\|P0AB10\|YMBA_ECOLI | 2.798163878 | 0.036198317 |
| ydcL | sp\|P64451\|YDCL_ECOLI | 2.82774542 | 2.04E-15 |
| ldhA | sp\|P52643\|LDHD_ECOLI | 2.837647013 | 2.28E-05 |
| amn | sp\|P0AE12\|AMN_ECOLI | 2.855927832 | 1.06E-13 |
| yehZ | sp\|P33362\|YEHZ_ECOLI | 2.946432256 | 0.017149234 |
| mdtE | sp\|P37636\|MDTE_ECOLI | 3.005567607 | 2.30E-05 |
| gadB | sp\|P69910\|DCEB_ECOLI | 3.026824738 | 1.75E-33 |
| yhjG | sp\|P37645\|YHJG_ECOLI | 3.114565954 | 0.004244044 |
| gadC | sp\|P63235\|GADC_ECOLI | 3.122545884 | 0.000234696 |
| gadA | sp\|P69908\|DCEA_ECOLI | 3.154673417 | 6.58E-36 |
| ydhF | sp\|P76187\|YDHF_ECOLI | 3.156625723 | 5.19E-05 |
| yjgR | sp\|P39342\|YJGR_ECOLI | 3.159652222 | 1.41E-12 |
| yghA | sp\|P0AG84\|YGHA_ECOLI | 3.175685027 | 1.40E-09 |
| ycaC | sp\|P21367\|YCAC_ECOLI | 3.185383669 | 2.44E-13 |
| phoE | sp\|P02932\|PHOE_ECOLI | 3.204721908 | 2.80E-10 |
| ycgM | sp\|P76004\|YCGM_ECOLI | 3.314835257 | 0.007920581 |
| tam | sp\|P76145\|TAM_ECOLI | 3.330887068 | 0.010474873 |
| yhbO | sp\|P45470\|YHBO_ECOLI | 3.335162899 | 0.000187361 |
| phoR | sp\|P08400\|PHOR_ECOLI | 3.473374236 | 0.001325039 |
| mdaB | sp\|P0AEY5\|MDAB_ECOLI | 3.504985139 | 1.26E-20 |
| dkgB | sp\|P30863\|DKGB_ECOLI | 3.638374711 | 5.12E-09 |
| yniA | sp\|P77739\|YNIA_ECOLI | 3.681925301 | 2.85E-07 |
| gldA | sp\|P0A9S5\|GLDA_ECOLI | 3.74947351 | 3.13E-05 |
| phnD | sp\|P16682\|PHND_ECOLI | 3.815420505 | 9.61E-10 |
| ugpQ | sp\|P10908\|UGPQ_ECOLI | 4.003964005 | 1.04E-06 |
| ahr | sp\|P27250\|AHR_ECOLI | 4.055942555 | 2.68E-05 |
| pstS | sp\|P0AG82\|PSTS_ECOLI | 4.113208109 | 3.15E-56 |
| cysQ | sp\|P22255\|CYSQ_ECOLI | 4.144617537 | 1.22E-07 |
| glsA1 | sp\|P77454\|GLSA1_ECOLI | 4.234450197 | 0.024320745 |
| yciF | sp\|P21362\|YCIF_ECOLI | 4.316219533 | 0.000228764 |
| ais | sp\|P45565\|AIS_ECOLI | 4.363027397 | 0.02438333 |
| kefG | sp\|P0A756\|KEFG_ECOLI | 4.489497738 | 0.01318169 |
| ydcS | sp\|P76108\|YDCS_ECOLI | 4.581943058 | 0.009304996 |
| phnB | sp\|P16681\|PHNB_ECOLI | 4.606620186 | 4.21E-11 |
| sodC | sp\|P0AGD1\|SODC_ECOLI | 4.625310091 | 1.37E-05 |
| **arnA** | sp\|P77398\|ARNA_ECOLI | 4.698344251 | 1.36E-11 |
| fic | sp\|P20605\|FIC_ECOLI | 4.74532692 | 0.001647533 |
| ugpB | sp\|P0AG80\|UGPB_ECOLI | 4.789402823 | 1.62E-50 |
| phoU | sp\|P0A9K7\|PHOU_ECOLI | 4.829226009 | 7.34E-29 |
| kdpC | sp\|P03961\|KDPC_ECOLI | 4.927327983 | 0.001190624 |
| pstB | sp\|P0AAH0\|PSTB_ECOLI | 5.294764534 | 6.41E-25 |
| yebE | sp\|P33218\|YEBE_ECOLI | 5.774676565 | 1.18E-14 |
| phoA | sp\|P00634\|PPB_ECOLI | 5.790096911 | 7.32E-85 |
| phoB | sp\|P0AFJ5\|PHOB_ECOLI | Inf | 8.96E-35 |
| **arnB** | sp\|P77690\|ARNB_ECOLI | Inf | 8.99E-08 |
| appA | sp\|P07102\|PPA_ECOLI | Inf | 7.69E-05 |
| yedP | sp\|P76329\|MPGP_ECOLI | Inf | 7.72E-05 |
| ugpC | sp\|P10907\|UGPC_ECOLI | Inf | 0.000683993 |
| **arnC** | sp\|P77757\|ARNC_ECOLI | Inf | 0.008862011 |
| yfcG | sp\|P77526\|YFCG_ECOLI | Inf | 0.018153582 |
| yraR | sp\|P45469\|YRAR_ECOLI | Inf | 0.012911411 |
| yhhA | sp\|P0ADX7\|YHHA_ECOLI | Inf | 0.018724338 |
| phnH | sp\|P16686\|PHNH_ECOLI | Inf | 0.025826417 |
| gspJ | sp\|P45761\|GSPJ_ECOLI | Inf | 0.036463755 |
| **PmrA** | sp\|P30843\|PMRA_ECOLI | Inf | 0.037433165 |
| alpA | sp\|P33997\|ALPA_ECOLI | Inf | 0.037433165 |

**Table S3.** Minimum inhibitory concentrations (MICs) of strains in MOPS minimal medium (1.32mM Pi) and low Pi MOPS minimal medium (0.132mM Pi)

| **Strain** | **PMXB (ug/ml)** |  |
|---|---|---|
|  | 1.32mM Pi | 0.132mM Pi |
| MG1655 | 1 | 1 |
| Δ*pmrB* | 1 | 1 |
| Δ*pmrB*::*pmrB* | 1 | 1 |
| Δ*arnB* | 1 | 1 |
| Δ*arnB*::*arnB* | 1 | 1 |

**Table S4.**
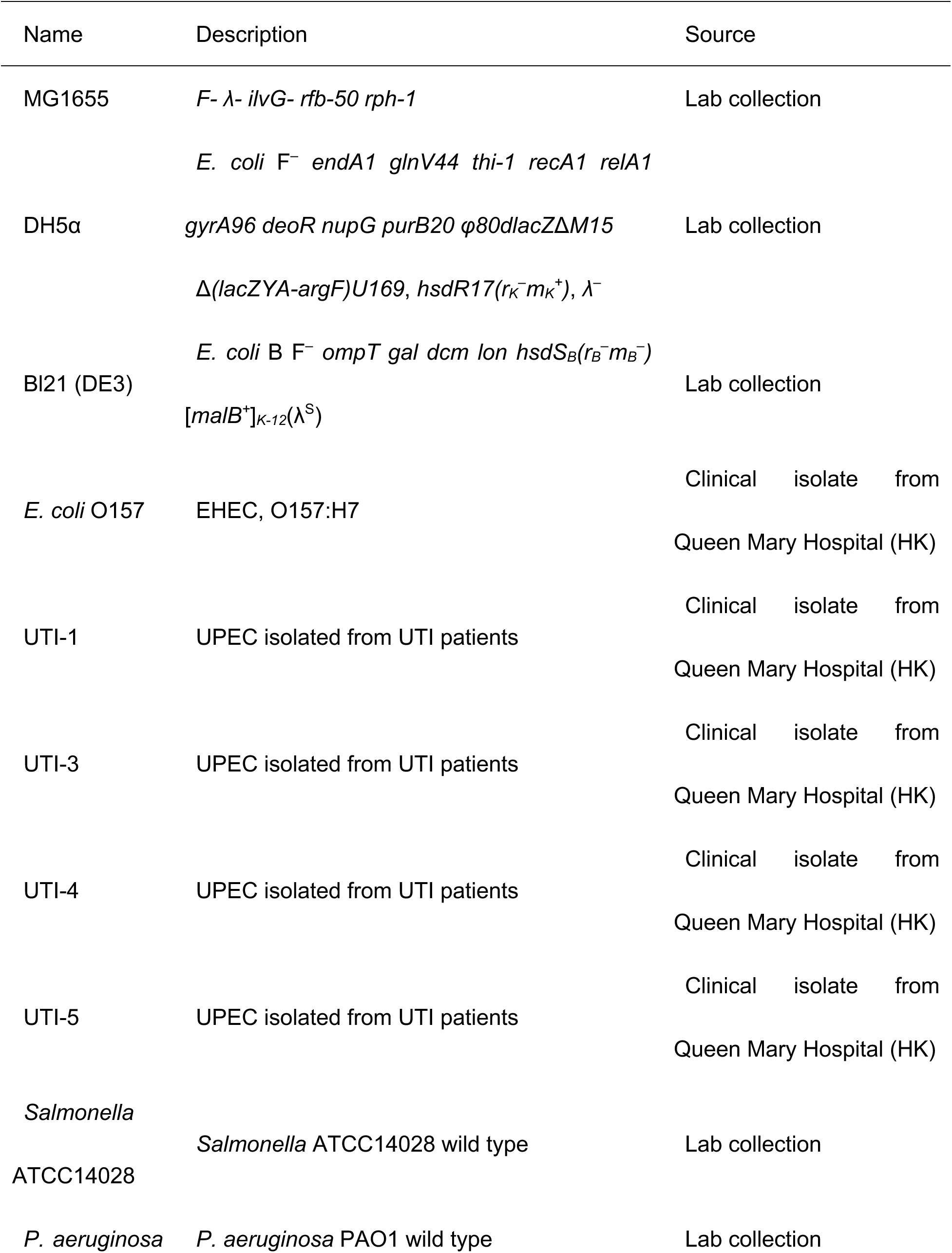
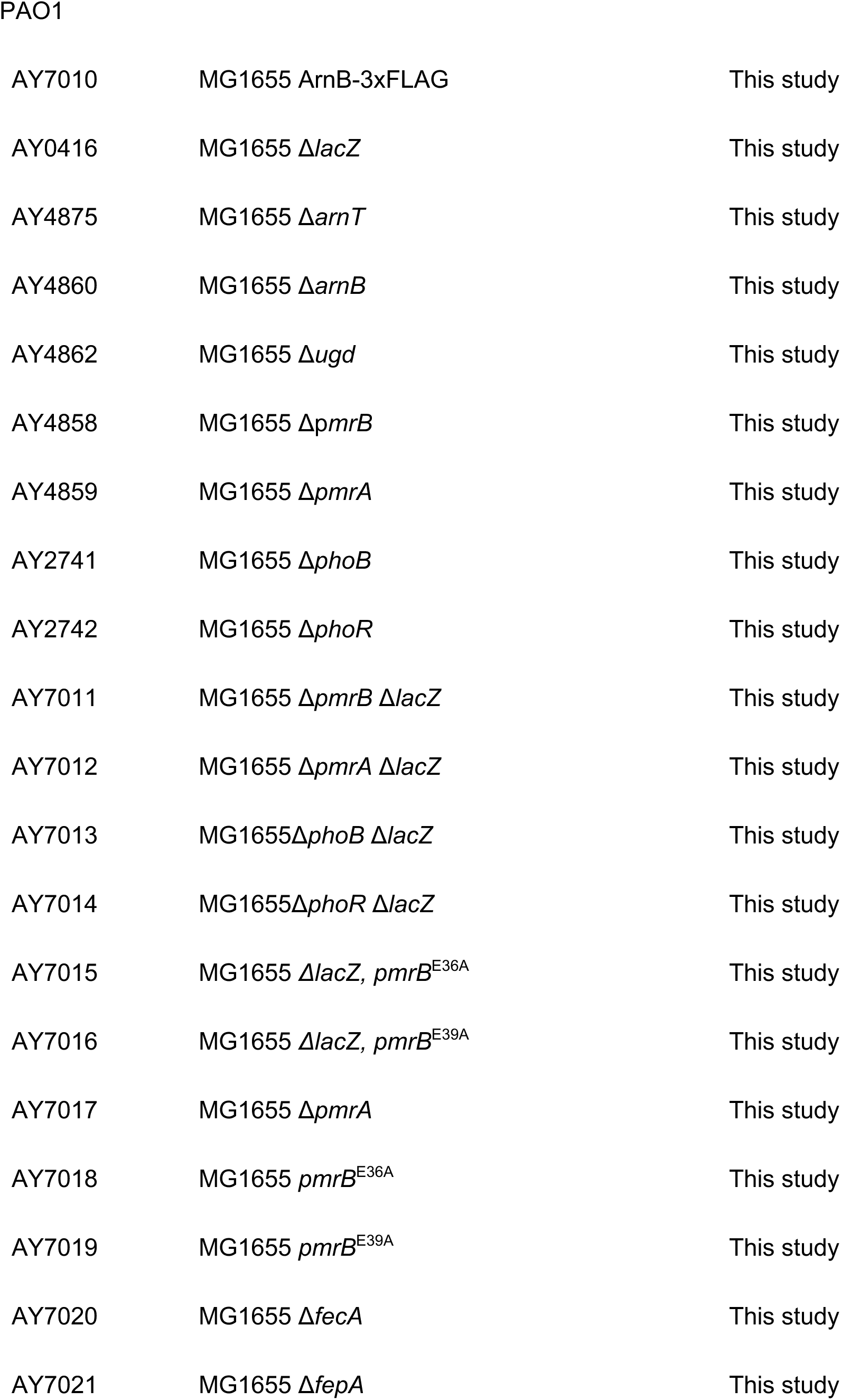
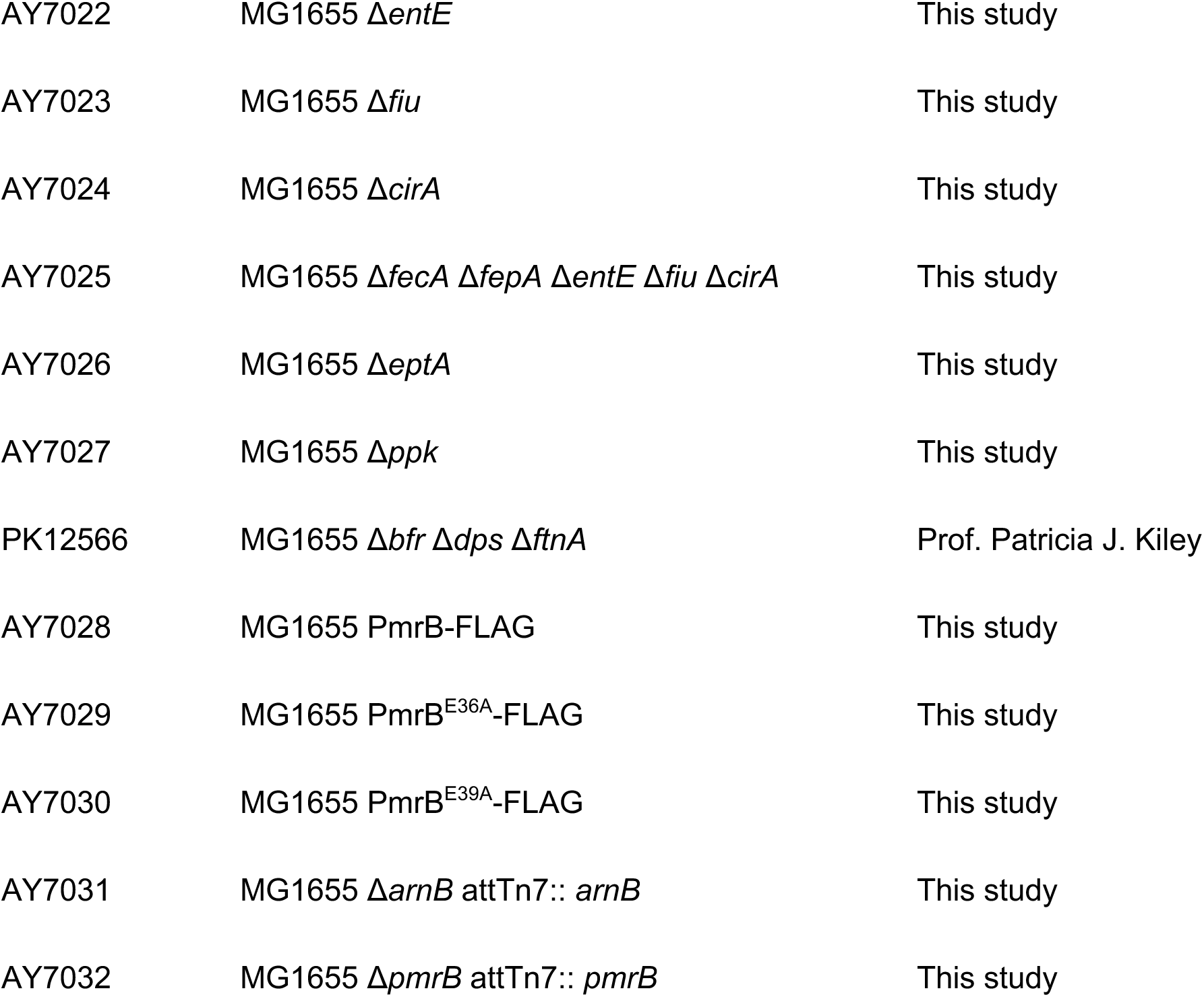
Bacteria strains used in this study.

| Name | Description | Source |
|---|---|---|
| MG1655 | *F- λ- ilvG- rfb-50 rph-1* | Lab collection |
| DH5α | *E. coli* F<sup>–</sup> *endA1 glnV44 thi-1 recA1 relA1 gyrA96 deoR nupG purB20 φ80dlacZΔM15 Δ(lacZYA-argF)U169*, *hsdR17(r<sub>K</sub><sup>–</sup>m<sub>K</sub><sup>+</sup>)*, *λ<sup>–</sup>* | Lab collection |
| Bl21 (DE3) | *E. coli* B F<sup>–</sup> *ompT gal dcm lon hsdS<sub>B</sub>(r<sub>B</sub><sup>–</sup>m<sub>B</sub><sup>–</sup>)* [*malB*<sup>+</sup>]<sub>*K-12*</sub>(λ<sup>S</sup>) | Lab collection |
| *E. coli* O157 | EHEC, O157:H7 | Clinical isolate from Queen Mary Hospital (HK) |
| UTI-1 | UPEC isolated from UTI patients | Clinical isolate from Queen Mary Hospital (HK) |
| UTI-3 | UPEC isolated from UTI patients | Clinical isolate from Queen Mary Hospital (HK) |
| UTI-4 | UPEC isolated from UTI patients | Clinical isolate from Queen Mary Hospital (HK) |
| UTI-5 | UPEC isolated from UTI patients | Clinical isolate from Queen Mary Hospital (HK) |
| *Salmonella* ATCC14028 | *Salmonella* ATCC14028 wild type | Lab collection |
| *P. aeruginosa* PAO1 | *P. aeruginosa* PAO1 wild type | Lab collection |
| AY7010 | MG1655 ArnB-3xFLAG | This study |
| AY0416 | MG1655 Δ*lacZ* | This study |
| AY4875 | MG1655 Δ*arnT* | This study |
| AY4860 | MG1655 Δ*arnB* | This study |
| AY4862 | MG1655 Δ*ugd* | This study |
| AY4858 | MG1655 Δp*mrB* | This study |
| AY4859 | MG1655 Δ*pmrA* | This study |
| AY2741 | MG1655 Δ*phoB* | This study |
| AY2742 | MG1655 Δ*phoR* | This study |
| AY7011 | MG1655 Δ*pmrB* Δ*lacZ* | This study |
| AY7012 | MG1655 Δ*pmrA* Δ*lacZ* | This study |
| AY7013 | MG1655Δ*phoB* Δ*lacZ* | This study |
| AY7014 | MG1655Δ*phoR* Δ*lacZ* | This study |
| AY7015 | MG1655 *ΔlacZ, pmrB*<sup>E36A</sup> | This study |
| AY7016 | MG1655 *ΔlacZ, pmrB*<sup>E39A</sup> | This study |
| AY7017 | MG1655 Δ*pmrA* | This study |
| AY7018 | MG1655 *pmrB*<sup>E36A</sup> | This study |
| AY7019 | MG1655 *pmrB*<sup>E39A</sup> | This study |
| AY7020 | MG1655 Δ*fecA* | This study |
| AY7021 | MG1655 Δ*fepA* | This study |
| AY7022 | MG1655 Δ*entE* | This study |
| AY7023 | MG1655 Δ*fiu* | This study |
| AY7024 | MG1655 Δ*cirA* | This study |
| AY7025 | MG1655 Δ*fecA* Δ*fepA* Δ*entE* Δ*fiu* Δ*cirA* | This study |
| AY7026 | MG1655 Δ*eptA* | This study |
| AY7027 | MG1655 Δ*ppk* | This study |
| PK12566 | MG1655 Δ*bfr* Δ*dps* Δ*ftnA* | Prof. Patricia J. Kiley |
| AY7028 | MG1655 PmrB-FLAG | This study |
| AY7029 | MG1655 PmrB<sup>E36A</sup>-FLAG | This study |
| AY7030 | MG1655 PmrB<sup>E39A</sup>-FLAG | This study |
| AY7031 | MG1655 Δ*arnB* attTn7:: *arnB* | This study |
| AY7032 | MG1655 Δ*pmrB* attTn7:: *pmrB* | This study |

**Table S5.** Plasmids used in this study.

| Name | Relevant characteristics | Source |
|---|---|---|
| pNN387 | Single copy plasmid containing NotI and HindIII restriction sites before *lacZ* | Lab collection |
| pNN387(P<sub>*arn*</sub>-*lacZ*) | Ligate promoter of *arn* into NotI/HindIII linearized pNN387 | This study |
| pNN387(P<sub>*ugd*</sub>-*lacZ*) | Ligate promoter of *ugd* into NotI/HindIII linearized pNN387 | This study |
| pNN387(P<sub>*phoQ*</sub>-*lacZ*) | Ligate promoter of *phoQ* into NotI/HindIII linearized pNN387 | This study |
| pNN387(P<sub>*mgtA*</sub>-*lacZ*) | Ligate promoter of *mgtA* into NotI/HindIII linearized pNN387 | This study |
| pET28a-N | t7 promoter in pET-28a is replaced by araBad promoter | This study |
| pET28a-N-3xFLAG | C-terminal 6xHis tag in pET-28a-N is replaced by 3xFLAG tag | This study |
| pET-N-PmrA-6xHis | Ligate *pmrA* into NcoI/XhoI linearized pET-28a-N | This study |
| pET-N-PmrA-3xFLAG | Ligate *pmrA* into NcoI/XhoI linearized pET-28a-N-3xFLAG | This study |
| pET-N-PmrA137TAG-6xHis | Mutate G137 to TAG in pET28A-N-PmrA-6xHis | This study |
| pEVOL-CouA-RS | Plasmid expression CouA tRNA synthetase | Lab collection |
| pCAGO | Editing plasmid containing CRISPR-Cas9 and λ-red | Gift from Prof. Changhao Bi |

